# From Transcriptomic Signatures to Active-Site Docking and ADMET Properties Prediction: Bioinformatics-Guided Exploration of Binding Mechanisms of Cardioprotective Phytoconstituents Against Dysregulated Targets in Hypertrophic Cardiomyopathy

**DOI:** 10.1101/2024.10.13.618070

**Authors:** Basavaraj Vastrad, Shivaling Pattanashetti, Chanabasayya Vastrad

## Abstract

Hypertrophic cardiomyopathy (HCM) is a global health problem characterized by left ventricle become thick and stiff with effect of indication including chest pain, fluttering, fainting and shortness of breath. In this investigation, we aimed to identify diagnostic markers and analyzed the therapeutic potential of essential genes. Next generation sequencing (NGS) dataset GSE180313 was obtained from the Gene Expression Omnibus (GEO) database and used to identify differentially expressed genes (DEGs) in HCM. DEGs were screened using the DESeq2 Rbioconductor tool. Then, Gene Ontology (GO) and REACTOME pathway enrichment analyses were performed. Moreover, a protein-protein interaction (PPI) network of the DEGs was constructed, and module analysis was performed. Next, miRNA-hub gene regulatory network, TF-hub gene regulatory network and drug-hub gene interaction network were constructed and analyzed. Next, diagnostic values of hub genes were assessed by receiver operating characteristic (ROC) curve analysis. Finely, molecular docking and ADMET studies were performed. By performing DEGs analysis, total 958 DEGs ( 479 up regulated genes and 479 down regulated genes) were successfully identified from GSE180313, respectively. GO and REACTOME pathway enrichment analyses revealed that GO functions and signaling pathways were significantly enriched including response to stimulus, multicellular organismal process, metabolism and extracellular matrix organization. The hub genes of FN1, SOX2, TUBA4A, RPS2, TUBA1C, ESR1, SNCA, LCK, PAK1 and APLNR might be associated with HCM. The hub gens of FN1 and TPM3, together with corresponding predicted miRNAs (e.g., hsa-mir-374a-5p and hsa-miR-8052), SH3KBP1 and ESR1 together with corresponding predicted TFs (e.g PRRX2 and STAT3) and CHRNA4 and CHRM3 together with corresponding predicted drug molecules (e.g. Succinylcholine and Umeclidinium) were found to be significantly correlated with HCM. Molecular Docking study revealed that all tested phytoconstituents effectively occupied LGALS3 and ESR1. ADMET analysis found that all screened compounds have adequate solubility and high projected intestine absorption, indicating their appropriateness for oral administration, This investigation could serve as a basis for further understanding the molecular pathogenesis and potential therapeutic targets of HCM.

## Introduction

Hypertrophic cardiomyopathy (HCM) is a prevalent heritable cardiovascular disease [Argulian et al. 2016], which affects around 1:500 (0.2%) of the world’s population [Massera et al. 2023]. HCM is primarily characterized by left ventricle become thick and stiff [Teekakirikul et al. 2019]. The clinical incidence of HCM is high, and its main features include chest pain, fluttering, fainting and shortness of breath [Varma et al. 2014]. Studies have revealed that the progression of HCM is related to genetic factors [Pérez-Sánchez et al. 2018]. Complications of HCM can include atrial fibrillation [Rowin et al. 2023], blocked blood flow [Seiler et al. 1991], mitral valve disease [Griffeth et al. 2024], dilated cardiomyopathy [Puckelwartz et al. 2021], heart failure [Liang et al. 2023], cardiac death [Moore et al. 2019], myocardial infarction [Ariss et al. 2022], myocardial ischemia [Cecchi et al. 2009], coronary artery disease [Tower-Rader Cecchi et al. 2017], cardiac arrest [Gupta et al. 2016], atherosclerosis [Maron et al. 1979], cardiac fibrosis [Xu et al. 2021], hypercholesterolemia [Marziliano et al. 2021], inflammation [Becker et al. 2020], hypertension [Arabadjian et al. 2023], obesity [Zaromytidou and Savvatis, 2023] and diabetes mellitus [Subramanian et al. 2023]. Medications (beta blockers, calcium channel blockers, mavacamten, amiodarone and Warfarin) [Wheeler et al. 2023; Bright et al. 1991; Nag et al. 2023; Boriani et al. 2002; Noseworthy et al. 2016] and Surgeries or other procedures (septal myectomy, septal ablation, implantable cardioverter-defibrillator, cardiac resynchronization therapy, ventricular assist device and heart transplant) [Cui et al. 2022; Maron et al. 2021; Radu et al. 2023; Sreenivasan et al. 2021; Loyaga-Rendon et al. 2021] are the current feasible treatment of HCM patients. Despite advancement in diagnosis and treatment, the prognosis of HCM patient’s remains destitute, this has become an effective topic of clinical and fundamental investigation. Hence, dynamic novel diagnosis biomarkers thoughtful the damage of heart muscles is engaging for the timely diagnoses and therapies of HCM.

The recent next generation sequencing (NGS) analysis of specimens from sufferers and normal individuals enables us to investigate various diseases [Ganekal et al. 2023; Giriyappagoudar et al. 2023]. Genetic mutations, epigenetic alterations and aberrant molecular signaling pathways are associated in the HCM [Seidman and Seidman, 2011; Greco and Condorelli, 2015; Hassoun et al. 2021]. Recently, numerous specific genes have been identified to participate in the advancement of HCM. For instance, it was reported that the expression level of ACTN2 [Chiu et al. 2010], MYBPC3 [Tudurachi et al. 2023], MYOZ2 [Osio et al. 2007], ACTC [Mogensen et al. 1999] and NEXN [Wang et al. 2010] were considerably higher in damaged heart muscles from HCM sufferers in contrast to normal heart muscles. RAS signaling pathway [Gelb and Tartaglia, 2011], Hippo/YAP signaling pathway [Wang et al. 2014], Akt/GSK-3β signaling pathway [Luckey et al. 2009], alcineurin-NFAT signaling pathway [Braz et al. 2003] and mTOR signaling pathway [Schramm et al. 2012] reveals that the treatment manipulation of such signaling pathways can offer novel enlightenment for the treatment of HCM. These finding suggested the key roles of few function genes in HCM development. However, the diagnostic value of many genes has not been studied in HCM.

NGS technology and bioinformatics analysis have been broadly applied to screen for genetic modification at the genomic level. Our main determination is to explore the connection between HCM and its complications. First, we download the GSE180313 [Ranjbarvaziri et al. 2021] dataset file in the NCBI-Gene Expression Omnibus database (NCBI-GEO) (https://www.ncbi.nlm.nih.gov/geo) [Clough and Barrett, 2016] database for analysis, then use DESeq2 [Love et al. 2014] to draw the differentially expressed genes (DEGs) distribution map of the HCM and normal control samples in the dataset. Immediately after, we performed gene ontology (GO) and REACTOME pathway enrichment analysis on these DEGs. By constructing protein-protein interaction (PPI) network, we screened for the highest scoring modules and hub genes. Immediately afterward miRNA-hub gene regulatory network, TF-hub gene regulatory network and drug-hub gene interaction network were built by Cytoscape to predict the underlying microRNAs (miRNAs), transcription factors (TFs) and drug molecules linked with hub genes. To validate that these hub genes can serve as biomarkers of HCM, we determined diagnostic value of hub gene by receiver operating characteristic (ROC) curve analysis. molecular docking and ADMET studies for phytoconstituents were carried out. This investigation will develop our perceptive of the molecular mechanisms of HCM and supply genomic-targeted therapy preference for HCM.

## Materials and Methods

### Next generation sequencing (NGS) data source

The publicly accessible NGS data were all obtained from the NCBI-GEO. NGS dataset deposited by Ranjbarvaziri et al. (2021), accession no. GSE180313, containing 20 samples (13 HCM samples and 7 normal control samples) were employed in the current investigation. The GSE180313 dataset generated using the GPL24676 platform (Illumina NovaSeq 6000 (Homo sapiens)).

### Identification of DEGs

Determining DEGs between HCM and normal blood samples was performed using DESeq2 [Love et al. 2014]. DESeq2 is an R bioconductor tool that allows users to determine DEGs for various experimental situations. The adjusted P-values (adj. P) and Benjamini and Hochberg false discovery rate were used to find statistically significant genes [Green and Diggle, 2007]. The screening criteria of DEGs were set as false discovery rate (FDR) ≤ 0.05, and |log2 fold change (FC)| > 0.828 for up regulated genes and |log2 fold change (FC)| < -1.219 for down regulated genes. The volcano plot and heatmap were generated using the “ggplot2” and “gplot” packages in the R bioconductor software.

### GO and pathway enrichment analyses of DEGs

Online biological information can be establish in the g:Profiler (http://biit.cs.ut.ee/gprofiler/) [Reimand et al. 2007] program. A essential bioinformatics tool for annotating genes and examining their biological processes is the GO (http://www.geneontology.org) [Thomas, 2017]. GO enrichment analysis includes biological processes (BP), cellular components (CC), and molecular functions (MF). The REACTOME pathway (https://reactome.org/) [Fabregat et al. 2018] is a database tool for analyzing large-scale molecular data sets produced by high-throughput experimental techniques to superior understand high-level biological processes and systems. P < 0.05 was considered statistically significant.

### Construction of the PPI network and module analysis

The Cytoscape software (version 3.10.2) (http://www.cytoscape.org/) [Shannon et al. 2003] and HIPPIE (Human Integrated Protein-Protein Interaction rEference) (https://cbdm-01.zdv.uni-mainz.de/~mschaefer/hippie/) [Schaefer et al. 2013] interactome database were used to establish the PPI analysis of the DEGs. First, the HIPPIE PPI database was used to construct the PPI network. The PPI network was visualized using Cytoscape (version 3.10.2). Subsequently, genes with the top interaction scores were screened as hub genes using the node degree [Luo et al. 2017], betweenness [Li et al. 2017], stress [Gilbert et al. 2021] and closeness [Li et al. 2020] algorithms of the Network Analyzer plugin in Cytoscape software. And PEWCC algorithm [Zaki et al 2013] was used for identifying significant modules.

### Construction of the miRNA-hub gene regulatory network

miRNet (https://www.mirnet.ca/) [Fan et al 2018] is a bioinformatics platform for predicting miRNAs. In the current investigation, the targets of the hub gene were predicted using eight databases: TarBase, miRTarBase, miRecords, miRanda, miR2Disease, HMDD, PhenomiR, SM2miR, PharmacomiR, EpimiR, starBase, TransmiR, ADmiRE, and TAM 2.0. The miRNA-hub gene regulatory network was depicted and visualized using Cytoscape software [Shannon et al. 2003].

### Construction of the TF-hub gene regulatory network

NetworkAnalyst (https://www.networkanalyst.ca/) [Zhou et al 2019] is a bioinformatics platform for predicting TFs. In the current investigation, the targets of the hub gene were predicted using Jasper database. The TF-hub gene regulatory network was depicted and visualized using Cytoscape software [Shannon et al. 2003].

### Construction of the drug-hub gene interaction network

NetworkAnalyst (https://www.networkanalyst.ca/) [Zhou et al 2019] is a bioinformatics platform for predicting drug molecules. In the current investigation, the targets of the hub gene were predicted using DrhuBank database. The drug-hub gene interaction network was depicted and visualized using Cytoscape software [Shannon et al. 2003].

### Receiver operating characteristic curve (ROC) analysis

To determine the sensitivity and specificity of the selected hub genes for HCM diagnosis, the visualization tool “pROC” package in R software [Robin et al. 2011] was used to construct a receiver operating characteristic (ROC) curve. The area under the curve (AUC) was calculated for each dataset, and hub genes with an AUC exceptional 0.7 were classified as holding a strong inequitable capacity for HCM.

### *Insilico* molecular docking studies

Swiss-model, RCSB PDB, Prank web, NCBI Gene, ChEMBL, BindingDB, DoGSiteScorer, ChemDraw, Avogadro tool, Autodock 1.7.1, and Autodock Vina tools, Biovia Discovery Studio client 2021, ADMET lab 3.0 web server.

### Receptor selection based on gene information

The selection of target receptor structuresfor molecular docking was initiated by identifying genes of interest relevant to the disease condition under study. Genes showing differential expression or known functional involvement in disease pathophysiology were prioritized based on genomic and transcriptomic data [Barabási et al., 2011; Subramanian et al., 2005]. The corresponding protein products of these genes were then retrieved using UniProt (https://www.uniprot.org/) and NCBI Gene (https://www.ncbi.nlm.nih.gov/gene/) databases. These databases provide curated information on gene-protein relationships, functional domains, isoforms, and organism-specific variants [UniProt 2023].

Once the protein names and UniProt accessions were determined, three-dimensional structures of these proteins were searched in the Protein Data Bank (PDB) (https://www.rcsb.org/). Priority was given to experimentally determined structures (X-ray crystallography or cryo-EM) derived from Homo sapiens, with resolutions ≤2.5 Å and co-crystallised ligands when available. Structures were evaluated for completeness, presence of functional domains, and biologically relevant binding conformations. In cases where multiple structures were available, the one with the most complete and biologically relevant ligand-protein interaction site was selected [Berman et al., 2000; Plewczynski et al., 2011].

If co-crystallised ligands were present in the selected PDB entry, these were examined to confirm their biological relevance (e.g., substrate, inhibitor, agonist). The functional nature of the ligand was cross-validated using ligand bioactivity databases such as ChEMBL (https://www.ebi.ac.uk/chembl/) and BindingDB (https://www.bindingdb.org/), where half-maximal inhibitory concentration (IC₅₀), binding affinity (Kᵢ), or other pharmacological data are reported [Gaulton et al., 2017; Liu et al., 2007]. To confirm the presence and accessibility of druggable binding pockets, cavity detection tools such as CASTp, Fpocket, or DoGSiteScorer were used [Tian et al., 2018; Le et al., 2009].

This ensured that the selected structure was suitable for molecular docking, with a validated and accessible ligand-binding domain. Overall, this systematic approach, starting from gene selection to receptor structure identification, ensured biological relevance, structural accuracy, and docking compatibility of the chosen targets. In the present study, we have selected two genes, upregulated LGALS3 and downregulated ESR1 in present study.

The LGALS3 gene that produces galectin-3, a β-galactoside-binding lectin that plays a role in hypertrophic cardiomyopathy by causing myocardial fibrosis, inflammation, and unfavorableremodeling [Emet et al., 2018; Yakar et al., 2016]. Elevated galectin-3 levels in HCM patients correlate with increased ventricular mass and fibrosis, as well as risk factors such as sudden cardiac death and arrhythmias, indicating its clinical significance in HCM development and prognosis.Galectin-3 is increased in stressed or failing myocardium, promoting macrophage activation, fibroblast proliferation, and collagen deposition. These processes are directly associated with abnormal cardiac remodeling, which drives diastolic dysfunction and disease progression in HCM [Sharma et al., 2004].

Estrogen receptor α (ESR1) inhibits cardiomyocyte hypertrophy, reduces inflammation, and regulates cellular stress responses. Estrogen-ERα signaling inhibits cardiomyocyte hypertrophy and pathological remodeling via genomic and non-genomic mechanisms [Luo et al., 2016].Although direct large-scale studies of ESR1 expression in HCM are rare, mutations in the ESR1 gene have been related with variation in left ventricular hypertrophy among HCM patients, indicating a role in disease phenotypic change [Lind et al., 2008].Furthermore, experimental evidence suggests that estrogen can decrease left ventricular and cardiomyocyte hypertrophy via receptor-mediated pathways, bolstering the rationale for studying ESR1 regulation in hypertrophic disorders [Ueda et al., 2020]. However, this selection does not necessarily imply that other potential targets lack significance or therapeutic relevance.

### Selection phytoconstituents

Puerarin is a natural isoflavone glycoside produced from Pueraria lobata. It’s been extensively explored for its cardioprotective and cardiac remodeling properties. Preclinical research suggests that puerarin effectively inhibits pathological hypertrophic development, decreases myocyte mortality, suppresses fibroblast proliferation, and attenuates inflammatory and oxidative stress pathways related to cardiac remodeling [Lv et al., 2022;Zhou et al., 2021].Oleuropein, a polyphenolic chemical found in olives and olive leaf extracts, is renowned for wide cardioprotective activities, including antioxidant, anti-inflammatory, and anti-remodeling actions [Christodoulou et al., 2024;Omar et al., 2010]. Swertiamarin is an iridoid glycoside that exhibits significant antioxidant and anti-inflammatory properties in heart damage models. Swertiamarin has exhibited considerable restoration of oxidative stress indicators and pro-inflammatory cytokines in experimental myocardial infarction, indicating its potential function in alleviating heart stress and remodeling [Wang et al., 2023].Mangiferin, a xanthone glucoside found in many plants, has been shown to provide myocardial protection, antifibrotic action, and anti-apoptotic properties [Imran et al., 2017;Hou et al., 2013].

Although the selected phytoconstituentsSwertiamarin, Puerarin, Oleuropein, and Mangiferinhave previously been reported to exhibit general cardioprotective, antioxidant, and anti-inflammatory activities in a variety of cardiovascular and myocardial injury models, their molecular mechanisms of action in hypertrophic cardiomyopathy remain unclear. Importantly, none of these drugs have been thoroughly tested for their structural interactions with HCM-specific, transcriptomically dysregulated targets such as the upregulated fibrotic mediator LGALS3 and the downregulated cardioprotective receptor ESR1. The current study thus employs a target-driven mechanistic repositioning strategy, in which well-characterized cardioprotective phytochemicals are re-examined for their potential to modulate disease-relevant molecular drivers of HCM, rather than being interpreted as established or disease-specific modulators in hypertrophic cardiomyopathy.

### Receptor and ligand preparation

The targets selected with their structural information are presented in Table 1. Using the Swiss-Model web server, the missing residues were remodelled and downloaded in pdb format [Waterhouse et al., 2018]. Following the identification of the binding sites utilising the server Prank web [Jendele et al., 2019],the protein’s pdb format was entered into Software Auto Dock Tools 1.7.1, and water molecules and atoms were eliminated. The receptor has also been checked for missing amino acid residues, Kollman charges have been fixed, and only polar hydrogens have been inserted. In order to cover the entire receptor, the grid was fixed using the Autogrid program. The grid dimension file [Morris et al., 2008] was then saved. The phytoconstituents (Fig. 1), structures were drawn using ChemDraw and copied as SMILES then loaded into Avogadro software to convert them 2D to 3D structures in PDB format [Hanwell et al., 2012]. The ligand’s pdb format was then loaded into Auto Dock Tools 1.7.1, and the ligand molecule’s root was identified, selected and saved as pdbqt format. Lastly, the pdbqt format was used to save the ligand molecule.

**Fig. 1.**
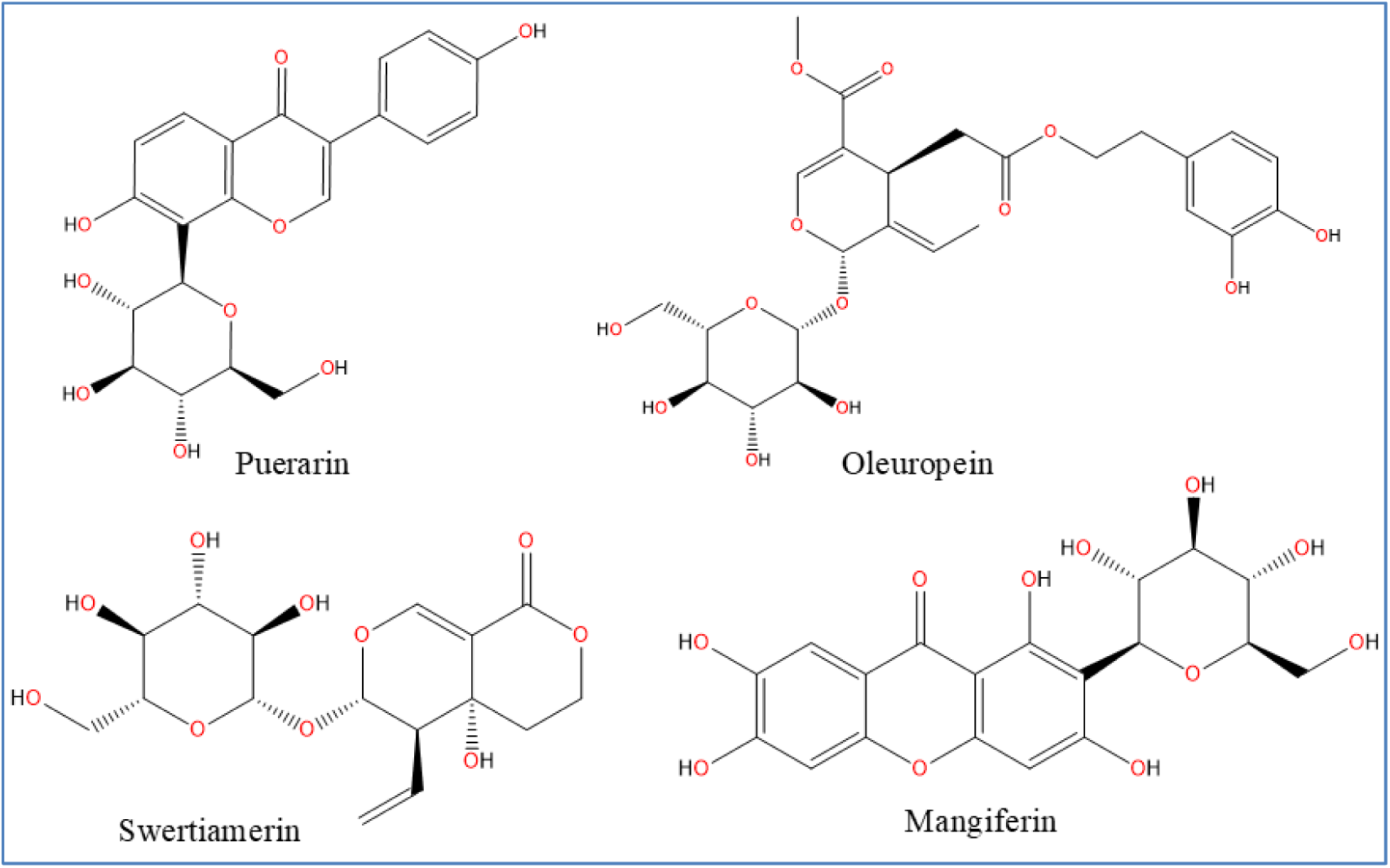
Chemical structures of the screened phytoconstituentsPuerarin, Oleuropein, Swertiamarin, and Mangiferinwere chosen for structure-based screening against hypertrophic cardiomyopathy-associated targets (LGALS3 and ESR1).

**Table 1.** Structural details of selected hypertrophic cardiomyopathyassociated targets used for molecular docking.

| Target gene | Topology of the genes in HCM | PDB ID | Resolution (Å) | Method of collection | Observed R-value | Cocrystal ligand |
| --- | --- | --- | --- | --- | --- | --- |
| LGALS3 | Up regulated | 8IU1 | 1.97 | X-ray diffraction | 0.175 | <u>QB2</u> |
| ESR1 | Down regulated | 1A52 | 2.80 | X-ray diffraction | 0.223 | <u>EST</u> |

### Performing autodockvina

Auto Dock Vina can be executed using the command line (cmd) or the Autodock tool. The configuration file was ready for Autodock Vina to execute; the grid dimension file that was previously saved includes the protein’s n-points, active site, and x, y, and z coordinates. That information was added to the configuration file, which was made to contain the protein’s active site details. It also comes with an output file in pdbqt format and a log file in .txt format. The command line was used to run Autodock Vina. “vina.exe -- config config.txt” was the command used to launch Auto Dock Vina. Docked coordinates were output in the pdbqt format when the program was finished. Receptor-ligand interactions were then visualised using the file, and binding affinity was ascertained using the log.txt file [Riyaphan et al., 2021].

### Visualization of molecular docking results

The docking results were visualized using the Biovia Discovery Studio Client 2021 application. The pdbqt files for the receptor and the output docked file were both loaded into Biovia, and a 2D docked image of the ligand was visualised and exported in PNG format, which is connected to various amino acids [James et al., 2025].

### *In silico* ADMET properties prediction

However, high binding affinity alone is insufficient. ADMET profiling, evaluating Absorption, Distribution, Metabolism, Excretion, and Toxicity, is essential to ensure that candidate compounds possess favourable pharmacokinetic properties and safety profiles. Without this, compounds may fail in later stages despite strong docking results. Integration of docking with in silico ADMET prediction streamlines compound selection by filtering out unsuitable leads early in the discovery pipeline [Xiong et al., 2021]. ADMET properties were predicted with the help of the ADMET lab 2.0 free web server, as previously reported by[Ahmad et al., 2023].

## Results

### Identification of DEGs

We selected GSE180313 NGS dataset to analyze and identify the DEGs. A total of 958 DEGs were identified, which include 479 up regulated DEGs and 479 down regulated DEGs (Table 2). The volcano plot and heatmap are shown in Fig. 2 and Fig. 3.

**Fig. 2.**
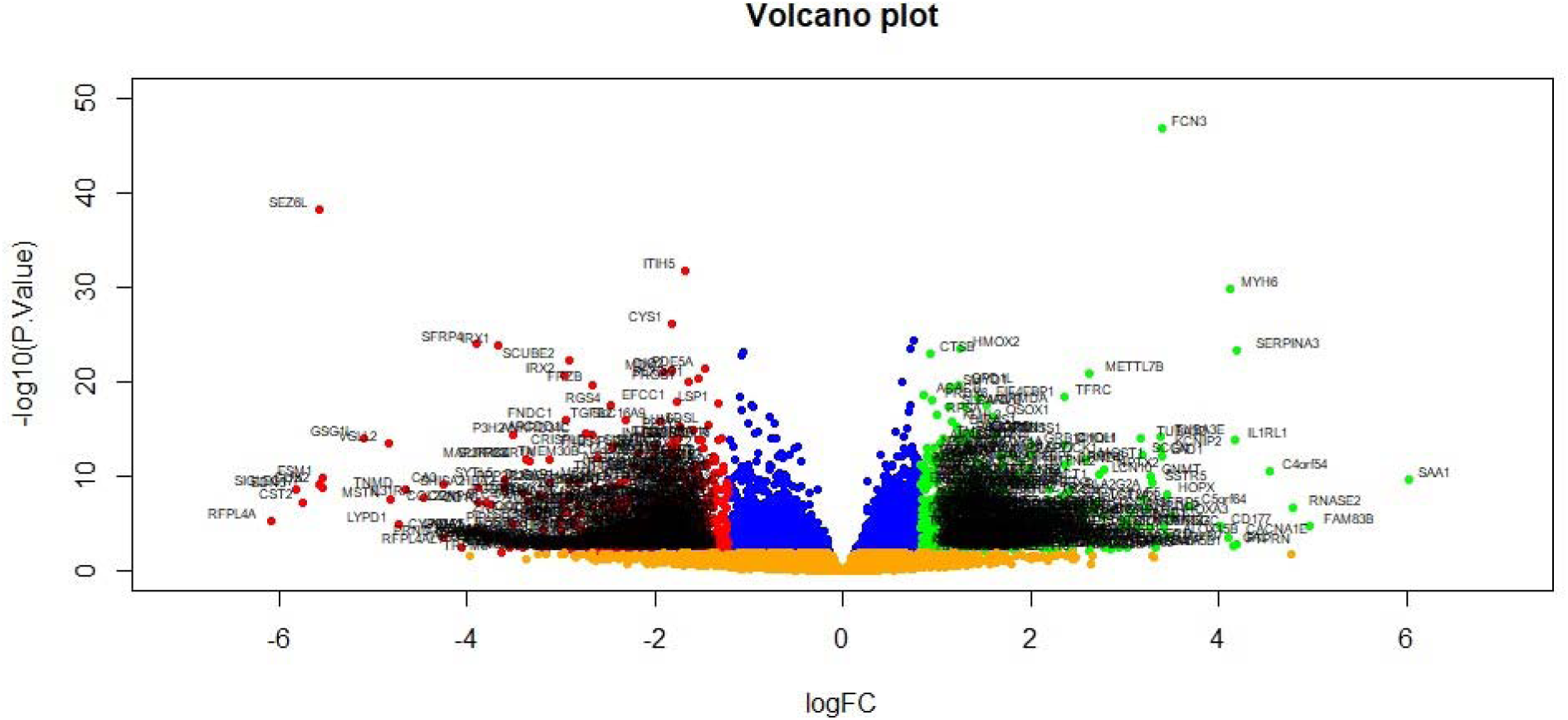
Volcano plot of differentially expressed genes. Genes with a significant change of more than two-fold were selected. Green dot represented up regulated significant genes and red dot represented down regulated significant genes.

**Fig. 3.**
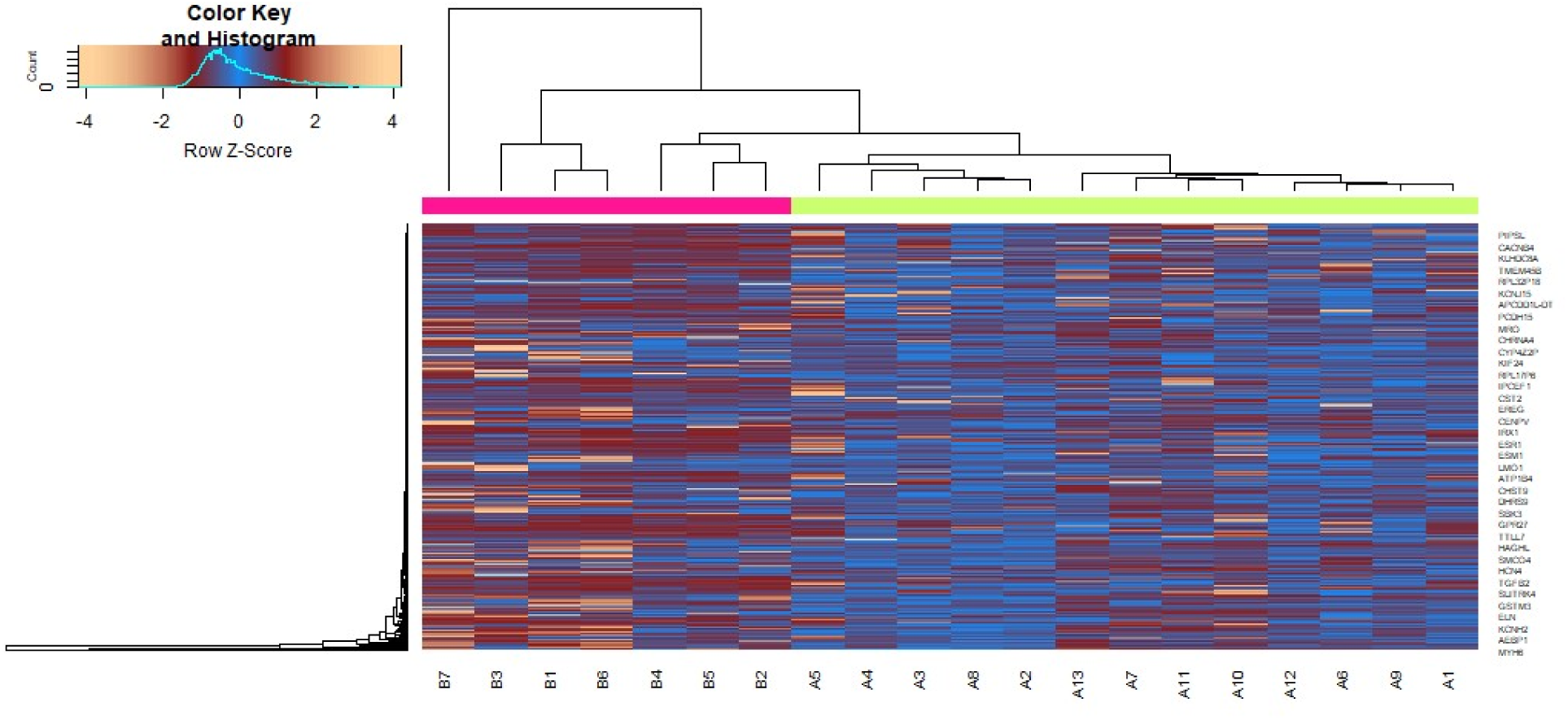
Heat map of differentially expressed genes. Legend on the top left indicate log fold change of genes. (A1 – A106 = HD samples; B1 – B46 = Normal control samples)

**Table 2.**
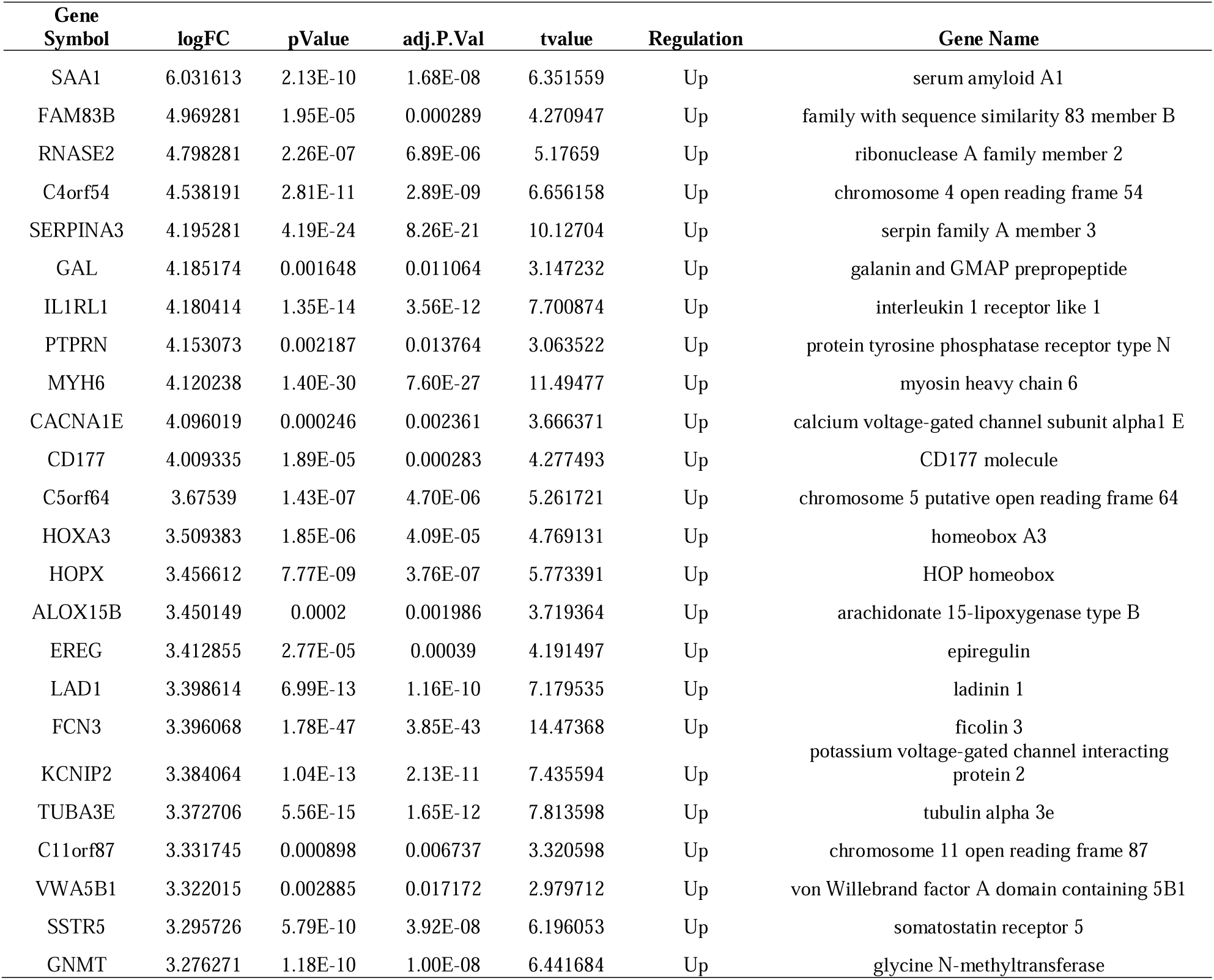
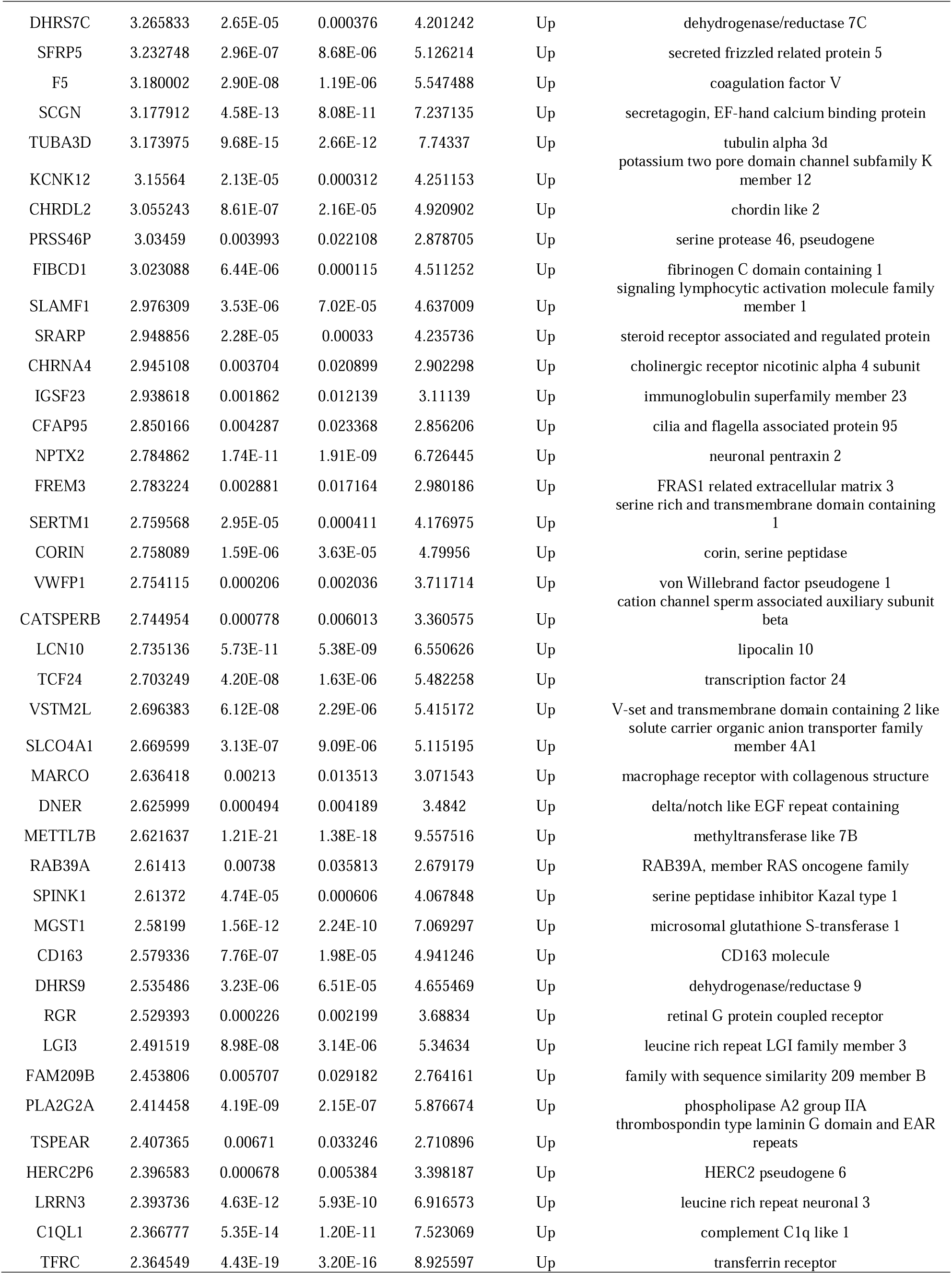
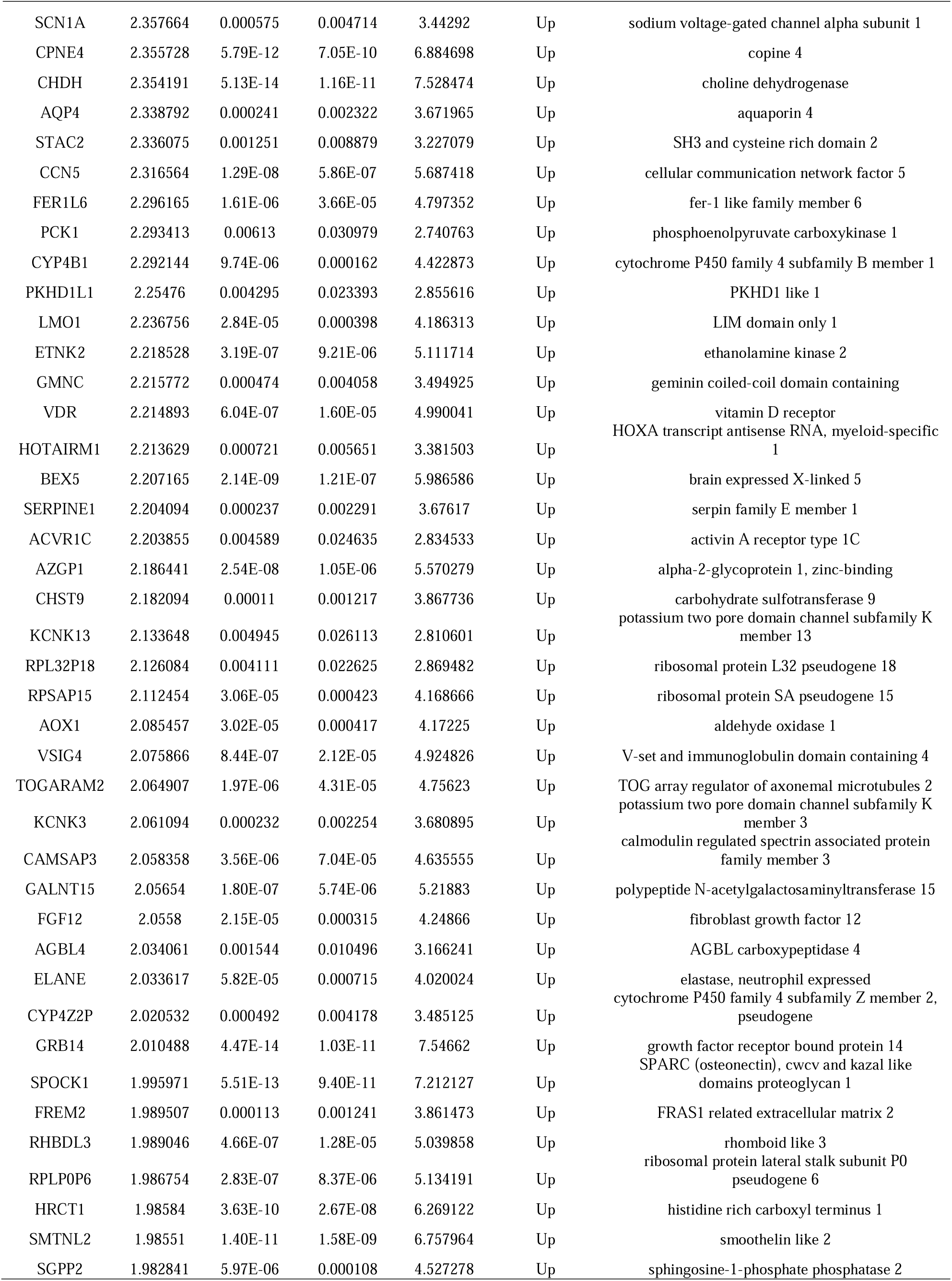
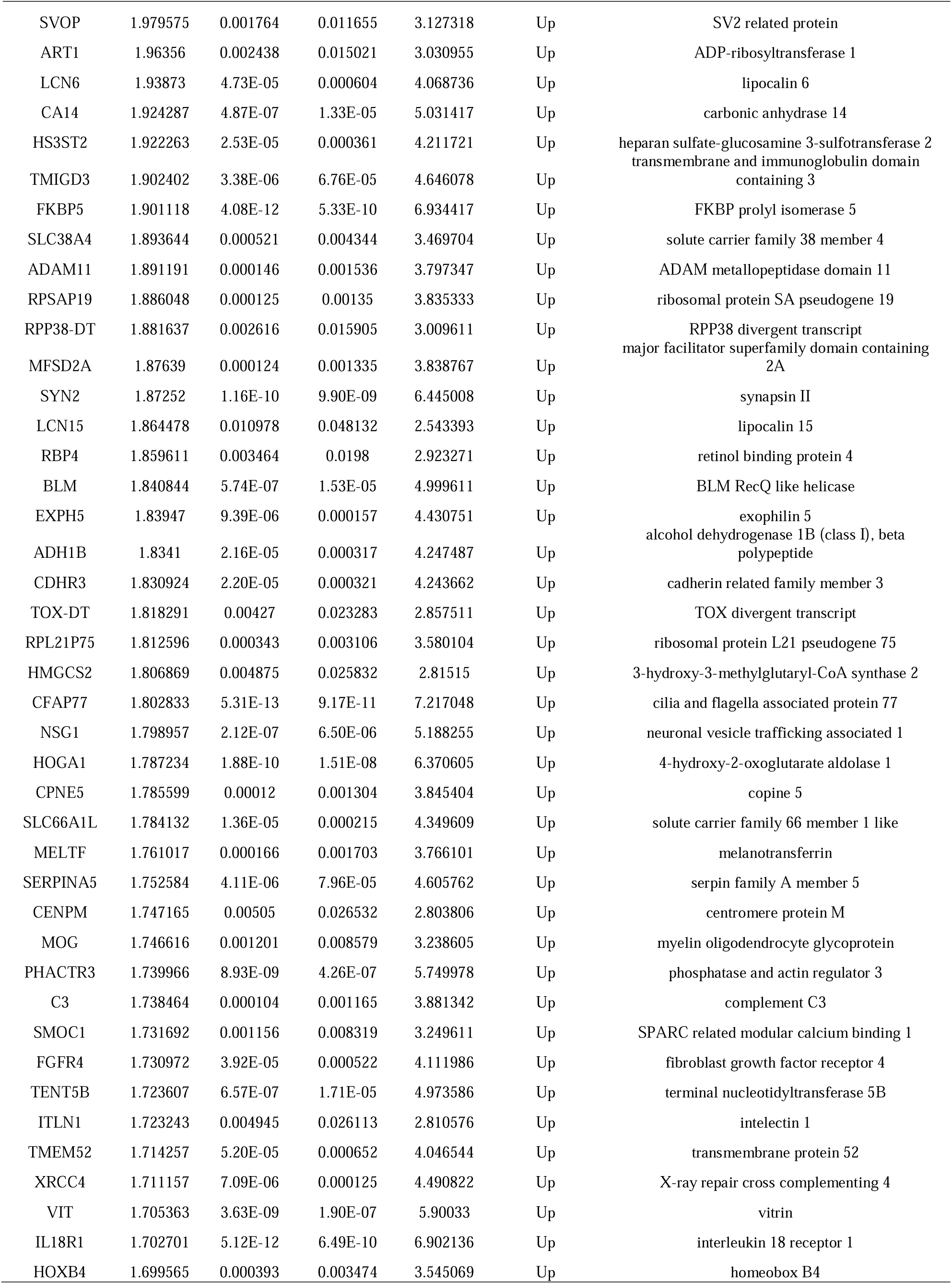
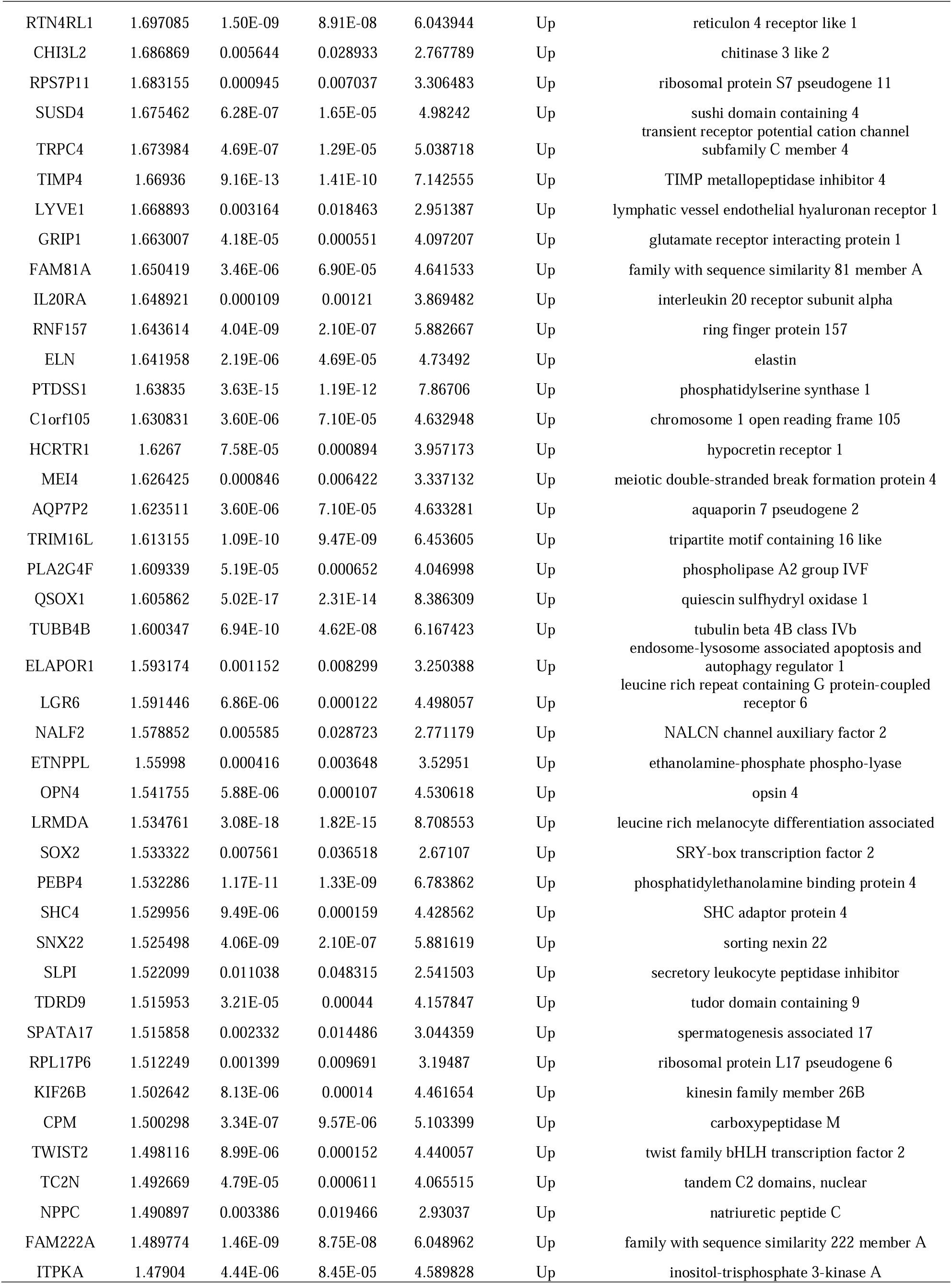
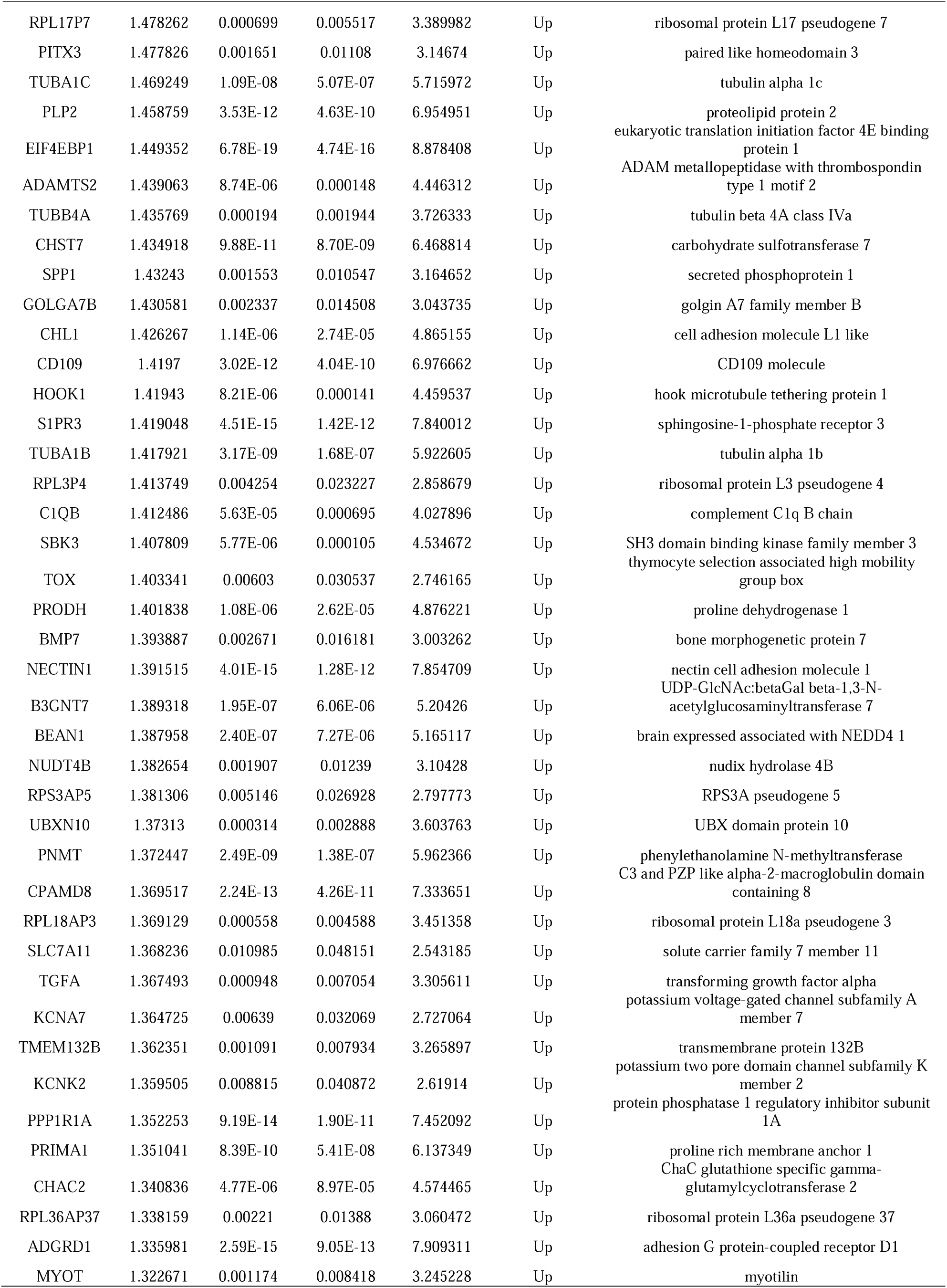
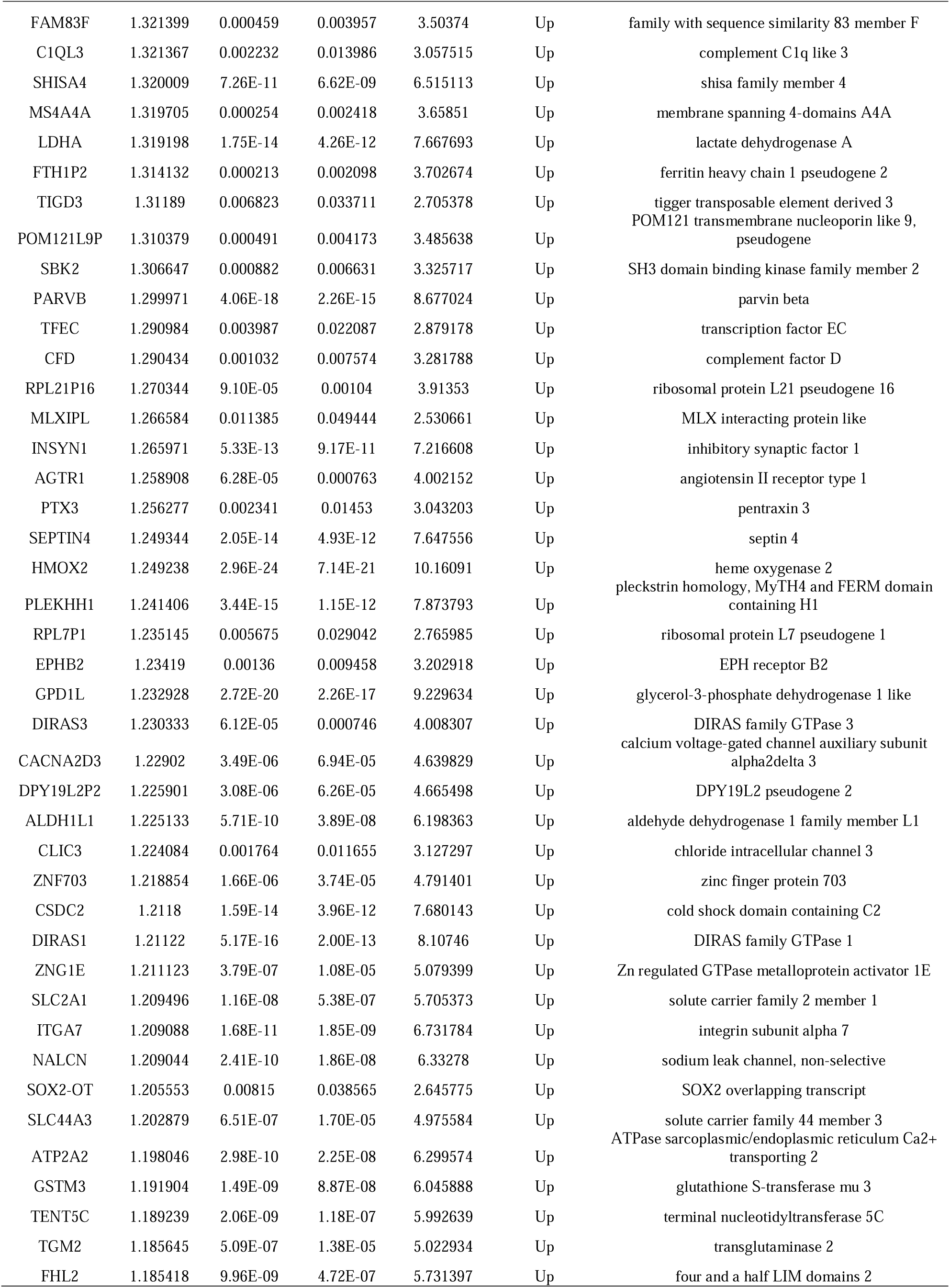
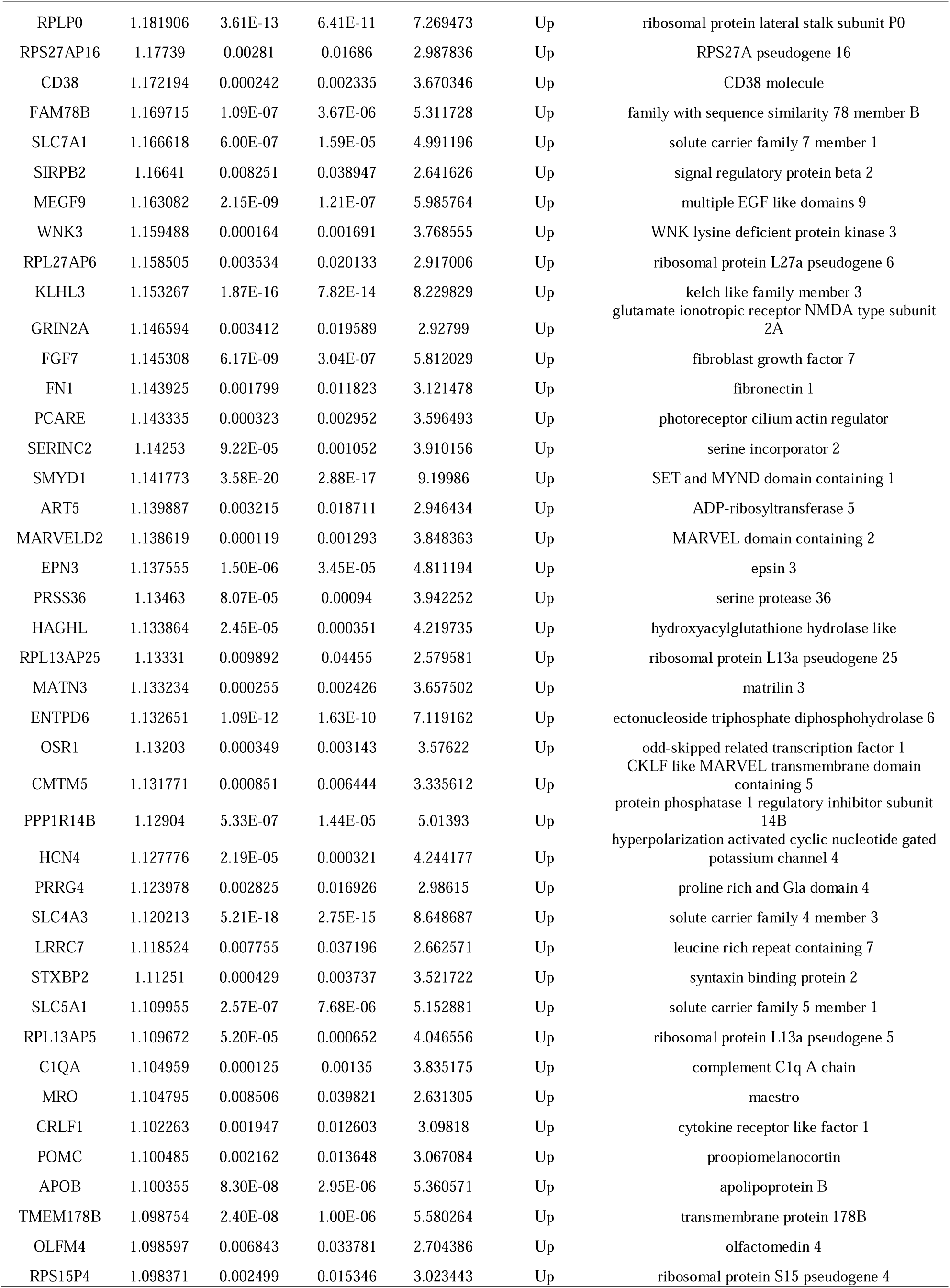
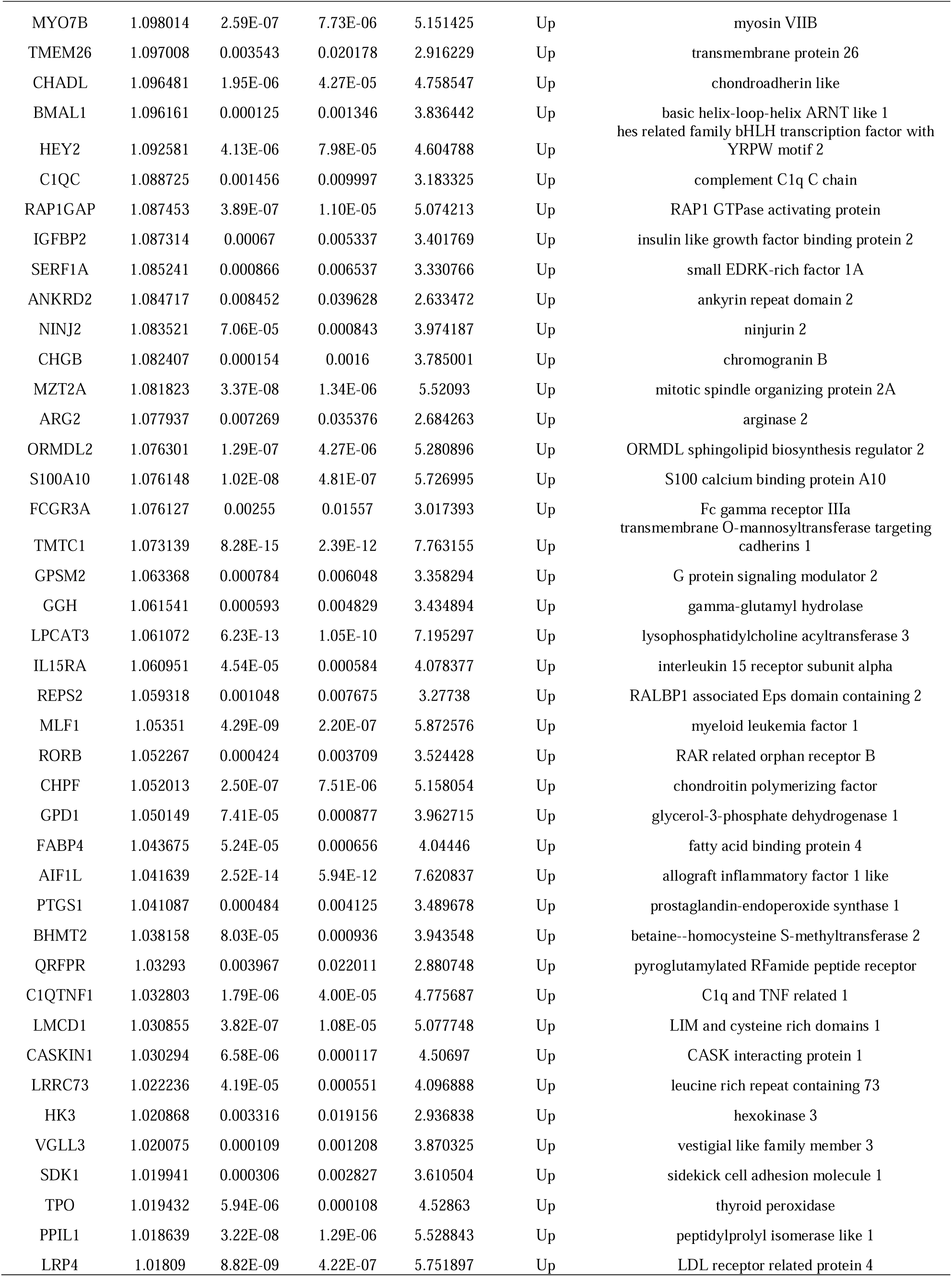
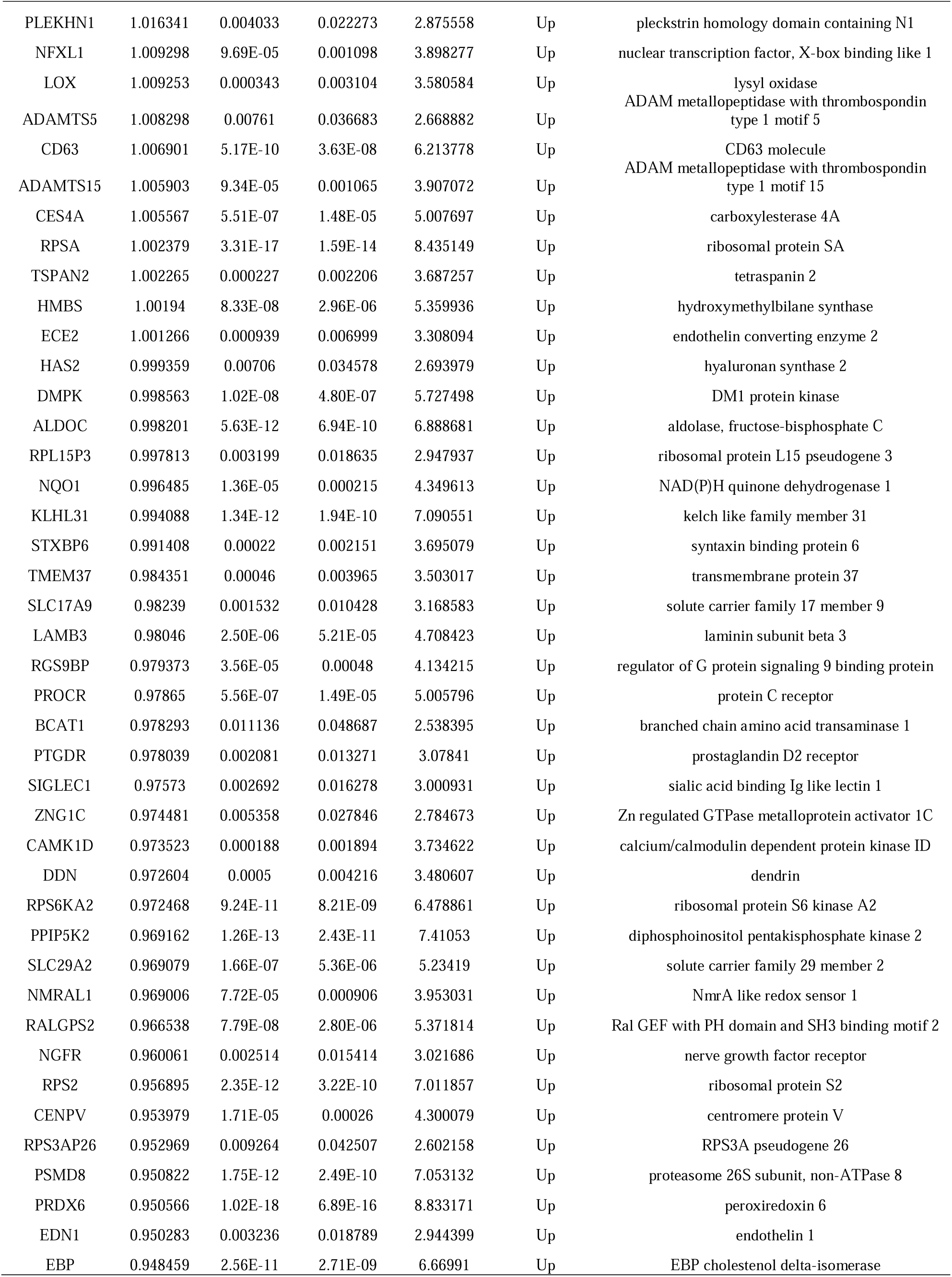
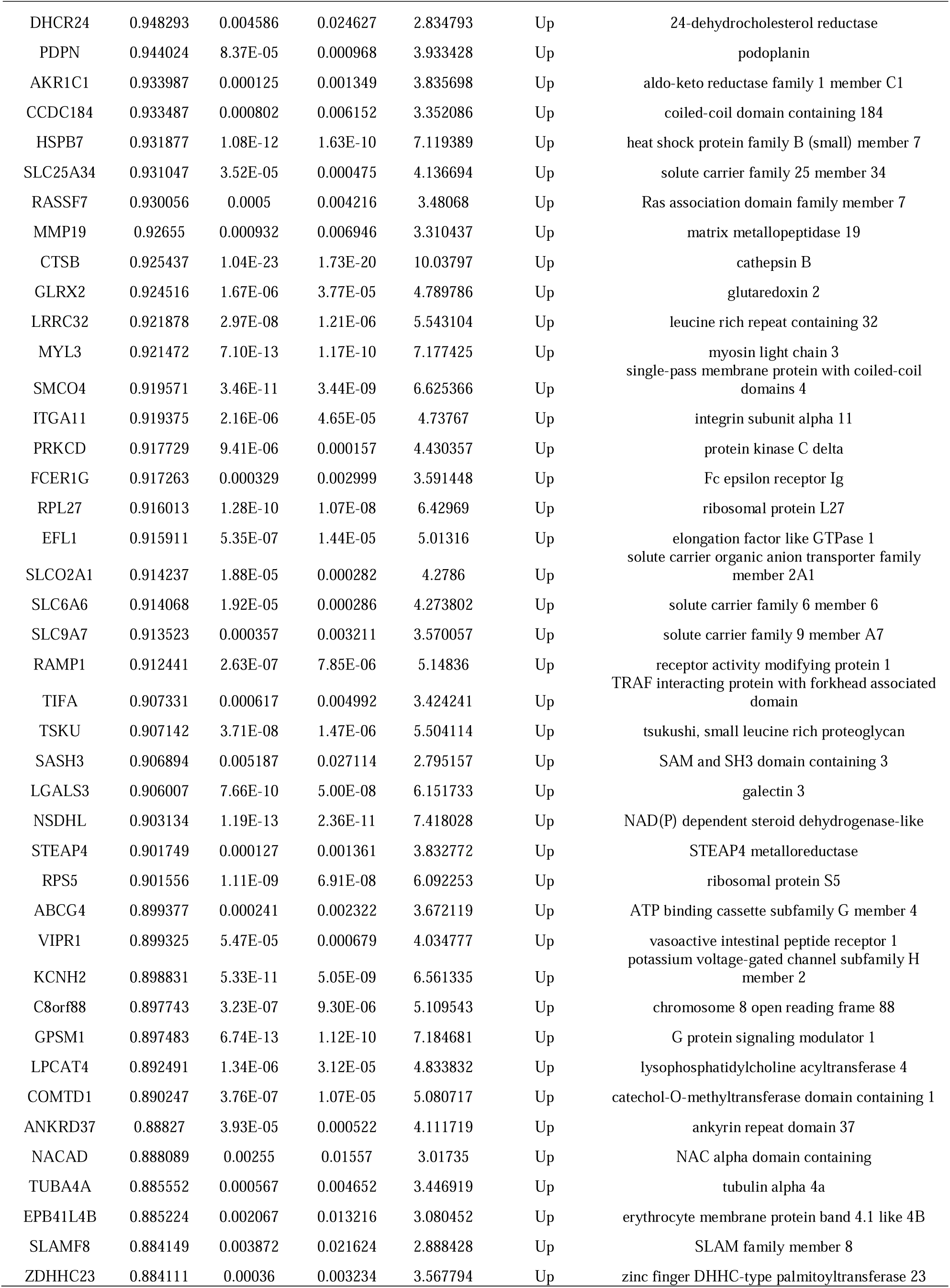
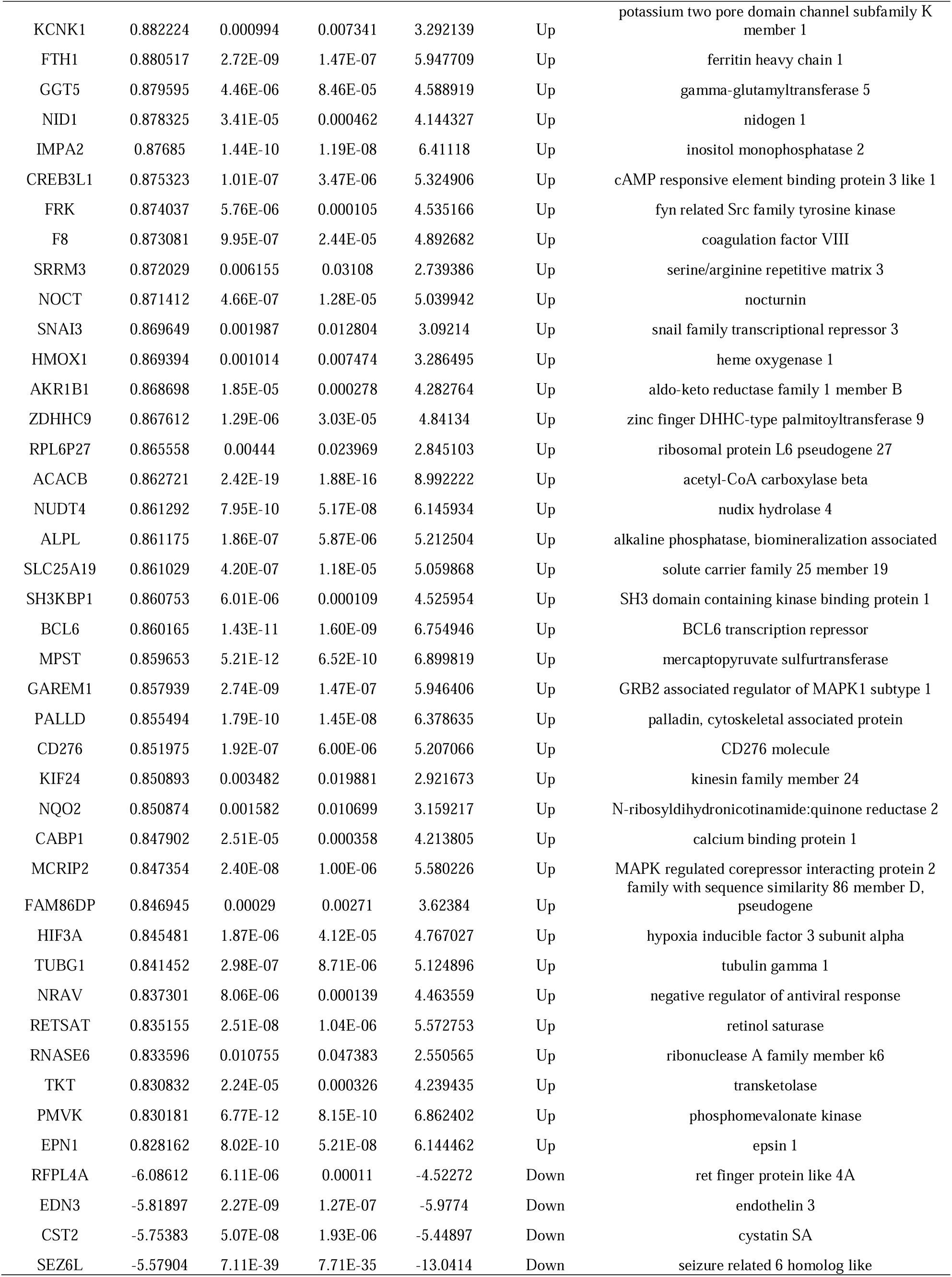
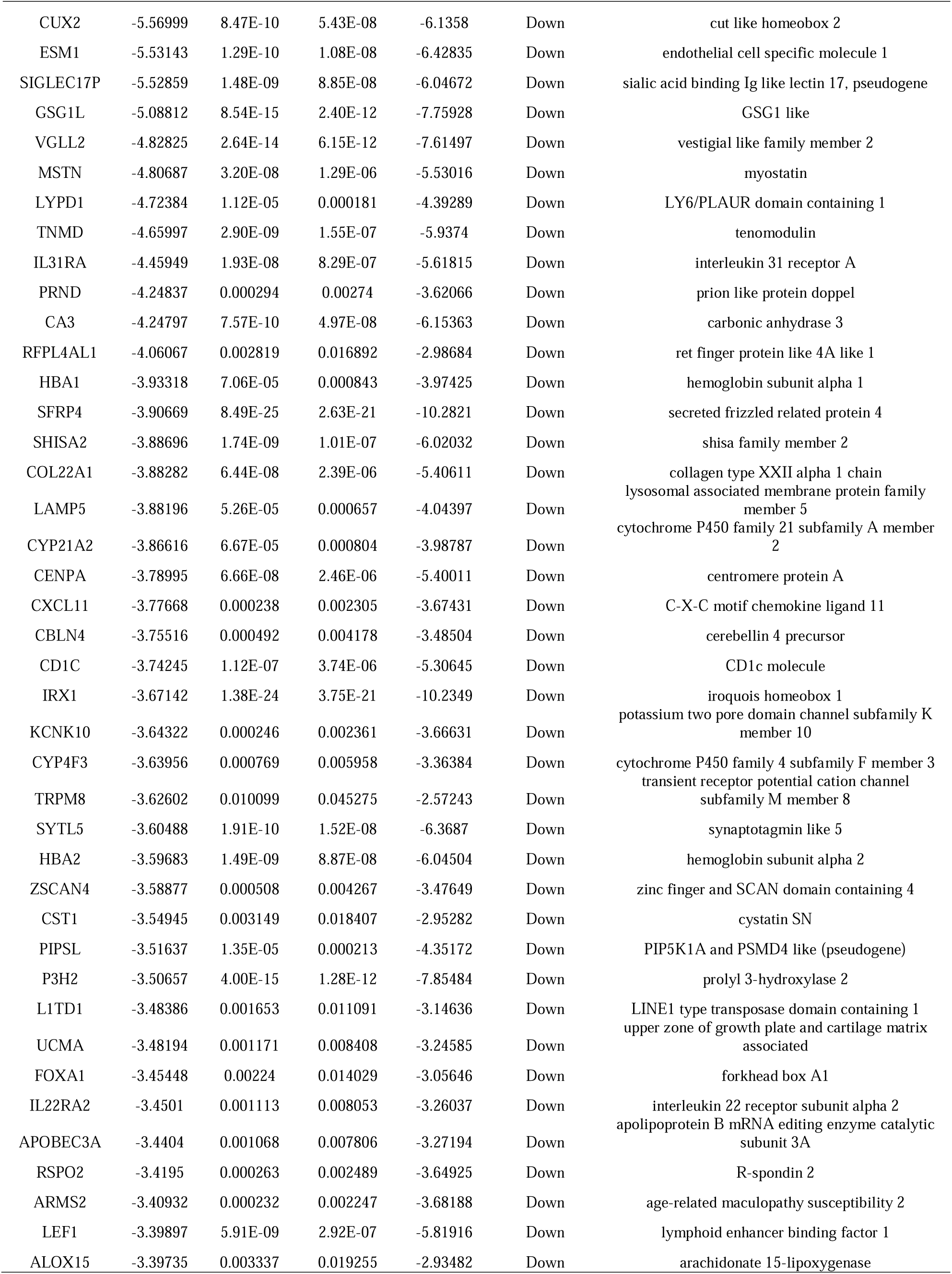
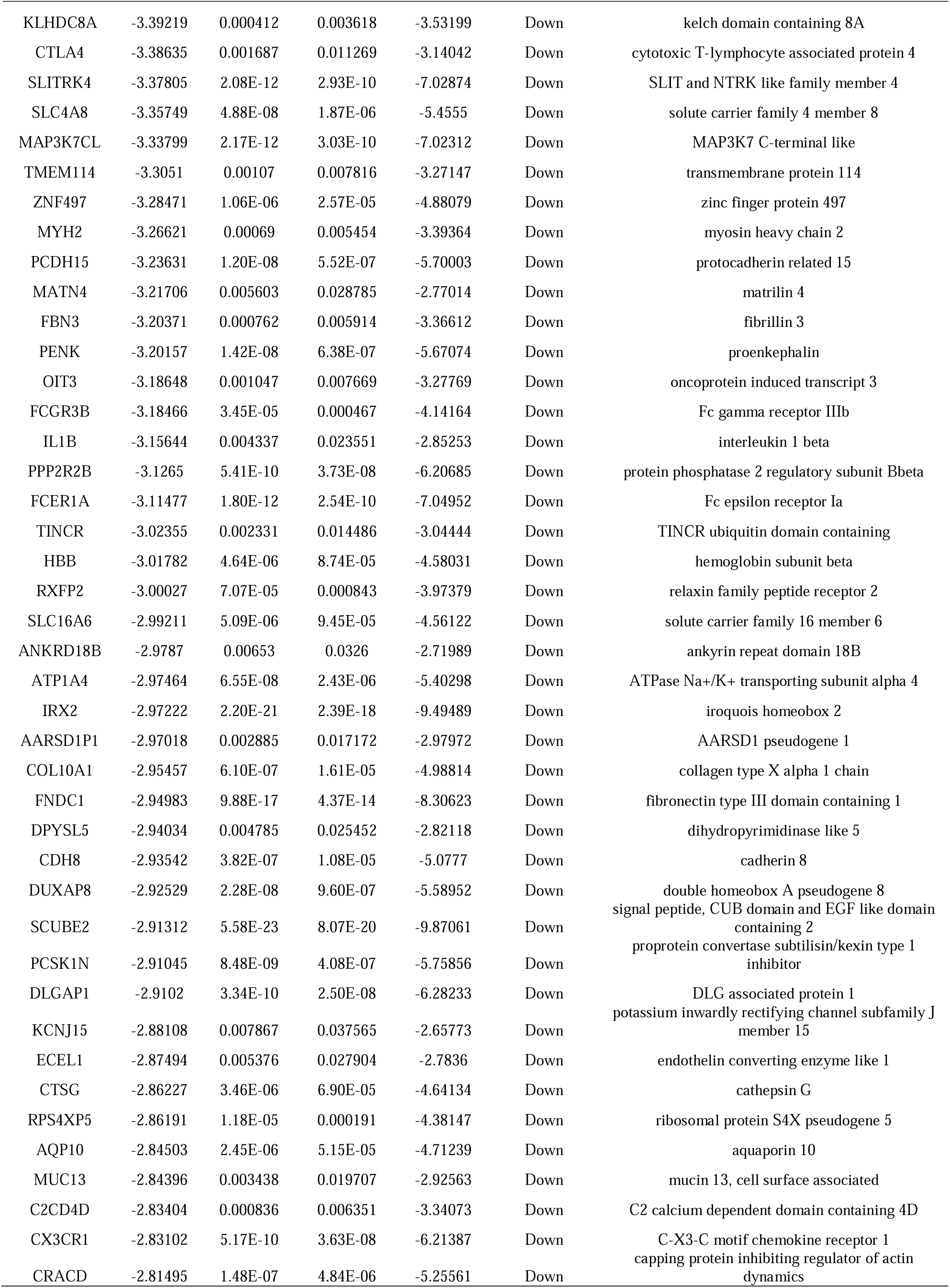
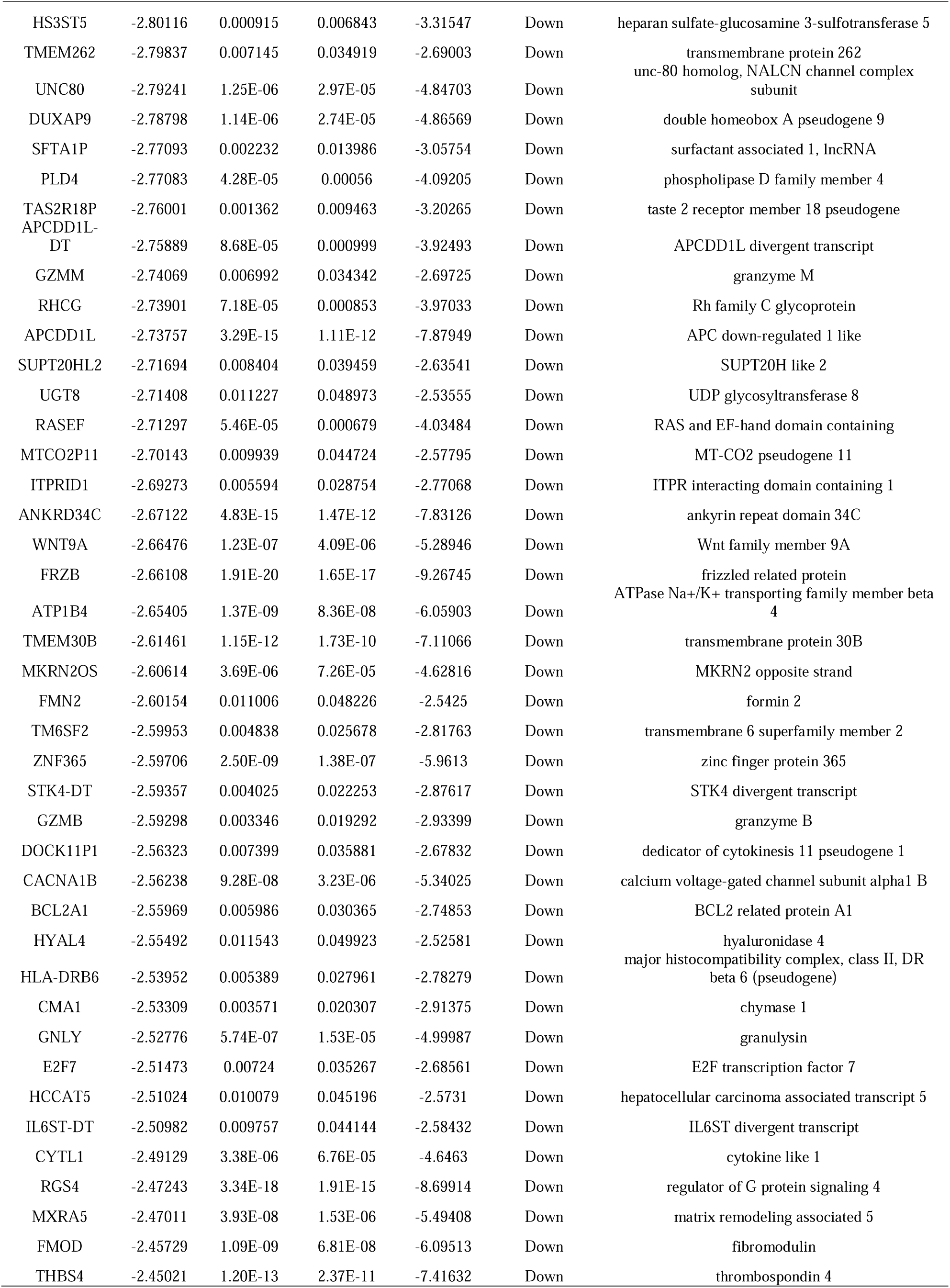
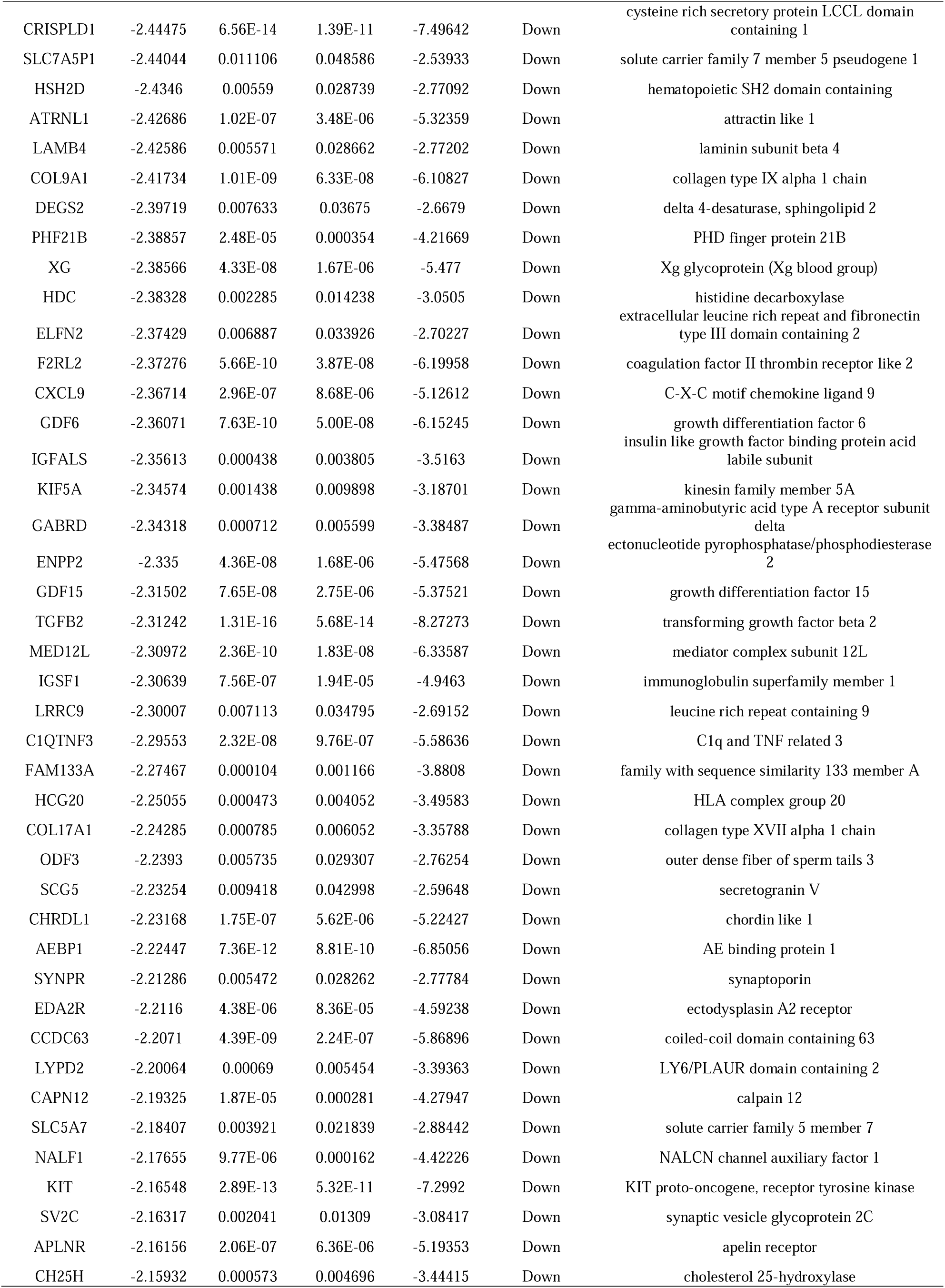
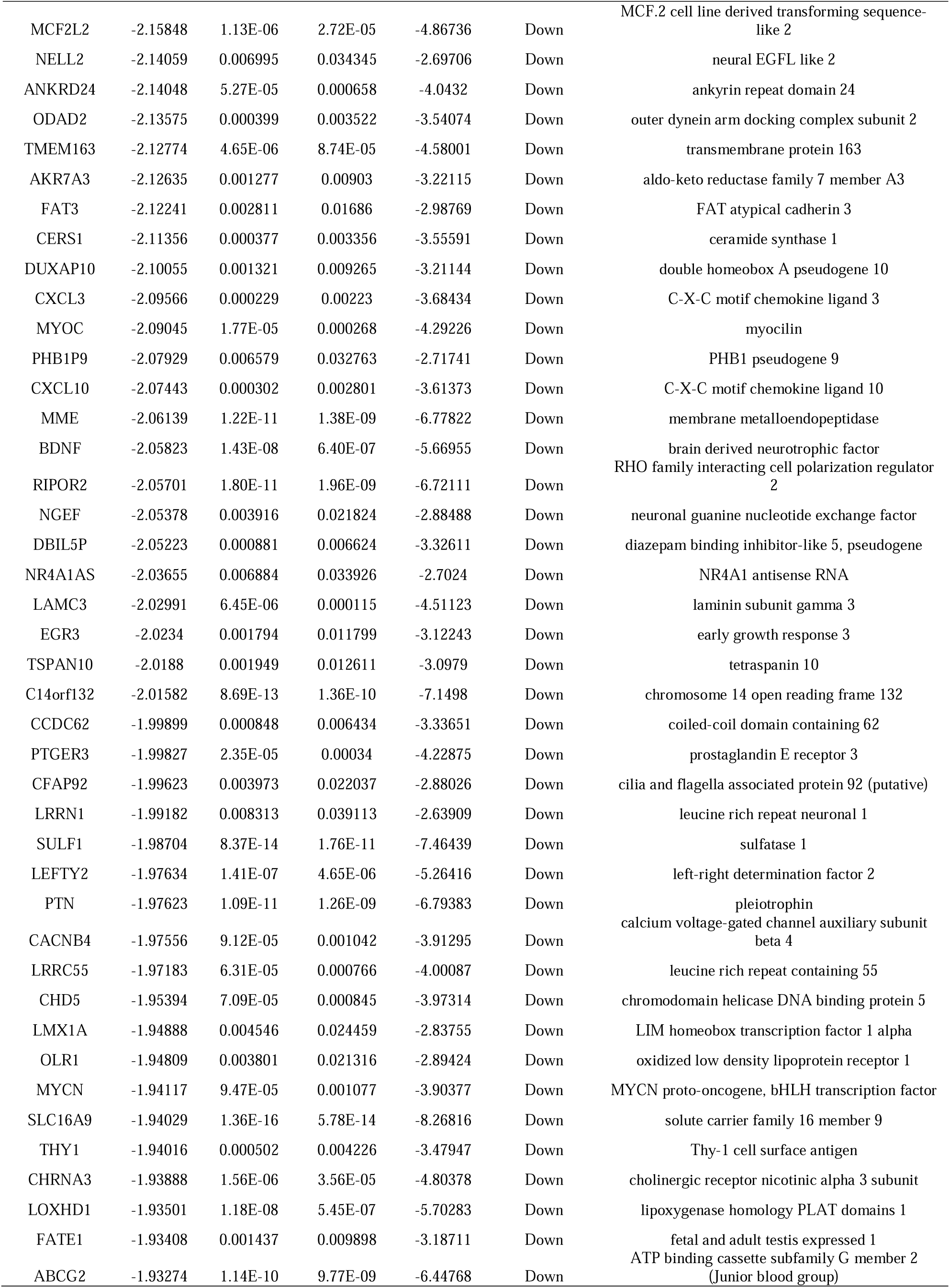
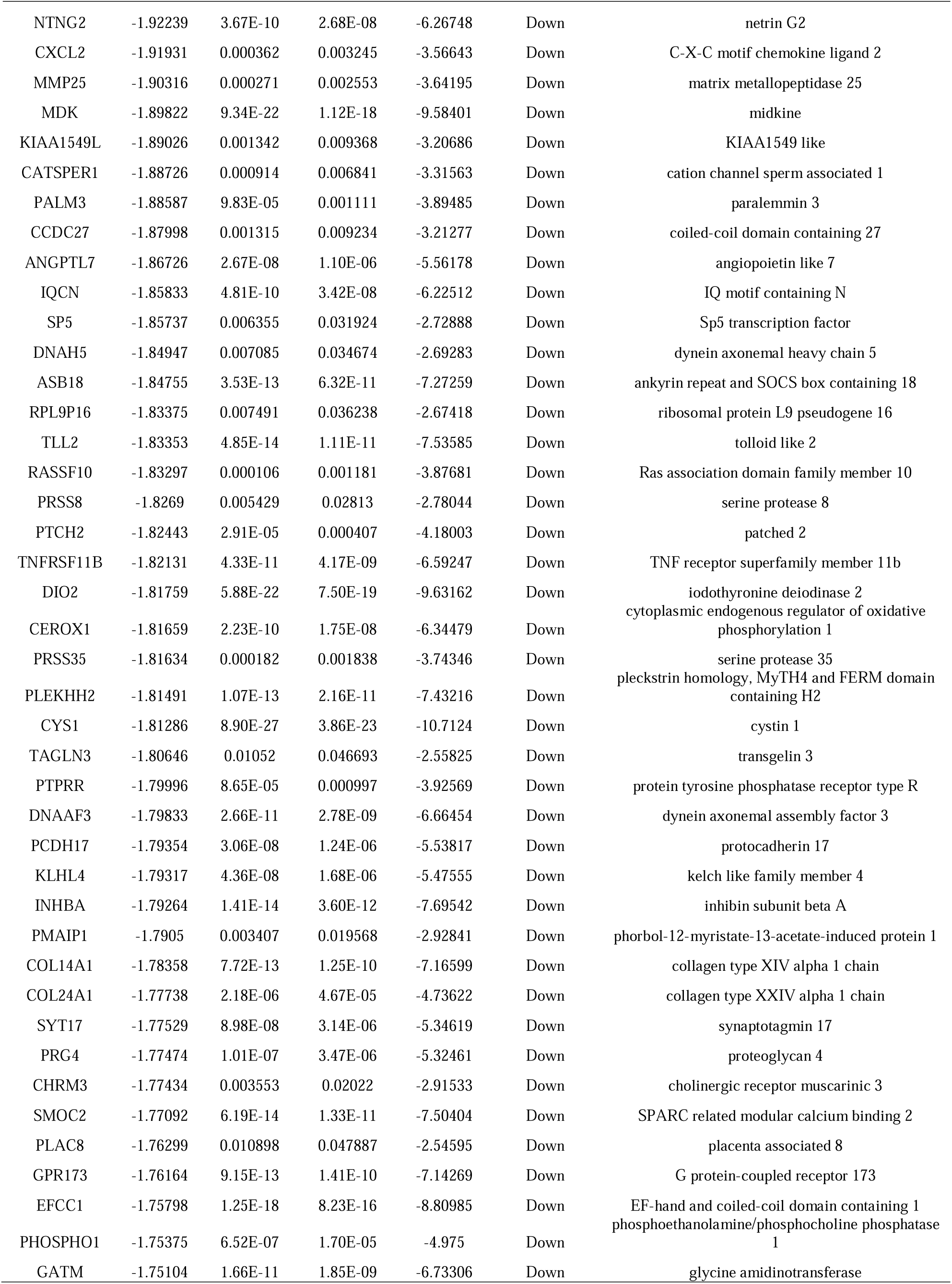
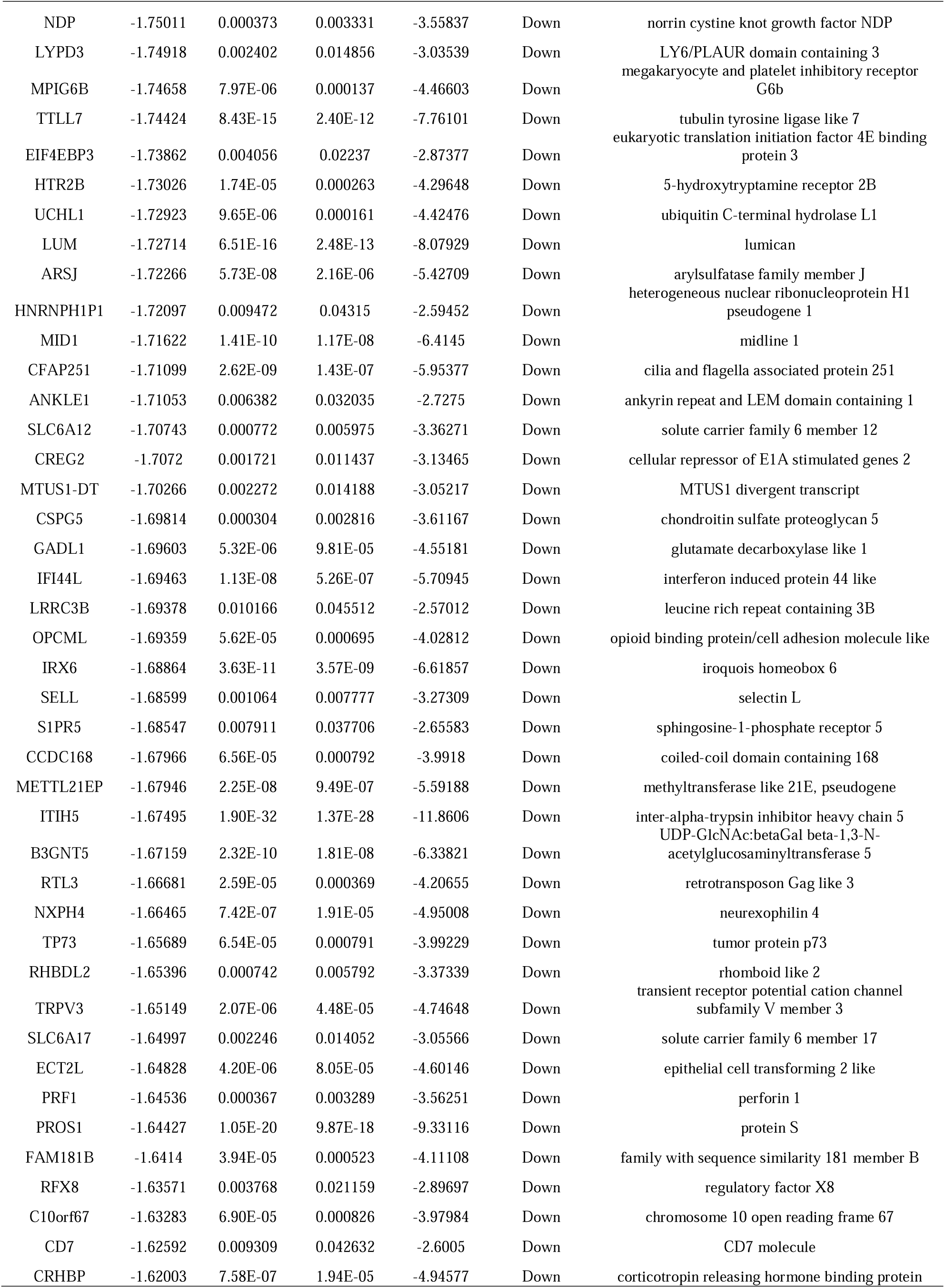
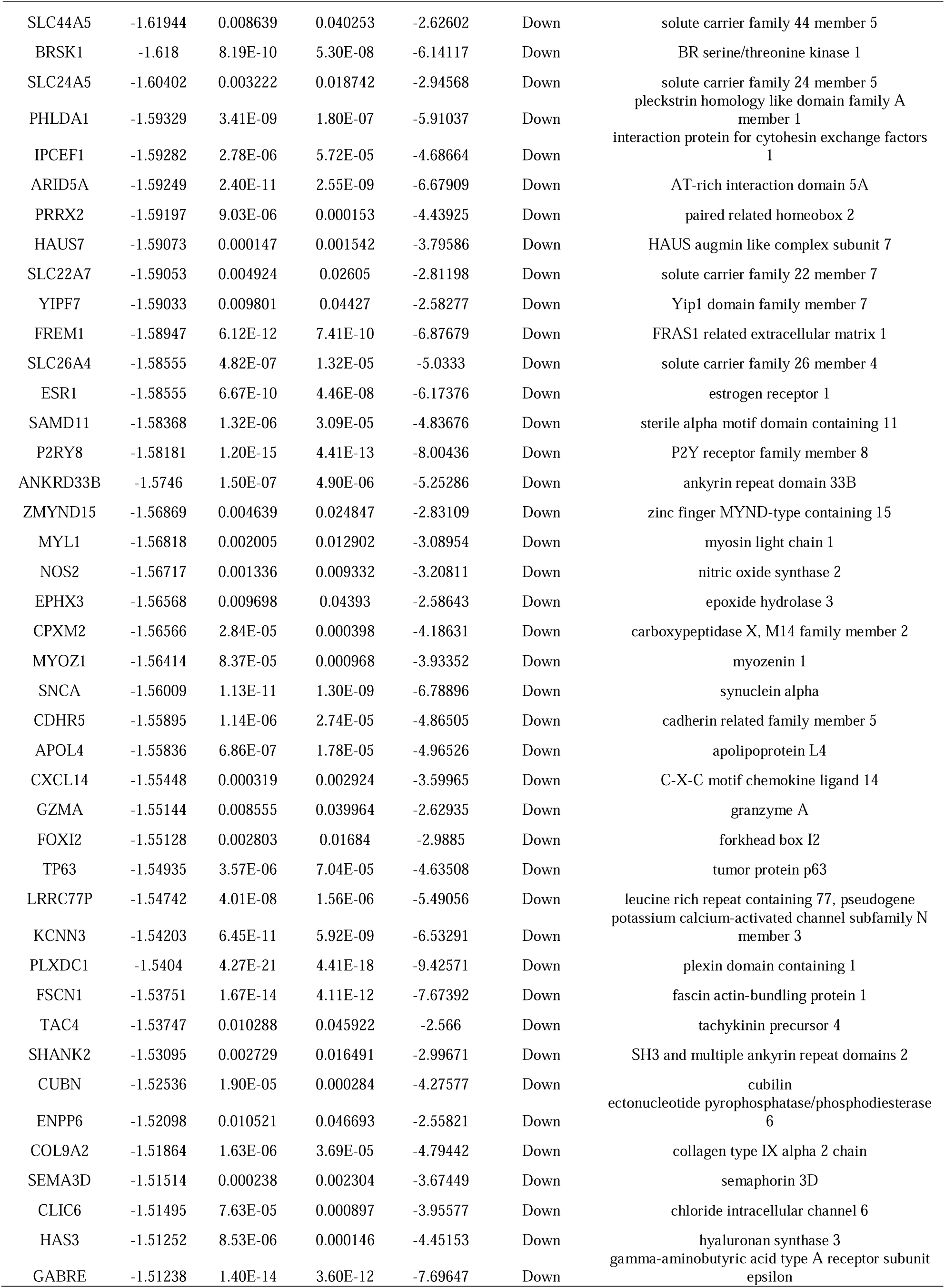
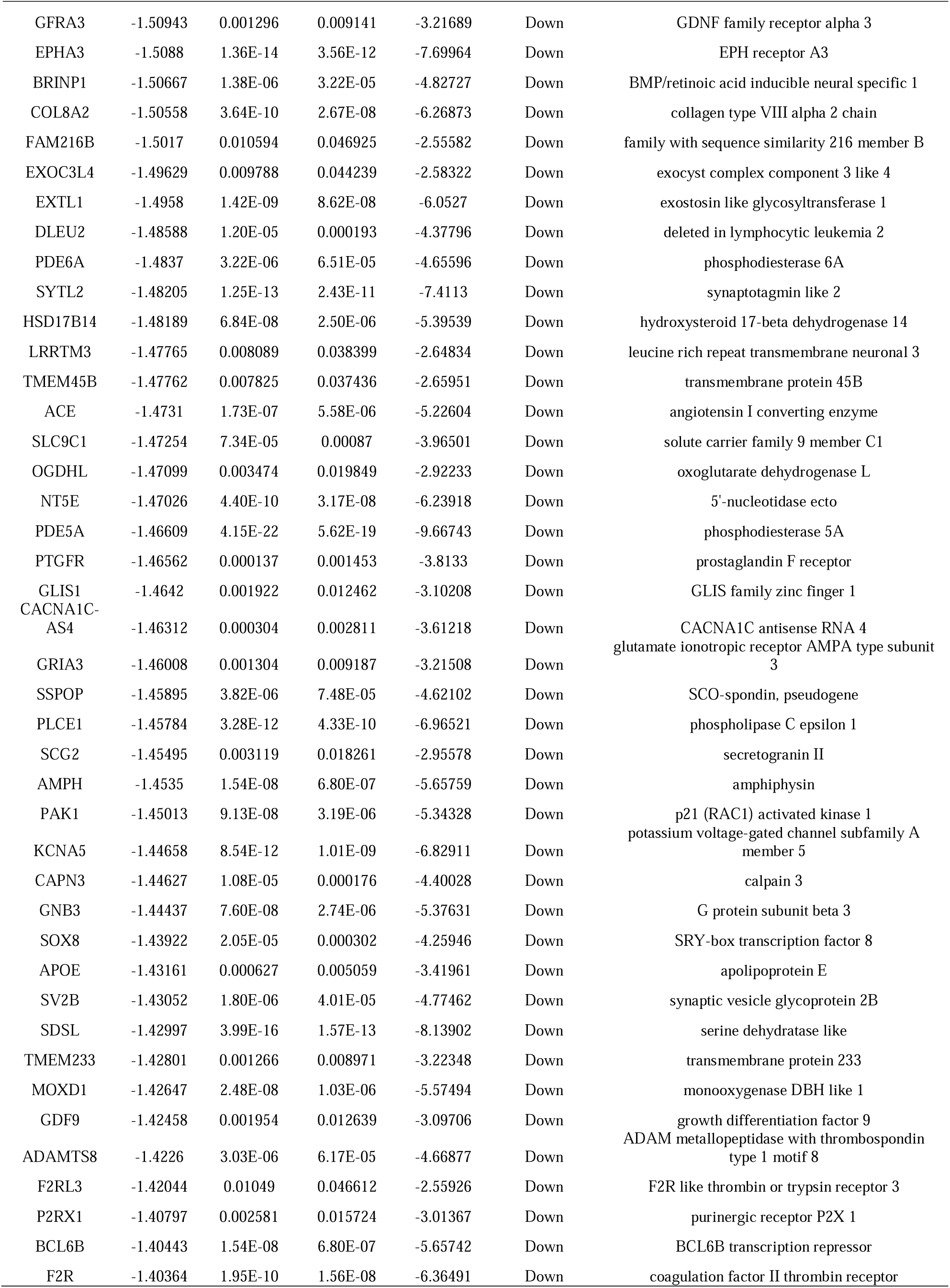
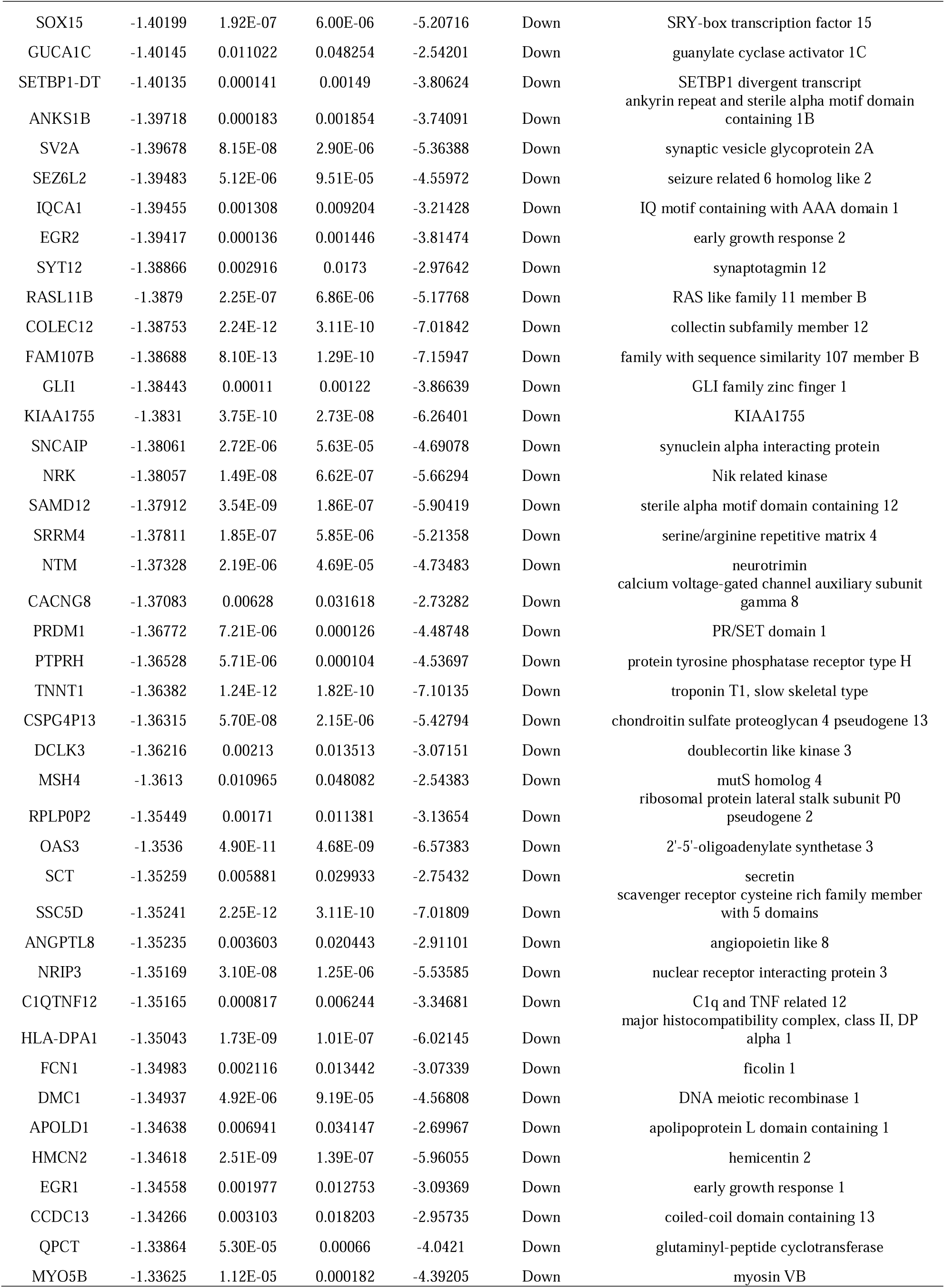
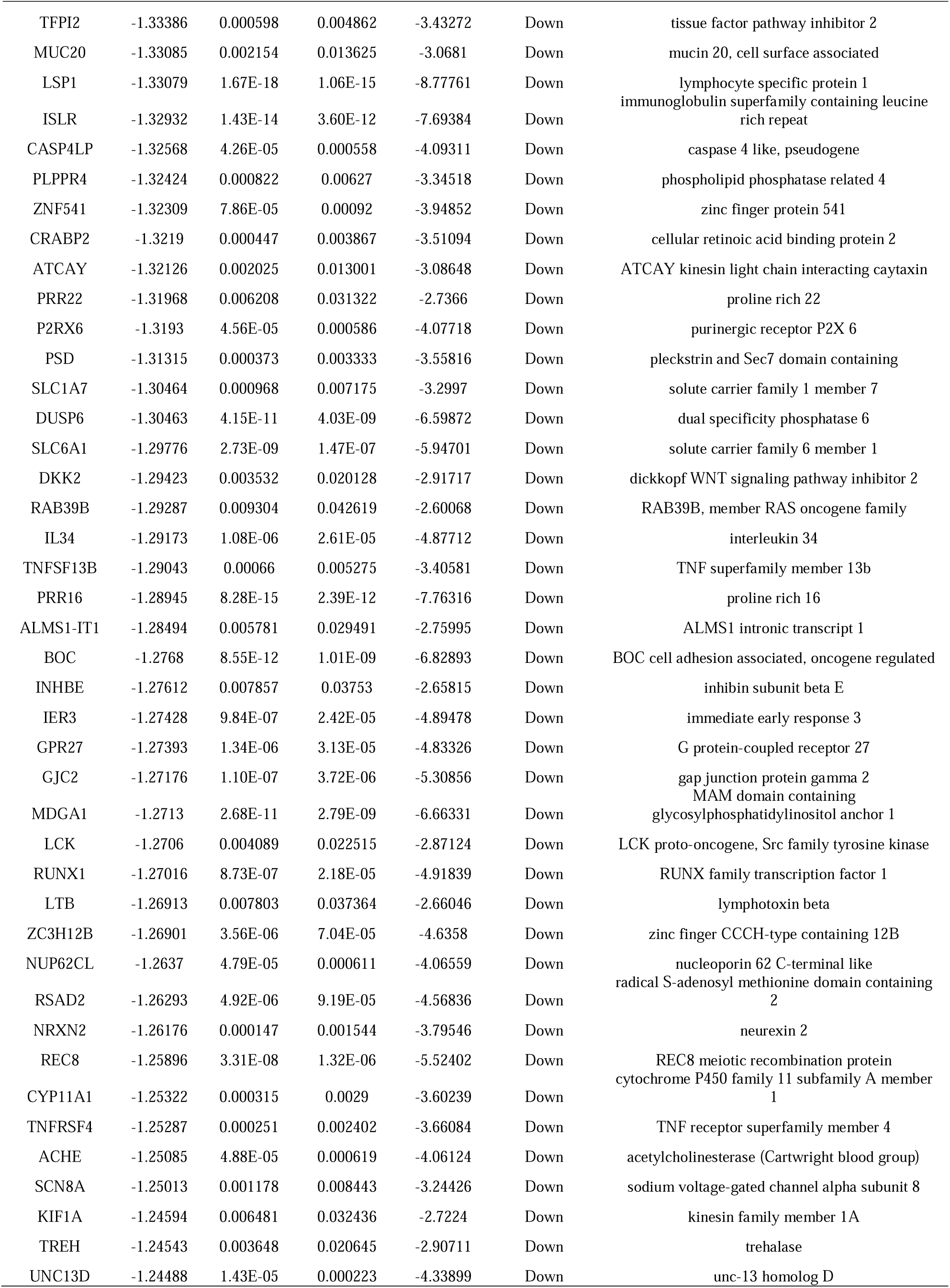
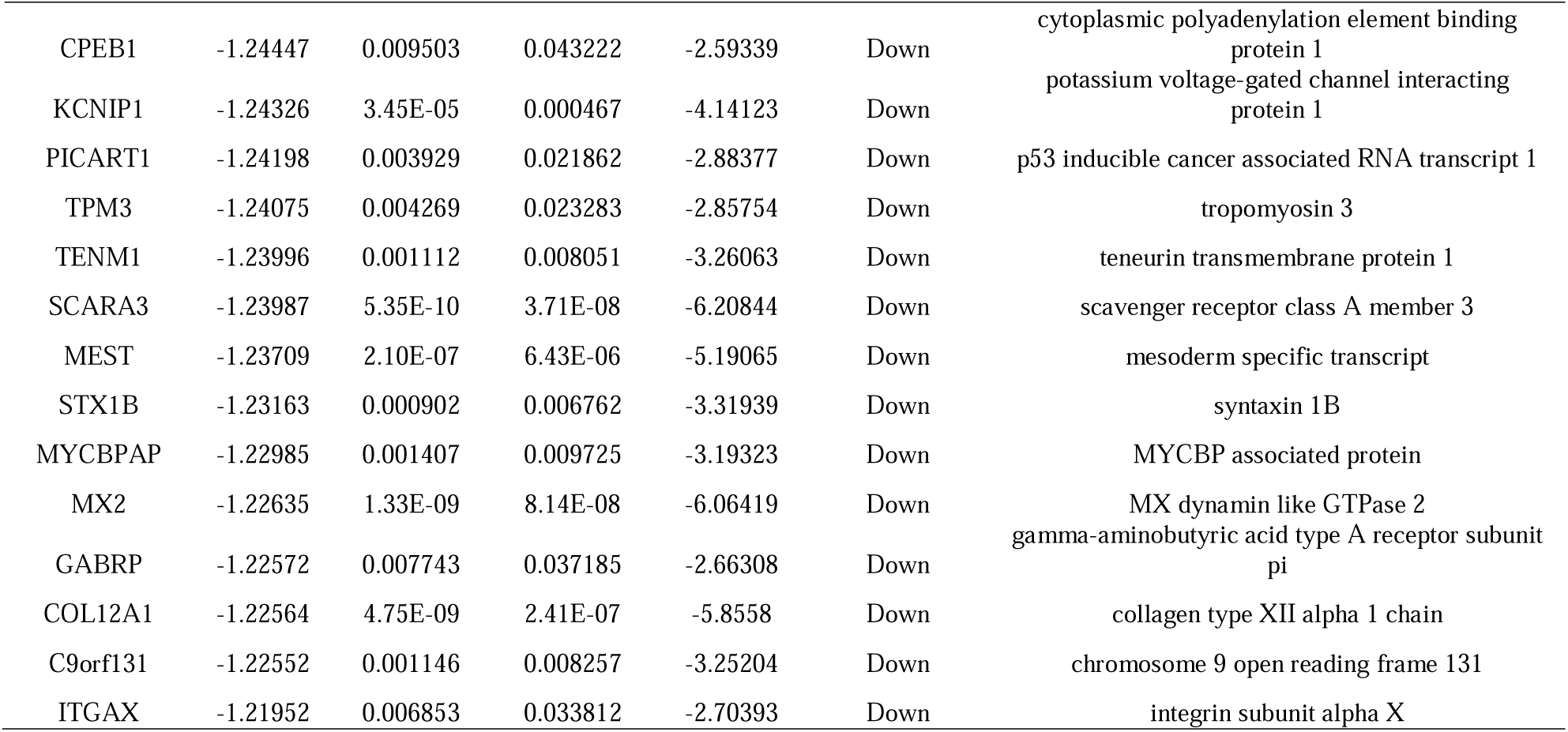
The statistical metrics for key differentially expressed genes (DEGs)

### GO and pathway enrichment analyses of DEGs

The GO enrichment analysis revealed the following most GO significant enrichment terms: (1) BP: response to stimulus, multicellular organism development, multicellular organismal process and cell communication; (2) CC: cell periphery, membrane, plasma membrane and cellular anatomical entity; and (3) MF: oxidoreductase activity, transporter activity, signaling receptor binding and molecular function regulator activity (Table 3 and Fig.4 to Fig.5). The REACTOME pathway enrichment analysis revealed that the DEGs were particularly enriched in metabolism, cardiac conduction, extracellular matrix organization and GPCR ligand binding (Table 4 and Fig.4 to Fig.5).

**Fig. 4.**
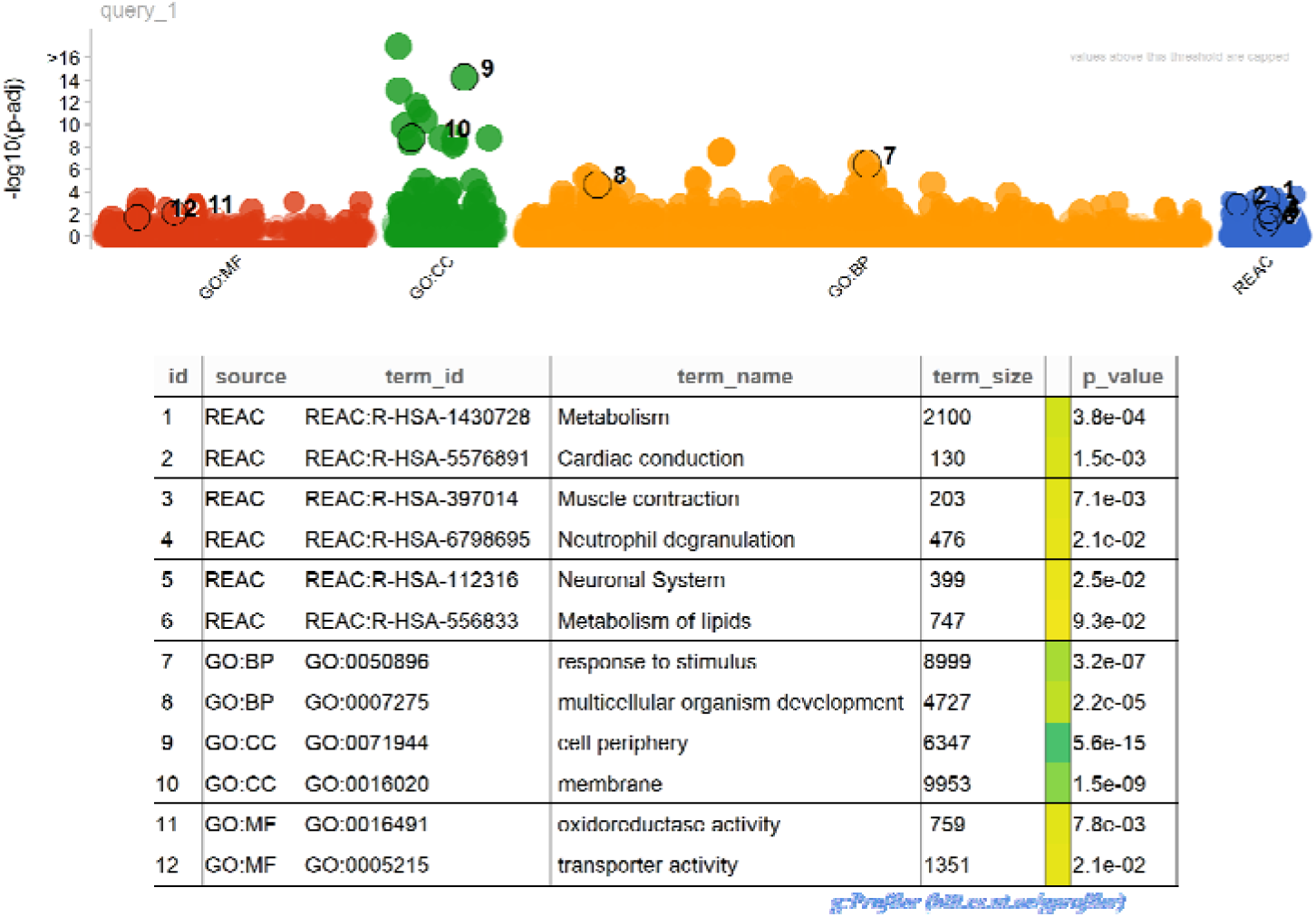
GO and REACTOME pathway enrichment analysis for up regulated genes. p < 0.05. Abbreviations: BP, biological process; CC, cell component; MF, molecular function. GO, Gene Ontology; REAC, REACTOME. The size of the circle represents the number of genes involved, and the abscissa represents the frequency of the genes involved in the term total genes.

**Fig. 5.**
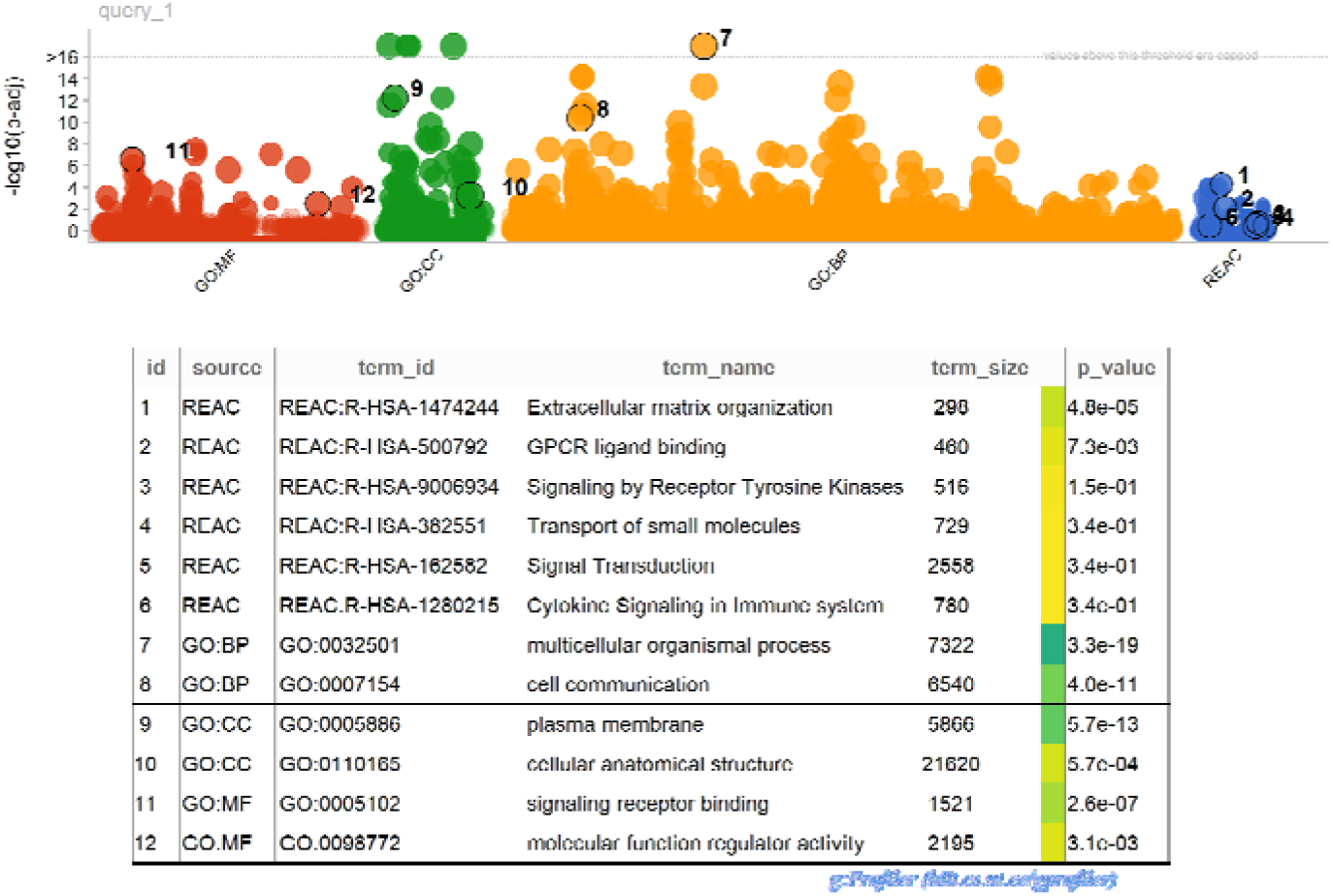
GO and REACTOME pathway enrichment analysis for down regulated genes. p < 0.05. Abbreviations: BP, biological process; CC, cell component; MF, molecular function. GO, Gene Ontology; REAC, REACTOME. The size of the circle represents the number of genes involved, and the abscissa represents the frequency of the genes involved in the term total genes.

**Table 3.**
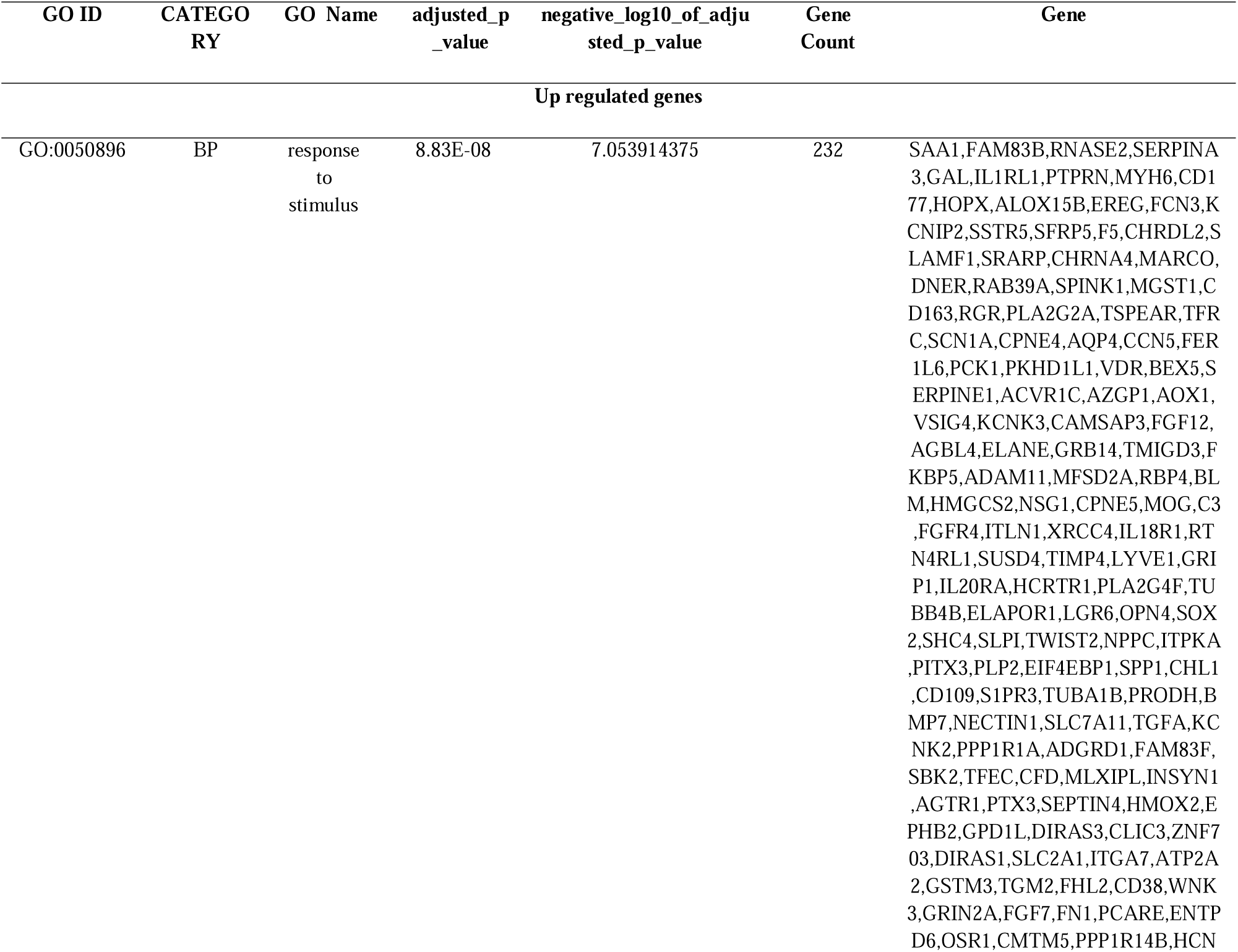
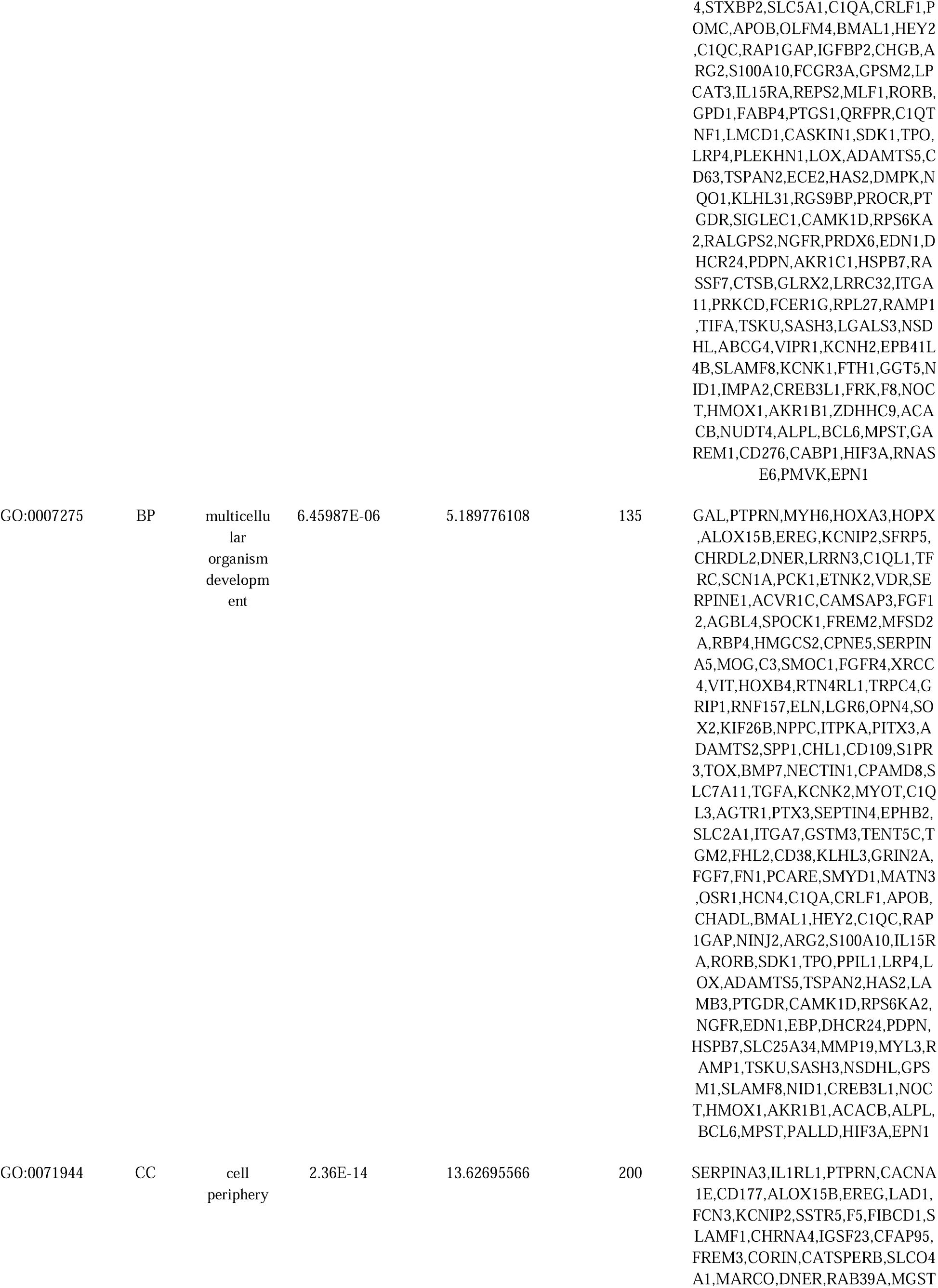
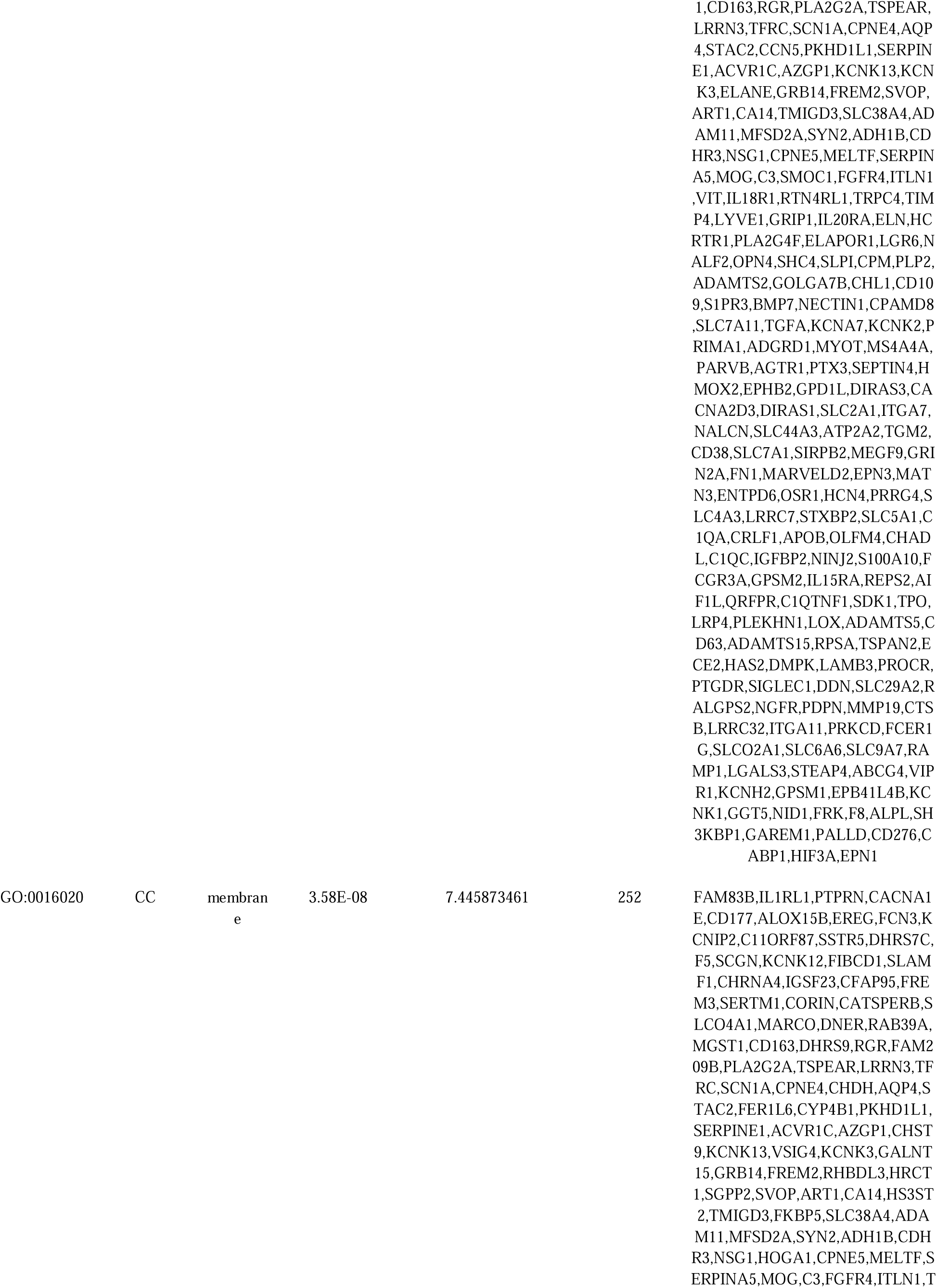
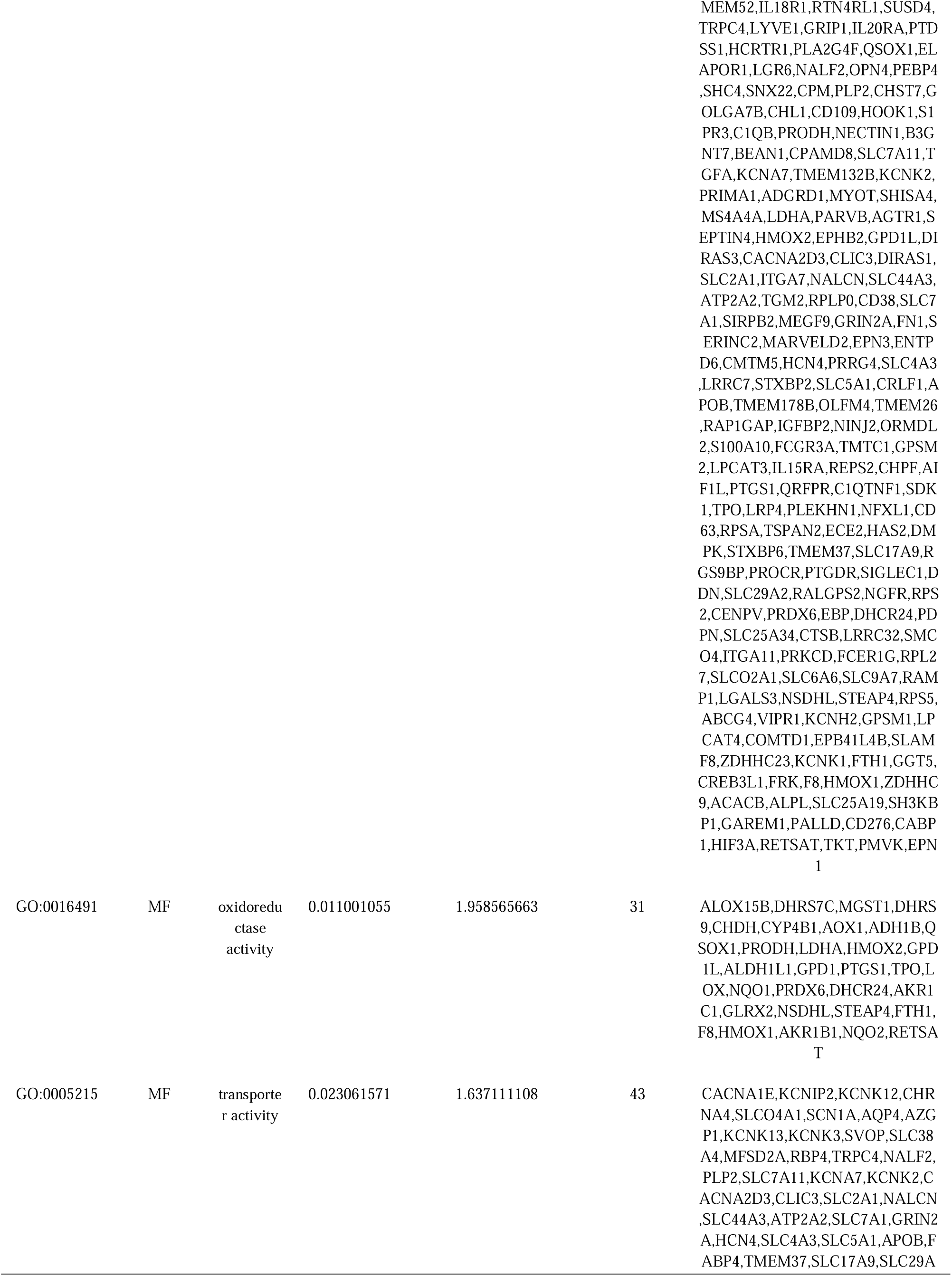
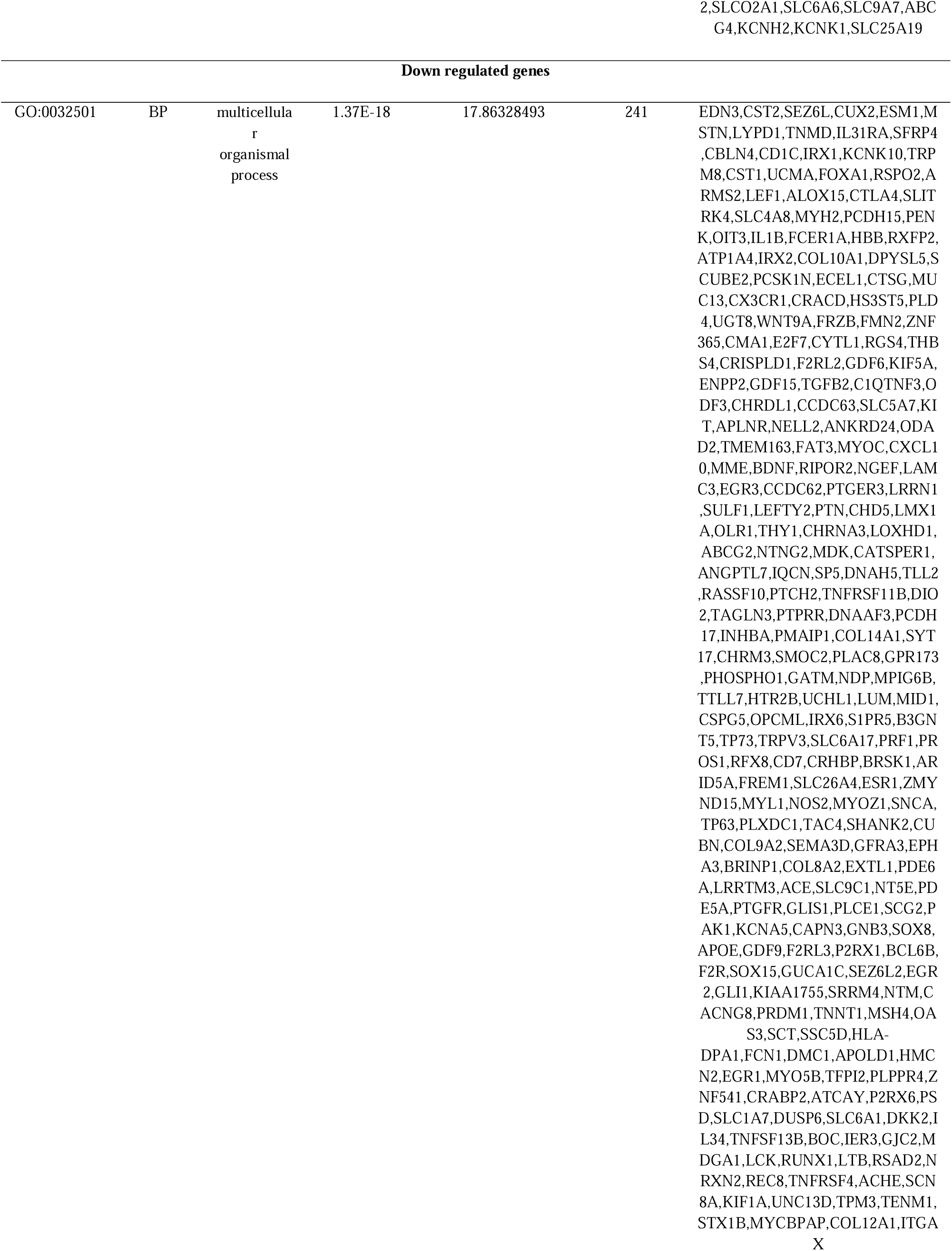
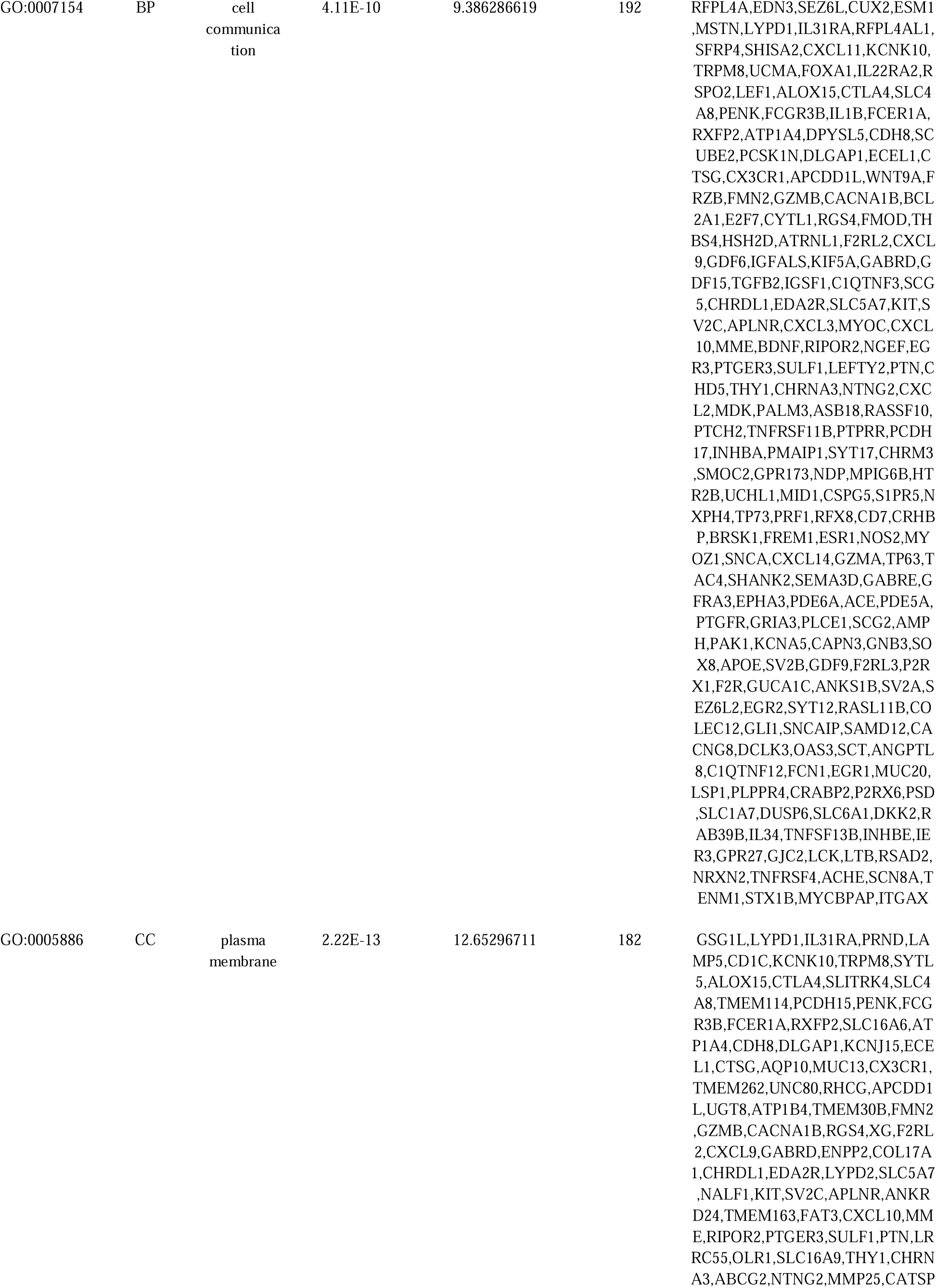
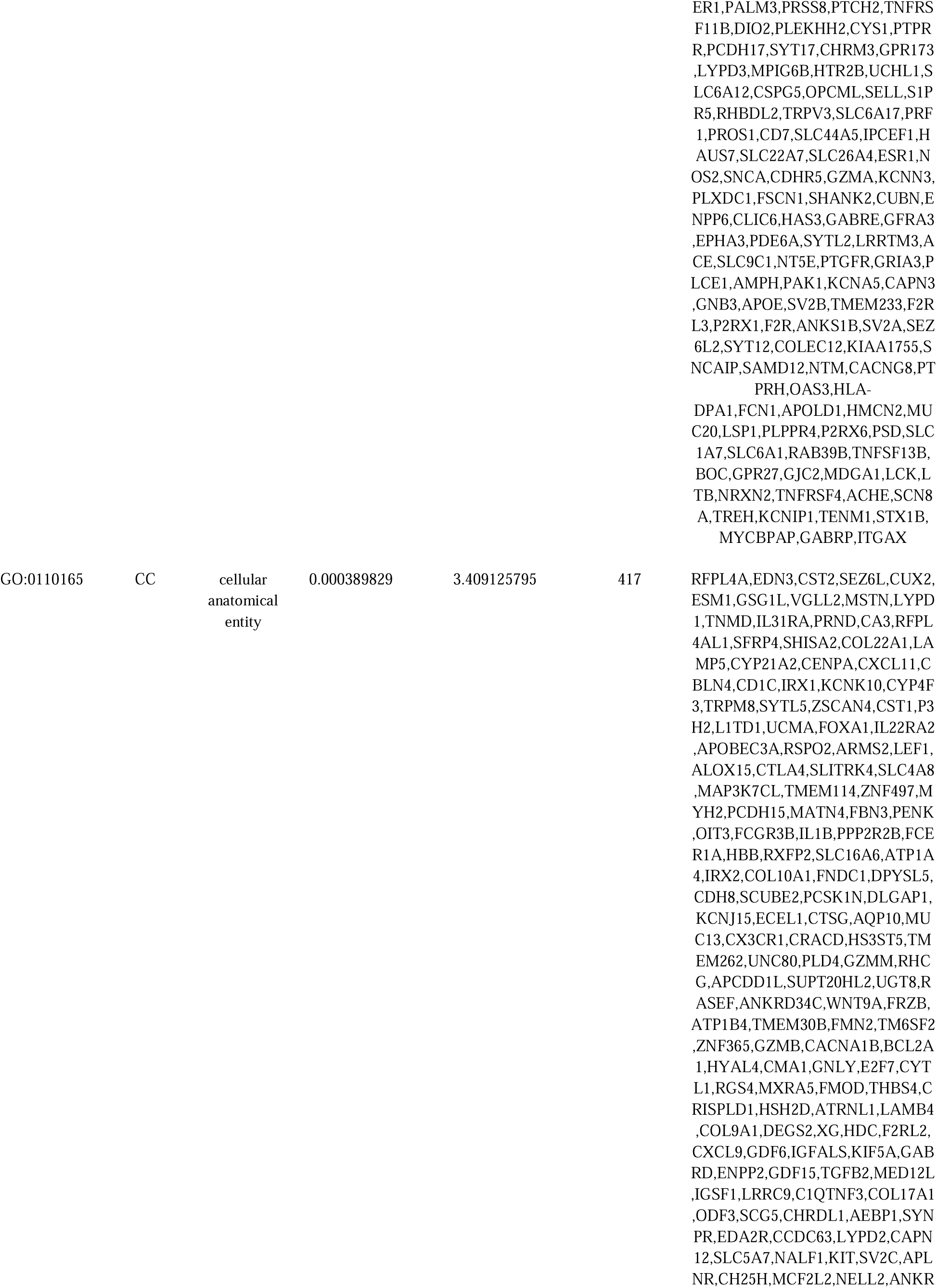
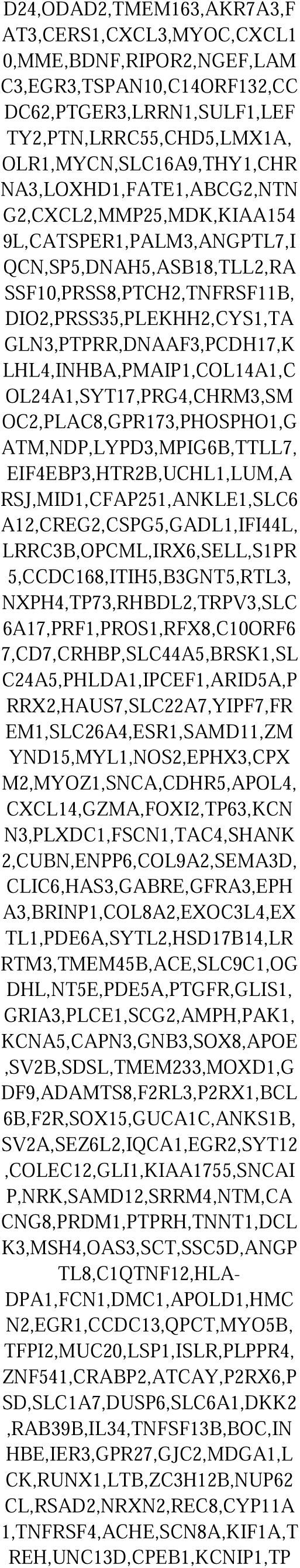
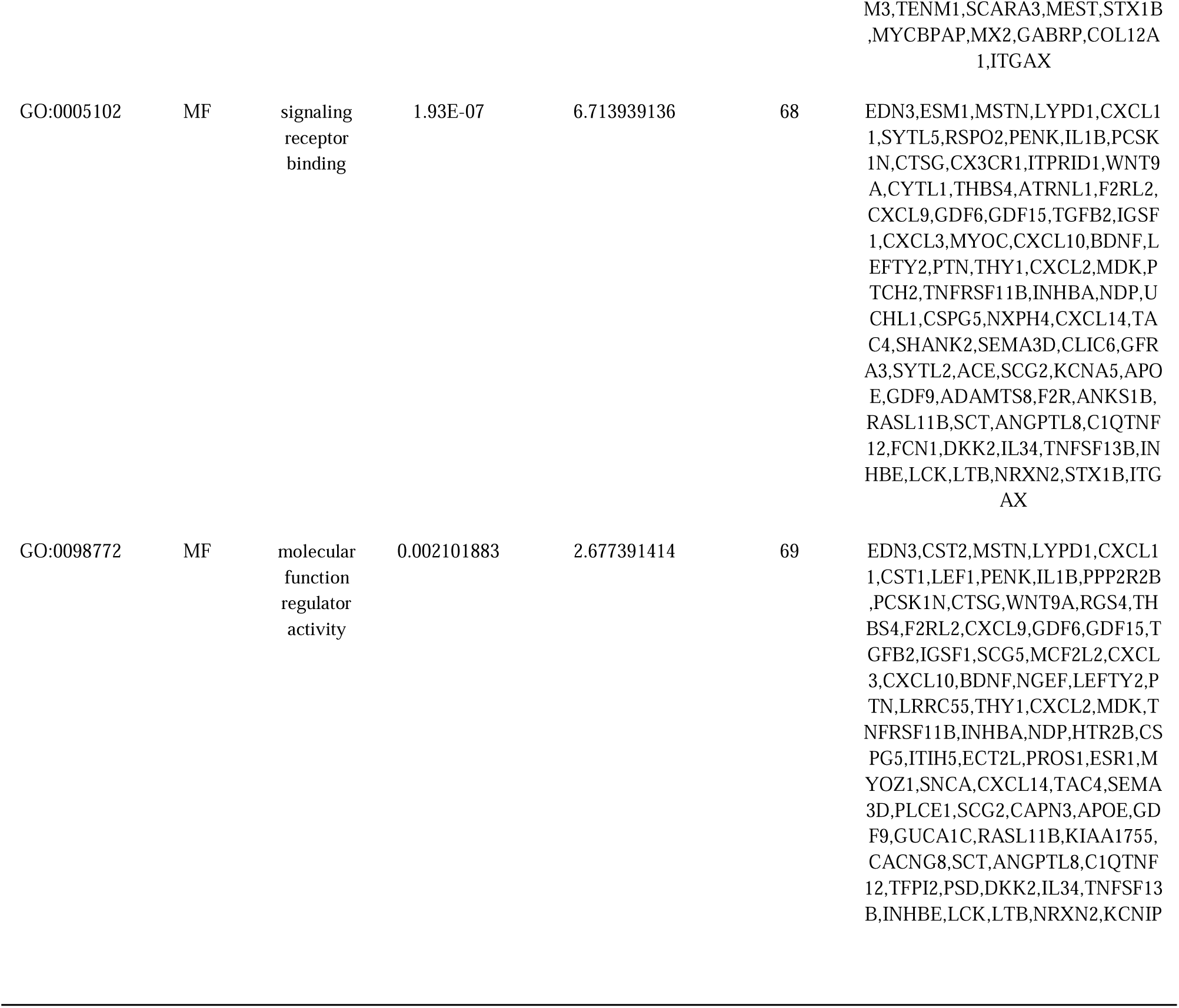
The enriched GO terms of the up and down regulated differentially expressed genes.

**Table 4.**
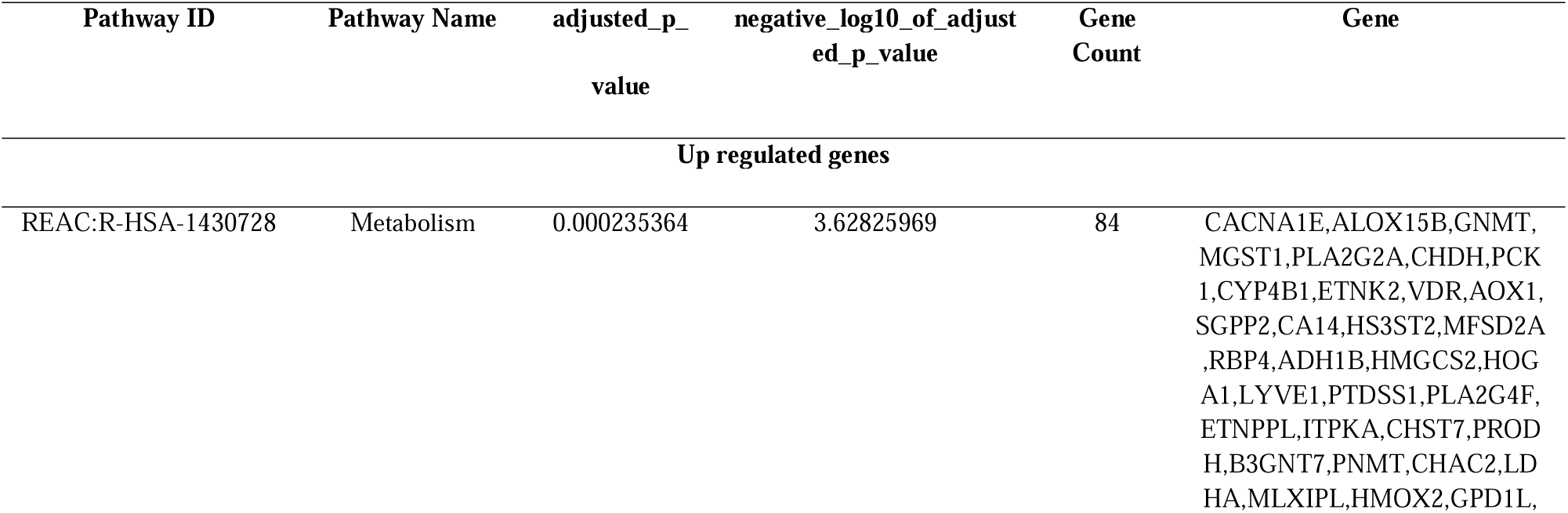
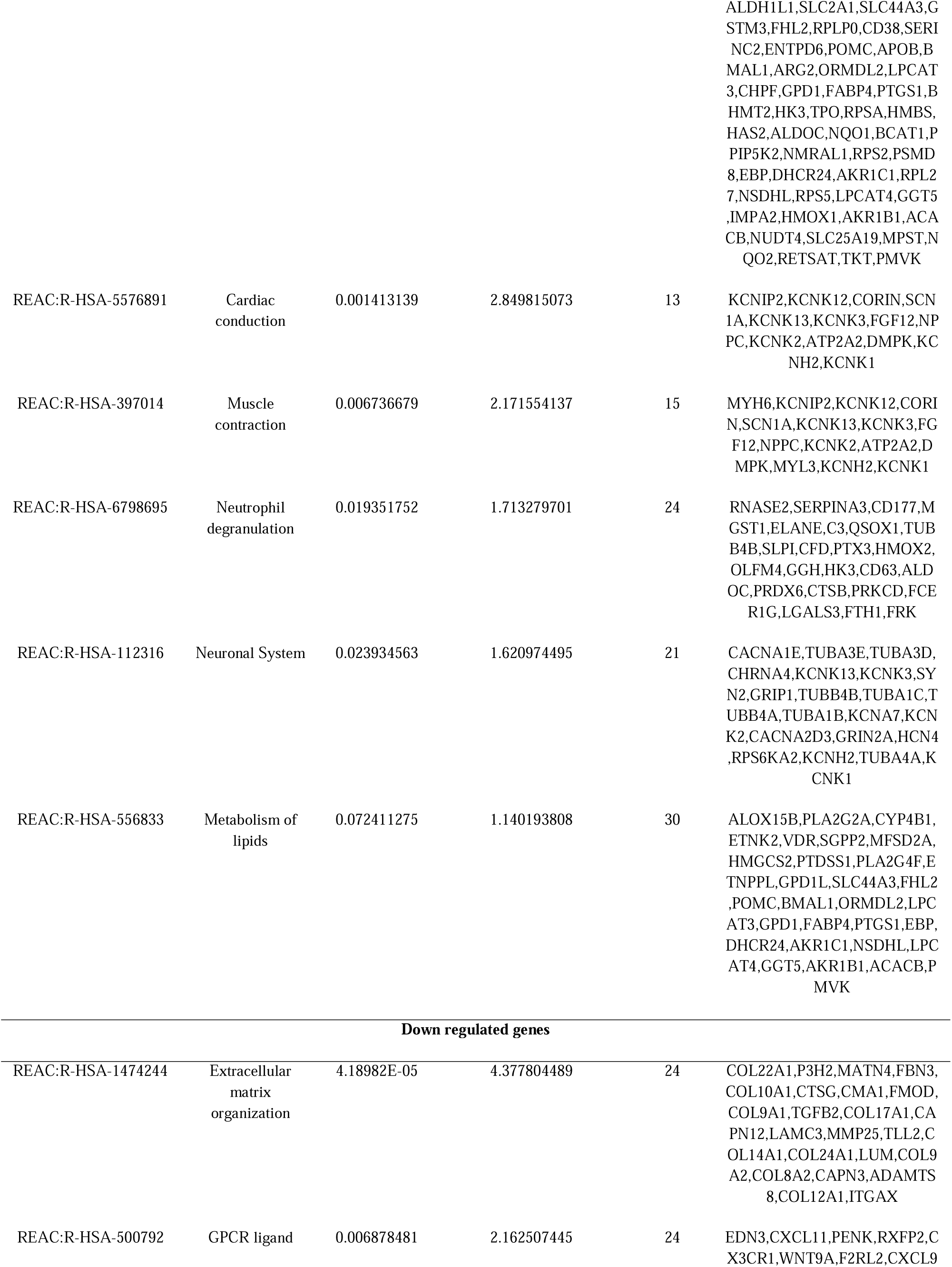
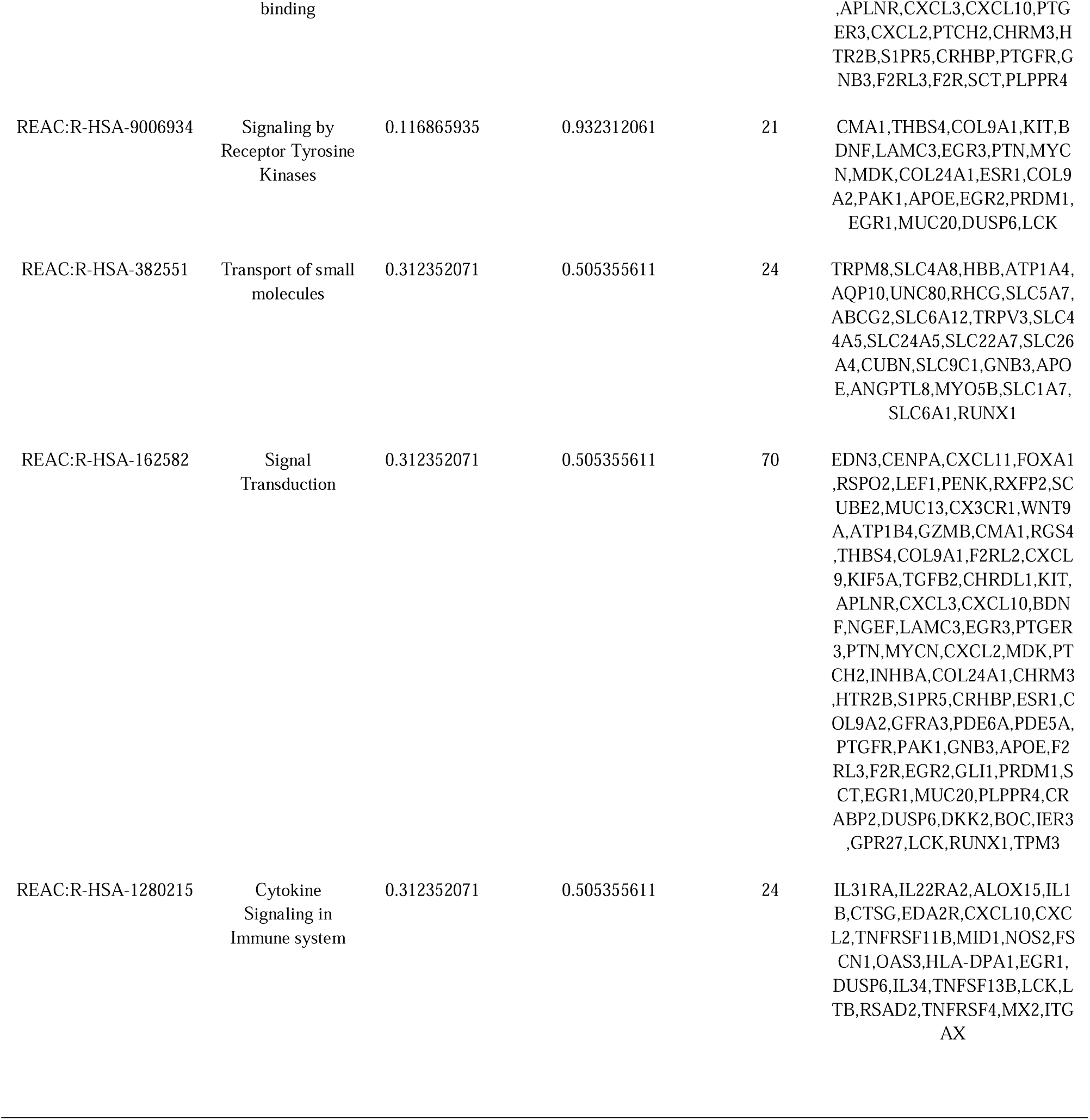
The enriched pathway terms of the up and down regulated differentially expressed genes.

### Construction of the PPI network and module analysis

The visual PPI network of 958 DEGs was constructed using Cytoscape software. The PPI network was composed of 5887 nodes and 10291 edges (Fig. 6). Among the 5887 nodes, 958 hub genes with the highest nodes degree, betweenness, stress and closeness were screened based on the Cytoscape software analysis results. The results were as follows: FN1, SOX2, TUBA4A, RPS2, TUBA1C, ESR1, SNCA, LCK, PAK1 and APLNR (Table 5). The two significant modules with the highest degree were screened, and GO and pathway enrichment of genes in these two modules was analyzed using g:Profiler online tools. The key modules were obtained using PEWCC. Module 1 contained 53 nodes and 66 edges (Fig. 7), mainly involved in response to stimulus, multicellular organism development and cell periphery (Fig. 8). Module 2 was comprised of 36 nodes and 40 edges (Fig. 9), which were associated with multicellular organismal process, cell communication and cytokine signaling in immune system (Fig. 10).

**Fig. 6.**
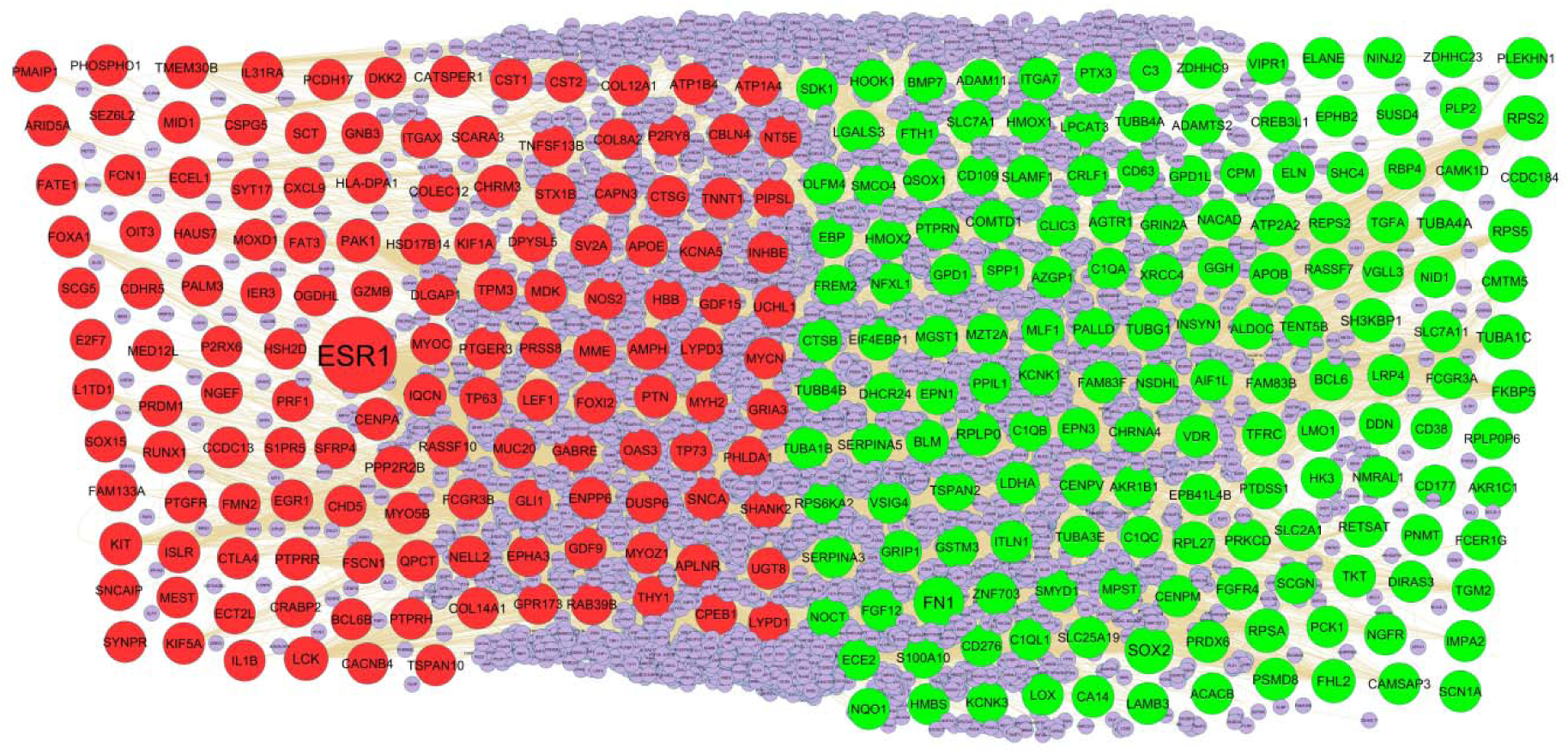
PPI network of DEGs. Up regulated genes are marked in parrot green; down regulated genes are marked in red.

**Fig. 7.**
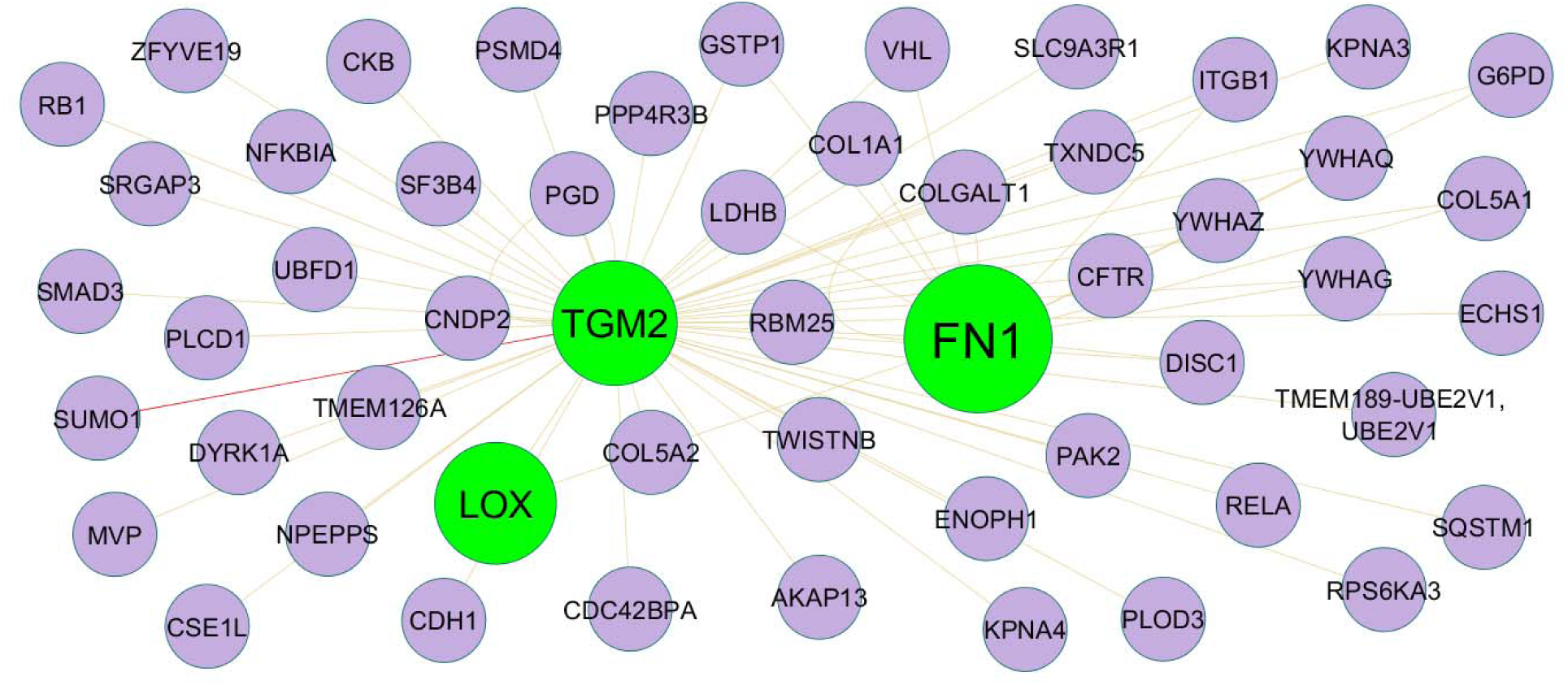
Modules 1 was isolated form PPI of up regulated genes. Module 1 has 53 nodes and 66 edges for up regulated genes. Up regulated genes are marked in green

**Fig. 8.**
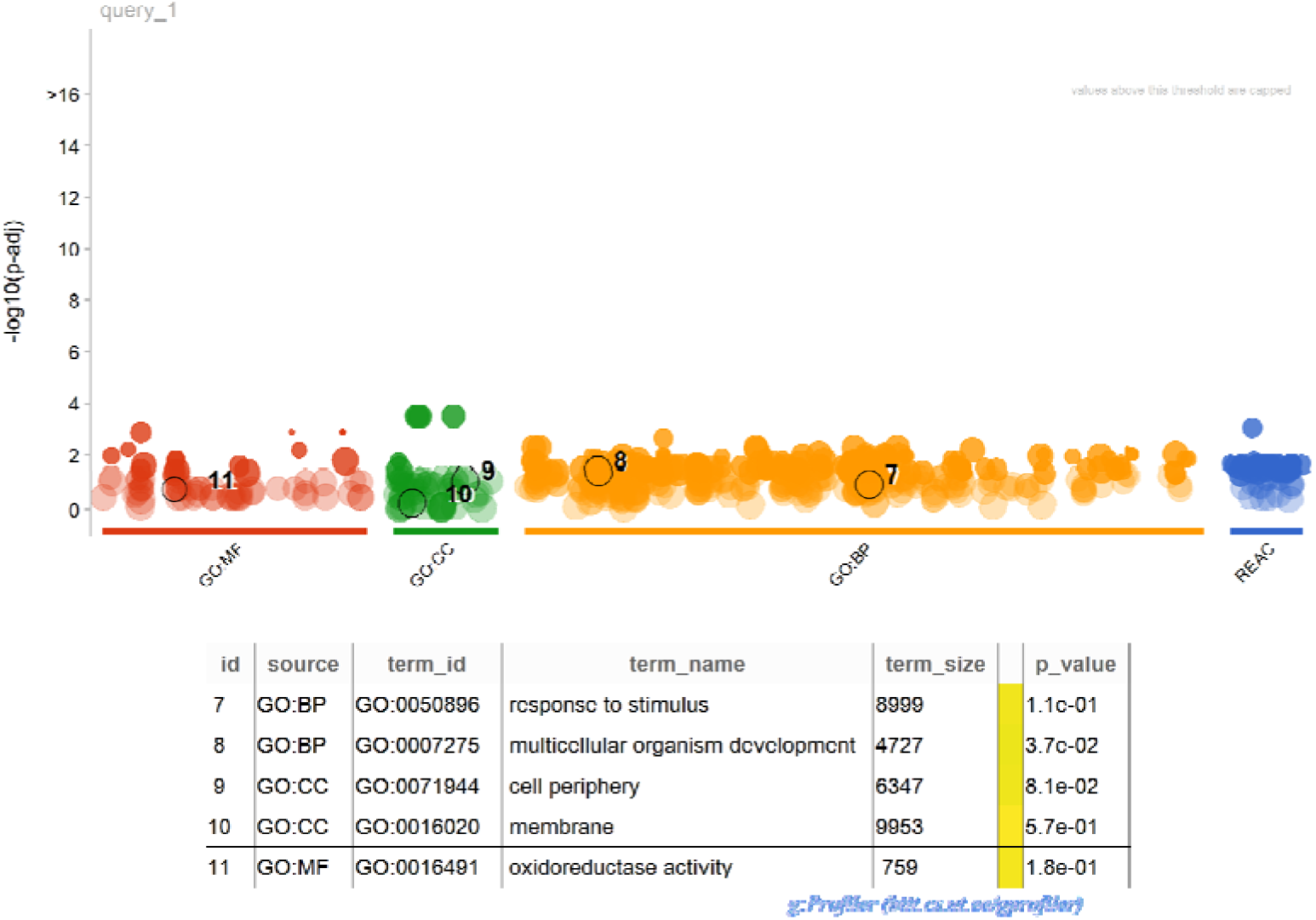
Enrichment analysis for module 1. The size of the circle represents the number of genes involved, and the abscissa represents the freqency of the genes involved in the term total genes.

**Fig. 9.**
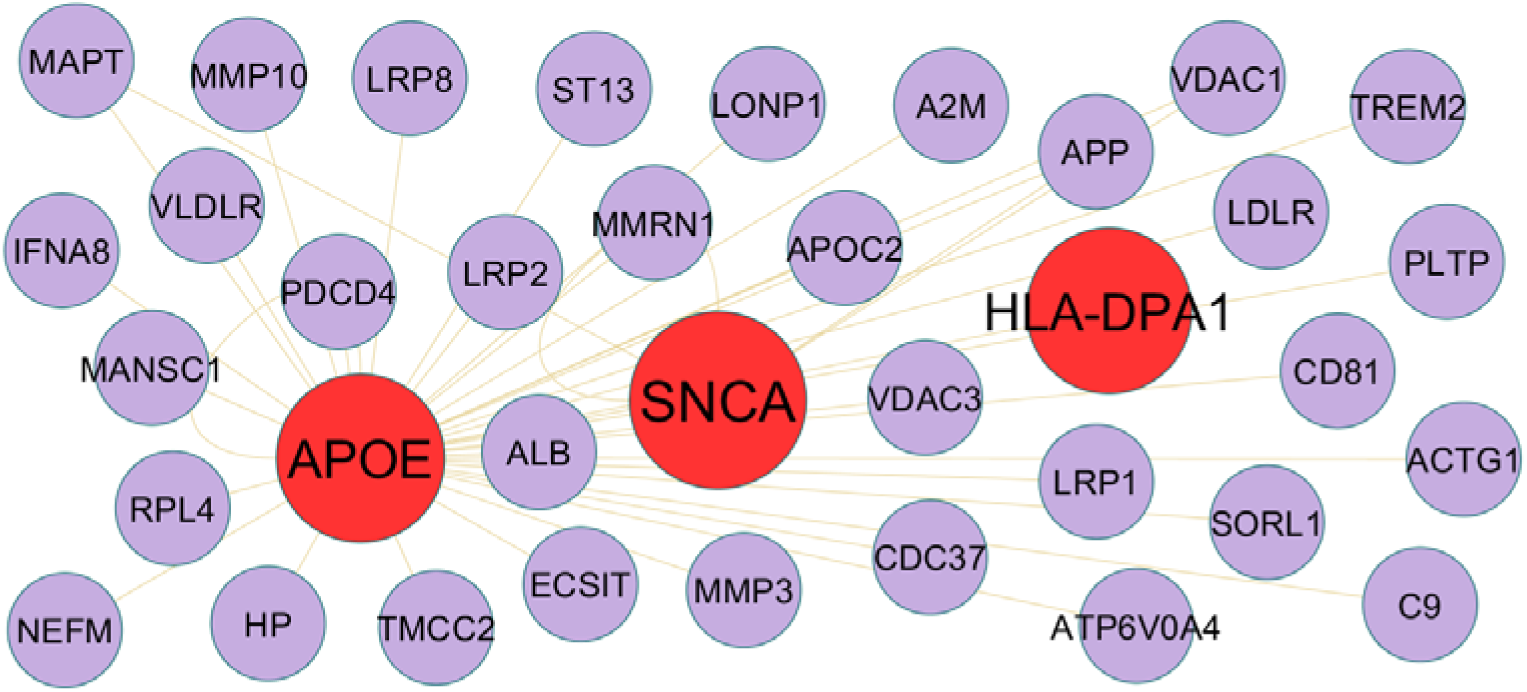
Module 2 was isolated form PPI of up regulated genes. Module 2 has 36 nodes and 40 edges for down regulated genes. Down regulated genes are marked in red

**Fig. 10.**
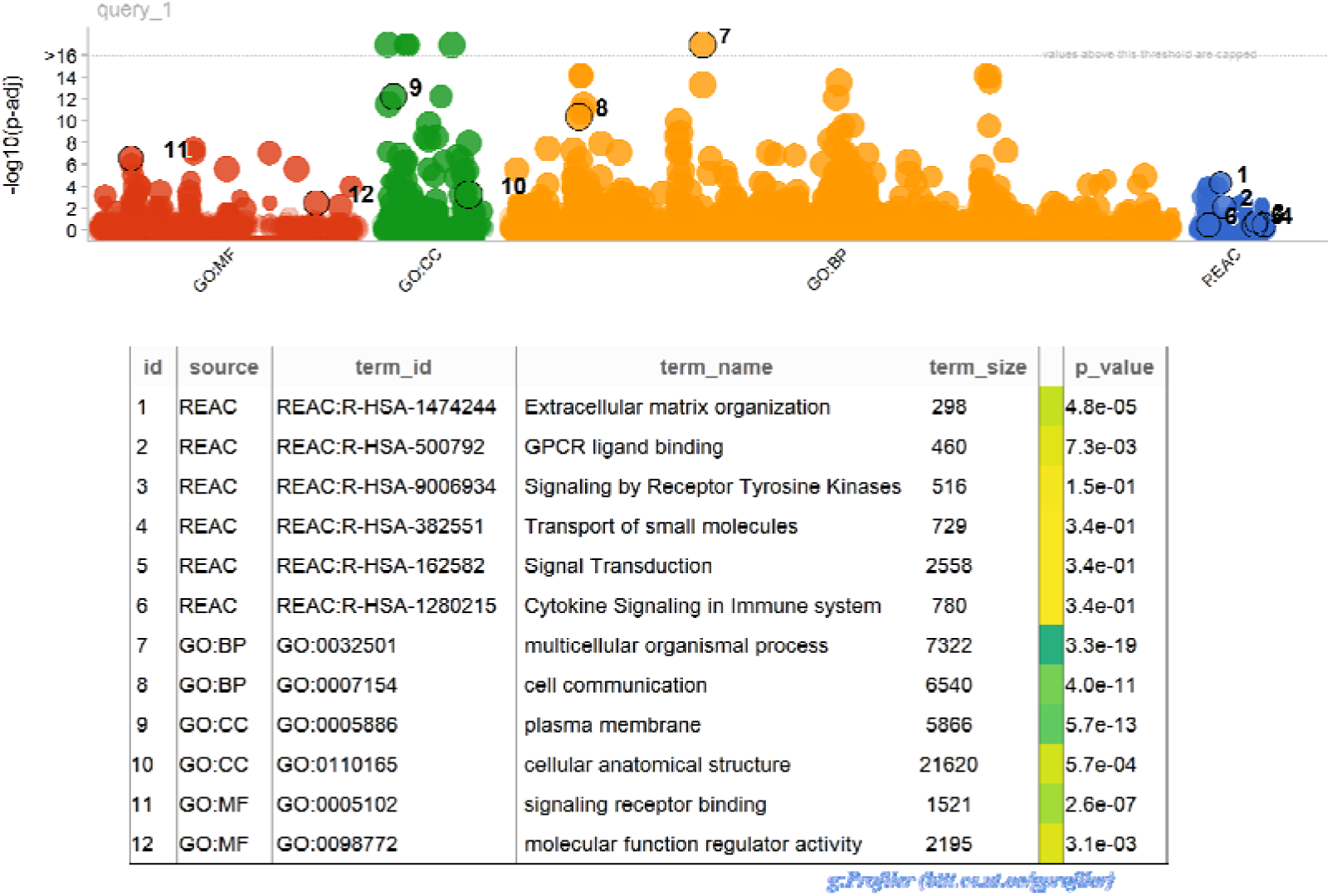
Enrichment analysis for module 2. The size of the circle represents the number of genes involved, and the abscissa represents the frequency of the genes involved in the term total genes.

**Table 5.**
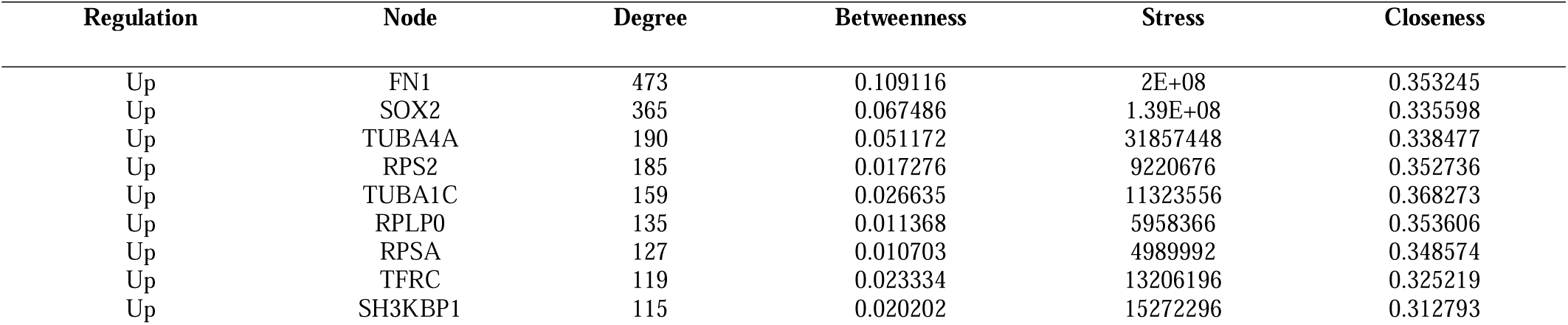
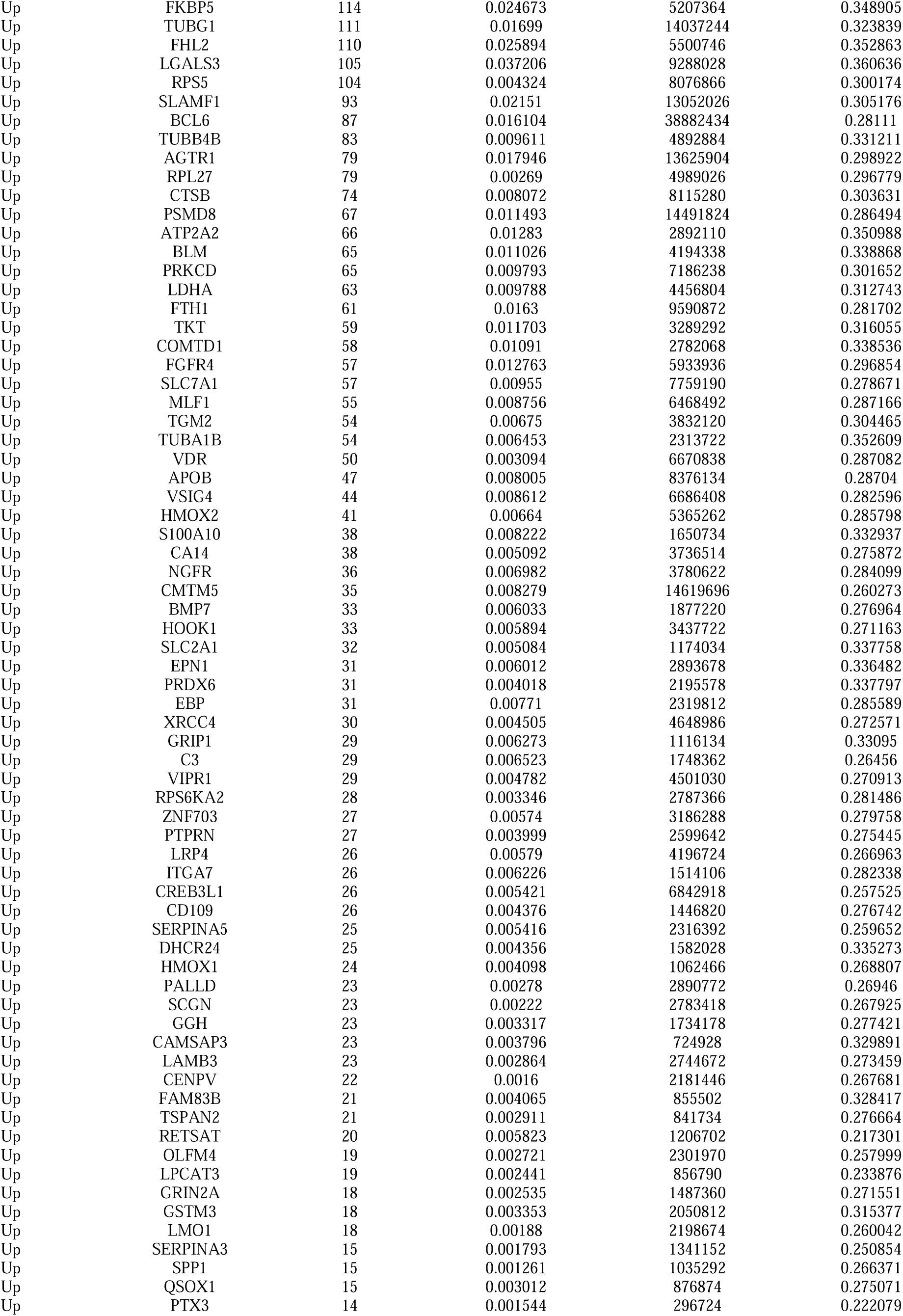
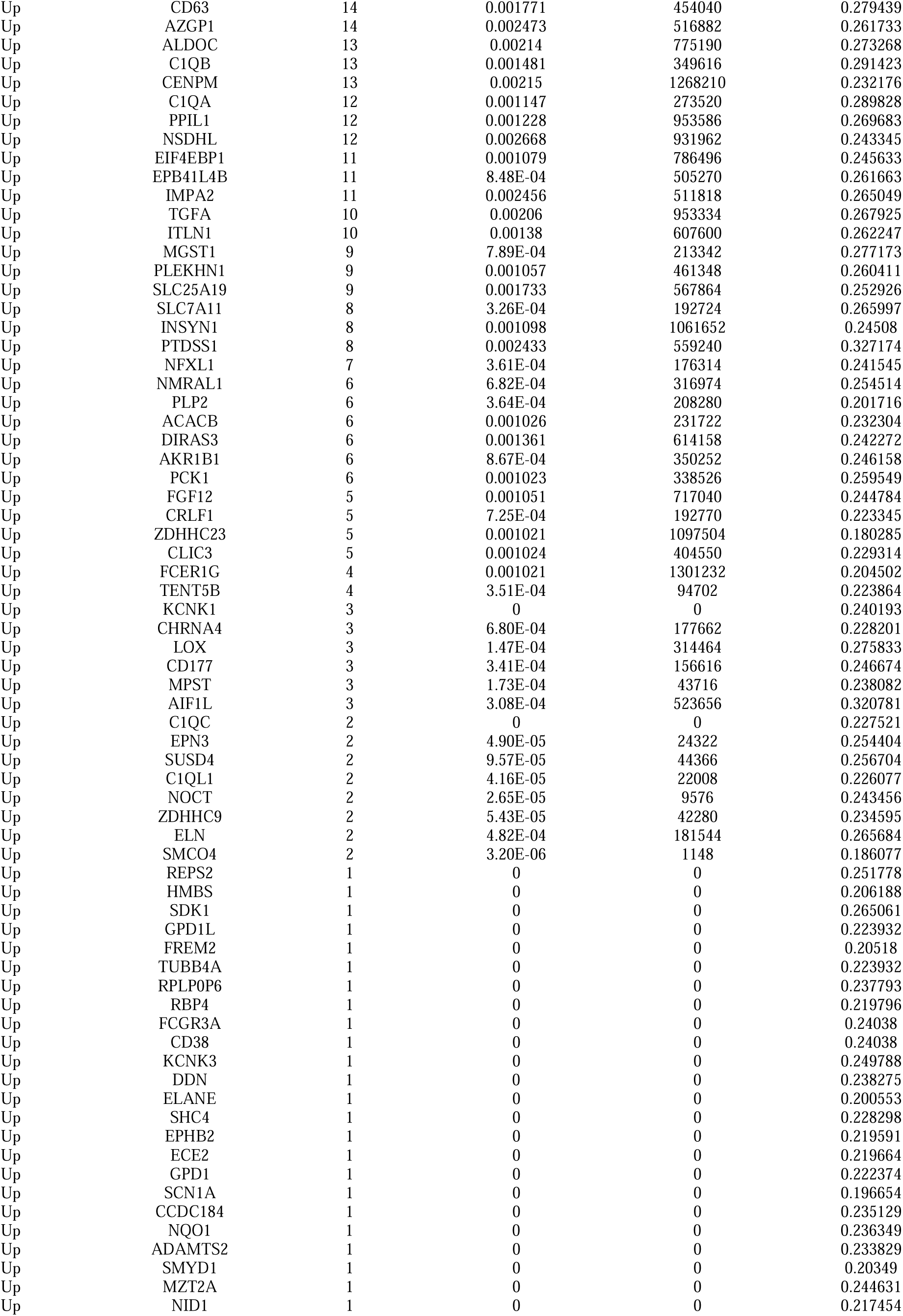
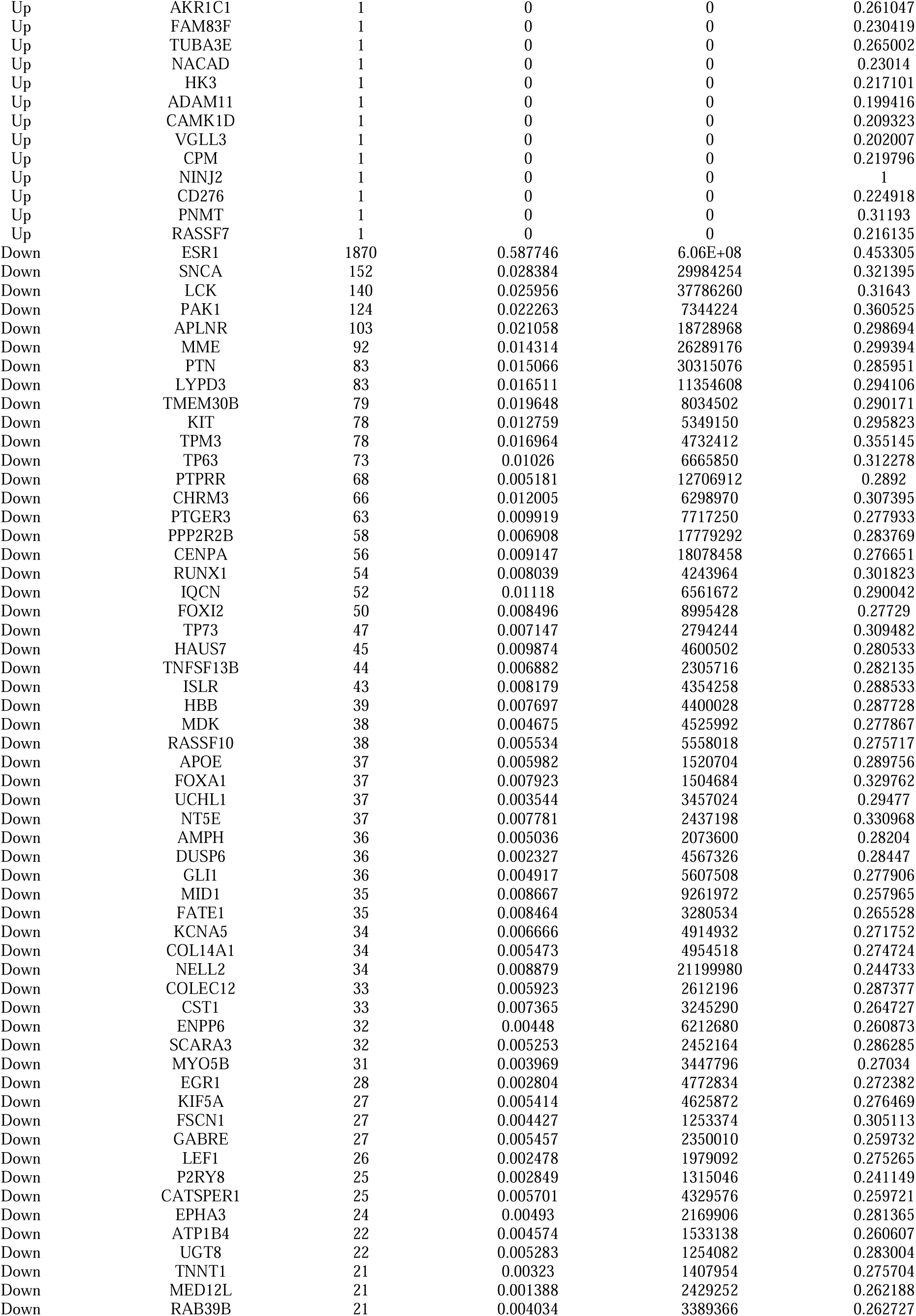
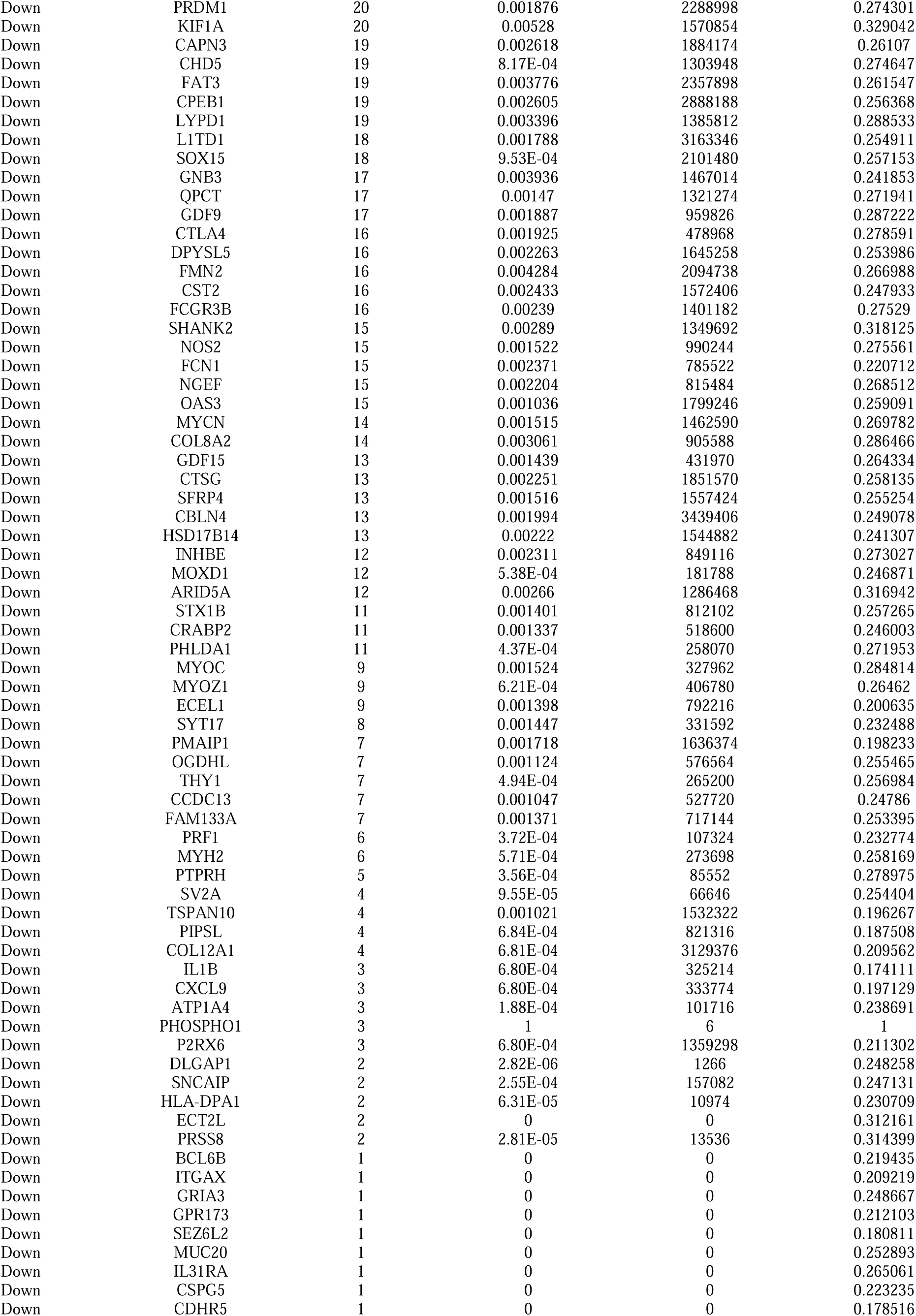
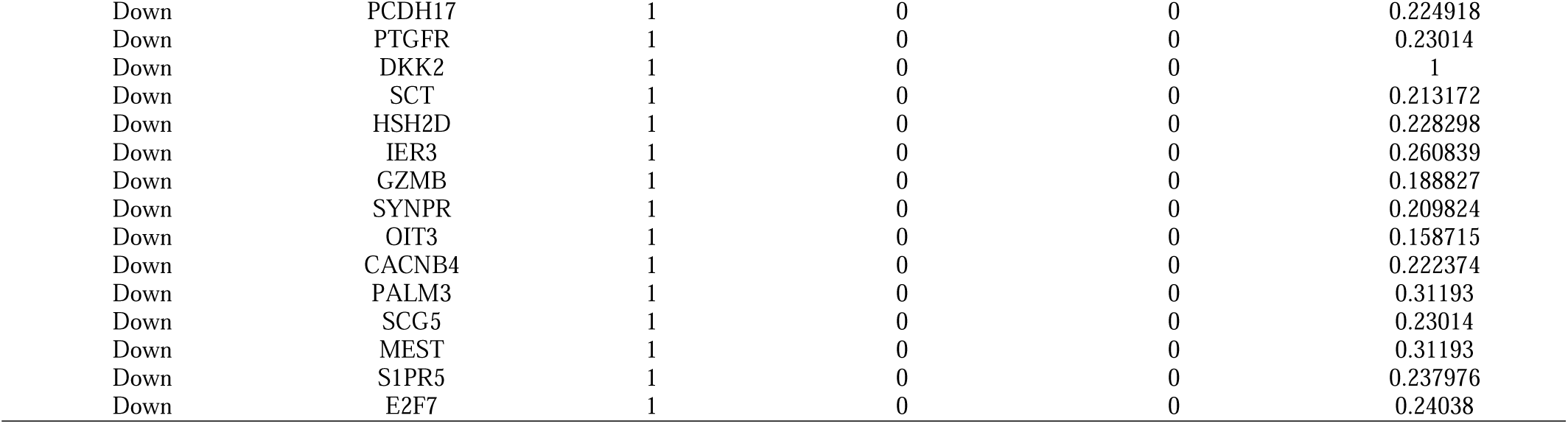
Topology table for up and down regulated genes.

### Construction of the miRNA -hub gene regulatory network

The related miRNA-hub gene regulatory network was shown in Fig. 11. The miRNA-hub gene regulatory network was composed of 2481 nodes [Hub gene:301, miRNA: 2180] and 28559 edges. In the miRNA-hub gene regulatory network, FN1 was regulated by different 439 miRNAs (ex: hsa-mir-374a-5p); TFRC was regulated by different 422 miRNAs (ex: hsa-miR-520d-5p); FKBP5 was regulated by different 305 miRNAs (ex: hsa-miR-34a-5p); FHL2 was regulated by different 291 miRNAs (ex: hsa-mir-4646-5p); RPLP0 was regulated by different 232 miRNAs (ex: hsa-mir-548aw); TPM3 was regulated by different 456 miRNAs (ex: hsa-miR-8052); ESR1 was regulated by different 266 miRNAs (ex: hsa-mir-302b-3p); CHRM3 was regulated by different 226 miRNAs (ex:hsa-miR-194-5p); TMEM30B was regulated by different 203 miRNAs (ex: hsa-mir-548i); PAK1 was regulated by different 162 miRNAs (ex: hsa-miR-524-5p) (Table 6).

**Fig. 11.**
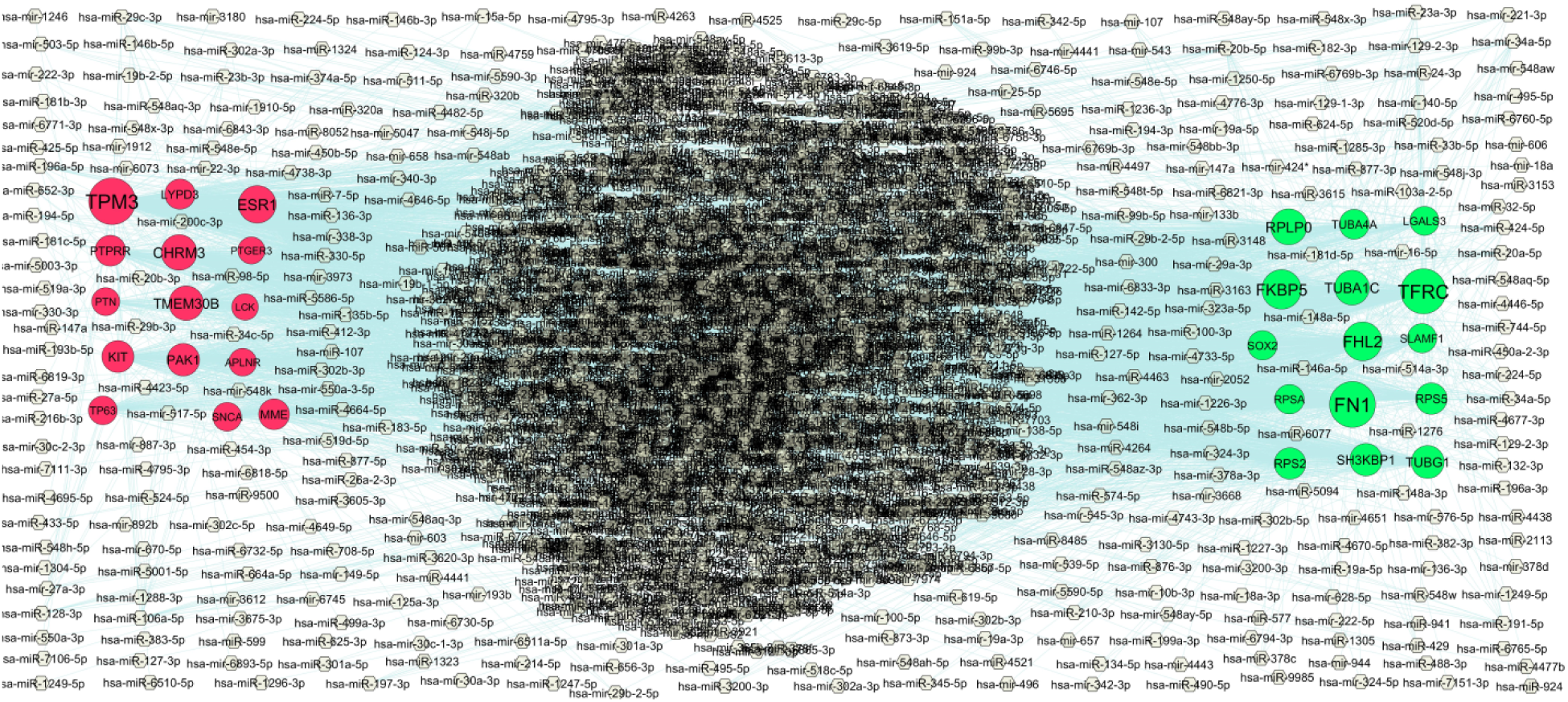
Hub gene - miRNA regulatory network. The light gray color diamond nodes represent the key miRNAs; up regulated genes are marked in green; down regulated genes are marked in red.

**Table 6.** MiRNA - hub gene and TF – hub gene topology table.

| Regulation | Hub Genes | Degree | MicroRNA | Regulation | Hub Genes | Degree | TF |
| --- | --- | --- | --- | --- | --- | --- | --- |
| Up | FN1 | 439 | hsa-mir-374a-5p | Up | SH3KBP1 | 20 | PRRX2 |
| Up | TFRC | 422 | hsa-miR-520d-5p | Up | FHL2 | 19 | SRY |
| Up | FKBP5 | 305 | hsa-miR-34a-5p | Up | FKBP5 | 17 | USF2 |
| Up | FHL2 | 291 | hsa-mir-4646-5p | Up | TFRC | 15 | BRCA1 |
| Up | RPLP0 | 232 | hsa-mir-548aw | Up | FN1 | 14 | STAT1 |
| Up | TUBA1C | 208 | hsa-mir-603 | Up | RPSA | 10 | PPARG |
| Up | TUBG1 | 170 | hsa-miR-148a-3p | Up | SOX2 | 9 | ARID3A |
| Up | SH3KBP1 | 159 | hsa-miR-135b-5p | Up | TUBA4A | 7 | TFAP2C |
| Up | RPS2 | 141 | hsa-mir-378a-3p | Up | TUBG1 | 6 | NFIC |
| Up | RPS5 | 139 | hsa-miR-182-3p | Up | RPLP0 | 6 | FOXF2 |
| Up | TUBA4A | 105 | hsa-mir-147a | Up | SLAMF1 | 6 | YY1 |
| Up | RPSA | 98 | hsa-miR-23a-3p | Up | TUBA1C | 5 | GATA3 |
| Up | SOX2 | 96 | hsa-miR-744-5p | Up | LGALS3 | 5 | NFATC2 |
| Up | SLAMF1 | 95 | hsa-miR-106a-5p | Up | RPS5 | 5 | RELA |
| Up | LGALS3 | 88 | hsa-miR-4525 | Up | RPS2 | 4 | SOX17 |
| Down | TPM3 | 456 | hsa-miR-8052 | Down | ESR1 | 26 | STAT3 |
| Down | ESR1 | 266 | hsa-mir-302b-3p | Down | TP63 | 21 | FOS |
| Down | CHRM3 | 226 | hsa-miR-194-5p | Down | PTPRR | 16 | MEF2A |
| Down | TMEM30B | 203 | hsa-mir-548i | Down | TPM3 | 13 | EN1 |
| Down | PAK1 | 162 | hsa-miR-524-5p | Down | SNCA | 11 | GATA2 |
| Down | KIT | 150 | hsa-miR-23a-3p | Down | LCK | 10 | FOXL1 |
| Down | PTPRR | 127 | hsa-mir-129-2-3p | Down | PTGER3 | 9 | POU2F2 |
| Down | MME | 126 | hsa-mir-181d-5p | Down | MME | 9 | NFYB |
| Down | LYPD3 | 115 | hsa-mir-27a-3p | Down | KIT | 8 | FOXC1 |
| Down | SNCA | 84 | hsa-miR-99b-5p | Down | PAK1 | 7 | REL |
| Down | TP63 | 82 | hsa-miR-1285-3p | Down | TMEM30B | 7 | PAX2 |
| Down | PTN | 66 | hsa-miR-320b | Down | LYPD3 | 6 | SRF |
| Down | APLNR | 61 | hsa-miR-146a-5p | Down | PTN | 6 | ELK4 |
| Down | PTGER3 | 49 | hsa-miR-5695 | Down | APLNR | 4 | TP53 |
| Down | LCK | 37 | hsa-miR-183-5p | Down | CHRM3 | 2 | HOXA5 |

### Construction of the TF-hub gene regulatory network

The related TF-hub gene regulatory network was shown in Fig. 12. The TF-hub gene regulatory network was composed of 382 [Hub gene:83, TF: 299] nodes and 2359 edges. In the TF-hub gene regulatory network, SH3KBP1 was regulated by 20 different TFs (ex: PRRX2); FHL2 was regulated by 19 different TFs (ex: SRY); FKBP5 was regulated by 17 different TFs (ex: USF2); TFRC was regulated by 15 different TFs (ex: BRCA1); FN1 was regulated by 14 different TFs (STAT1); ESR1 was regulated by 26 different TFs (ex: STAT3); TP63 was regulated by 21 different TFs (ex: FOS); PTPRR was regulated by 16 different TFs (ex: MEF2A); TPM3 was regulated by 13 different TFs (ex: MEF2A); SNCA was regulated by 11 different TFs (ex: GATA2) (Table 6).

**Fig. 12.**
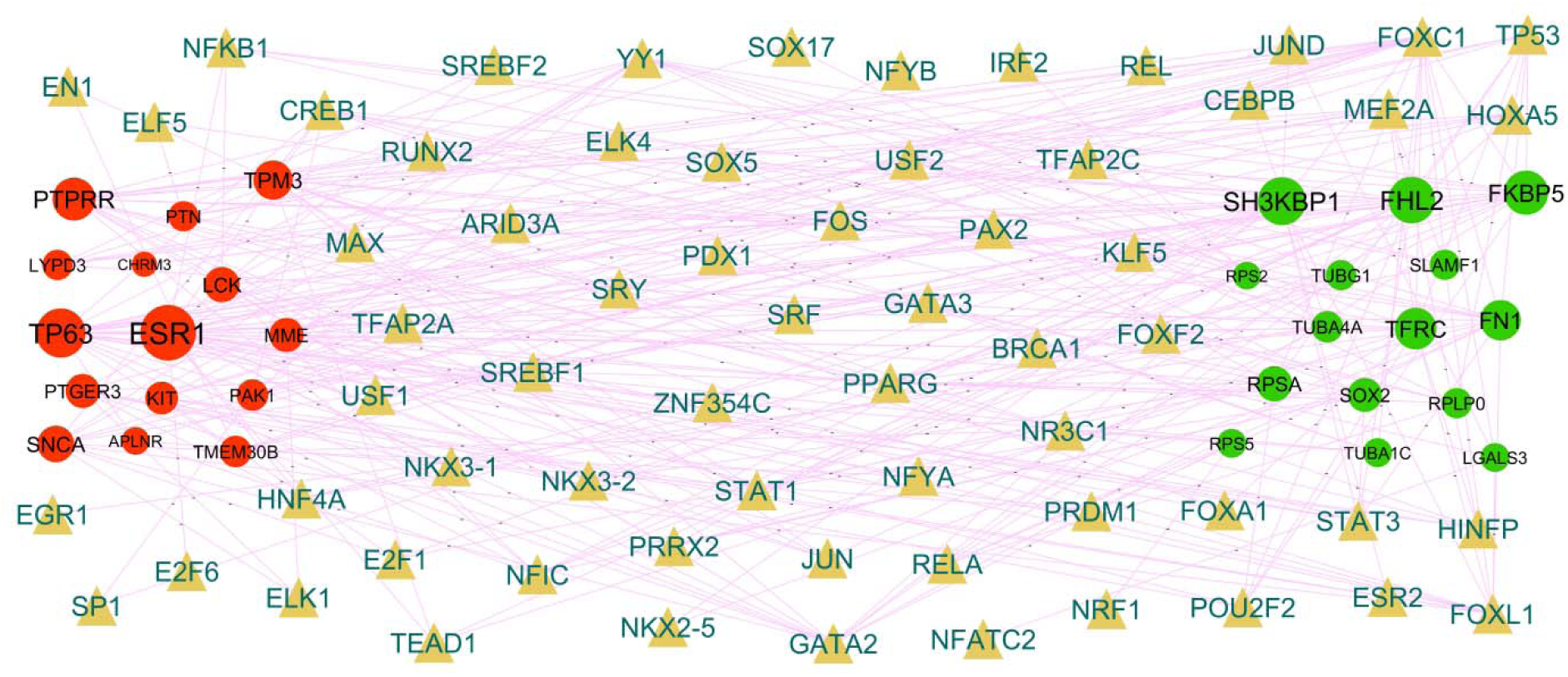
Hub gene - TF regulatory network. The goldenrod color triangle nodes represent the key TFs; up regulated genes are marked in dark green; down regulated genes are marked in dark red.

### Construction of the drug-hub gene interaction network

The related drug-hub gene interaction network was shown in Fig. 13 and Fig. 14. In the drug-hub gene interaction network, up regulated genes such as CHRNA4 was targeted by 50 different drug molecules (ex: Succinylcholine), GRIN2A was targeted by 34 different drug molecules (ex: Memantine), SCN1A was targeted by 27 different drug molecules (ex: Phenacemide), VDR was targeted by 22 different drug molecules (ex: Lexacalcitol) and CA14 was targeted by 21 different drug molecules (ex: Zonisamide) (Table 7), while down regulated genes such as CHRM3 was targeted by 94 different drug molecules (ex: Umeclidinium), GABRE was targeted by 76 different drug molecules (ex: Bromazepam), NOS2 was targeted by 53 different drug molecules (ex: Dipyrithione), KIT was targeted by 28 different drug molecules (ex: Midostaurin) and LCK was targeted by 26 different drug molecules (ex: Nintedanib) (Table 7).

**Fig. 13.**
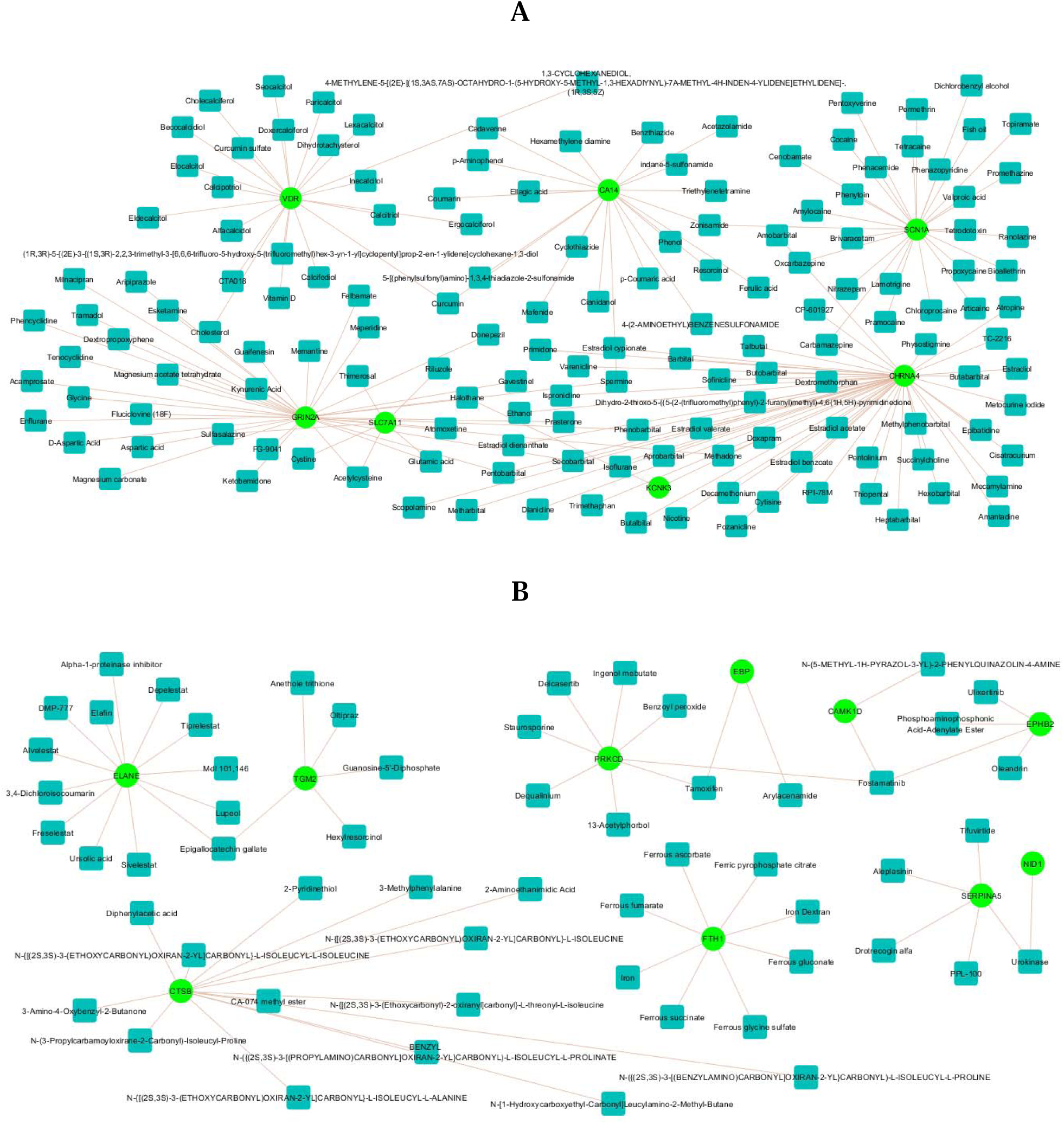
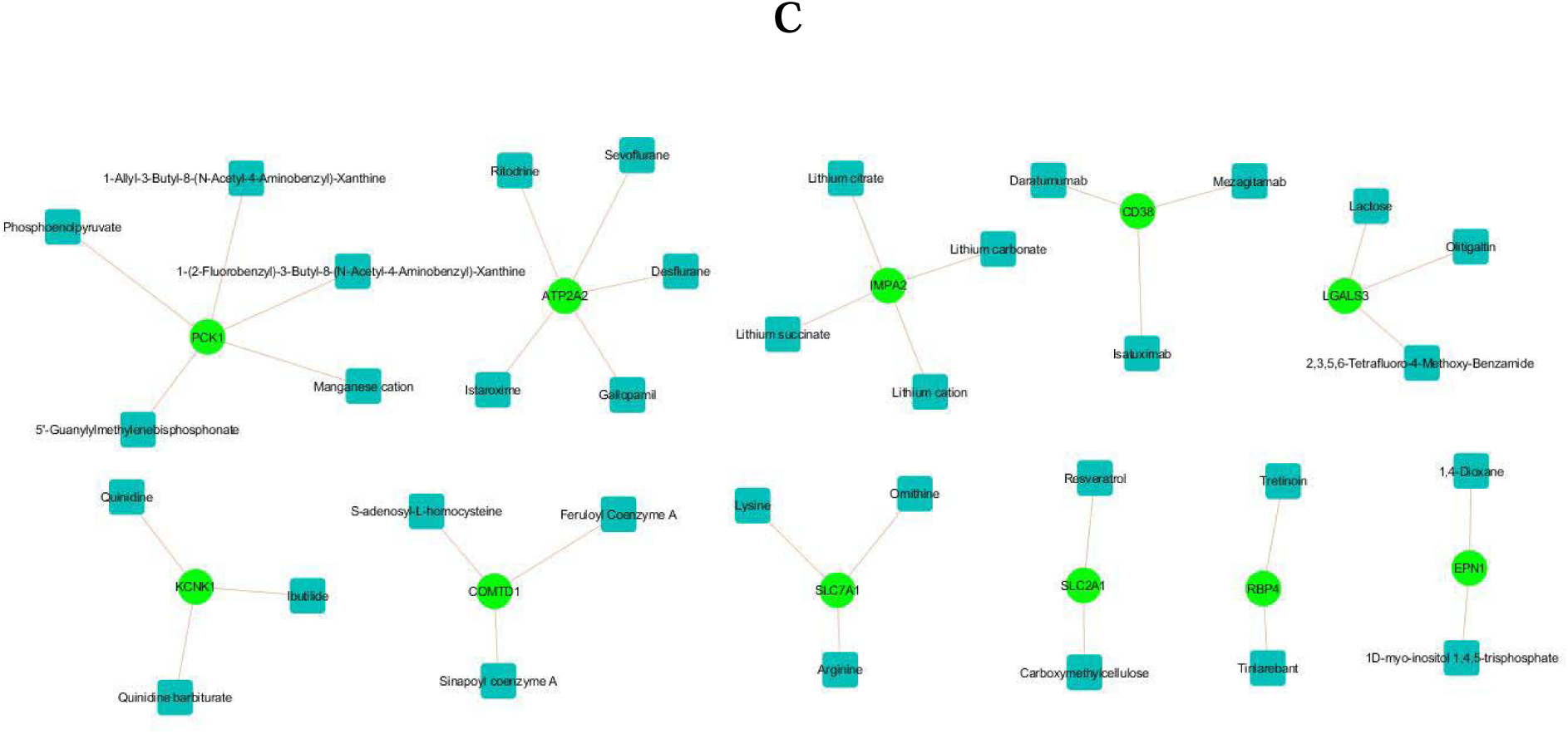
Three drug-hub gene interaction networks include 1) Network A 2) Network B 3) Network C 4) Network D. The blue rectangle nodes represent the drug molecule; up regulated genes are marked in green

**Fig. 14.**
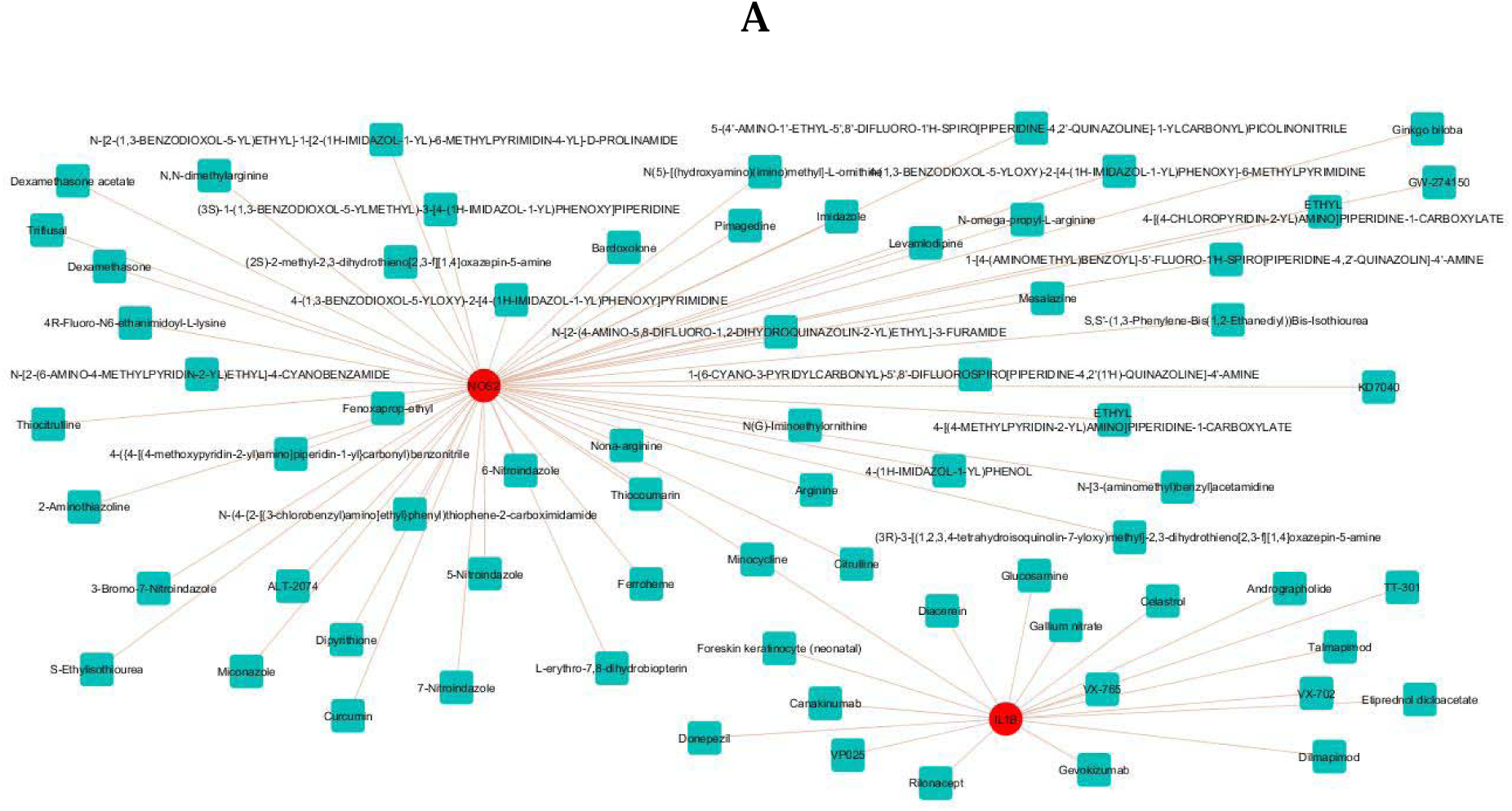
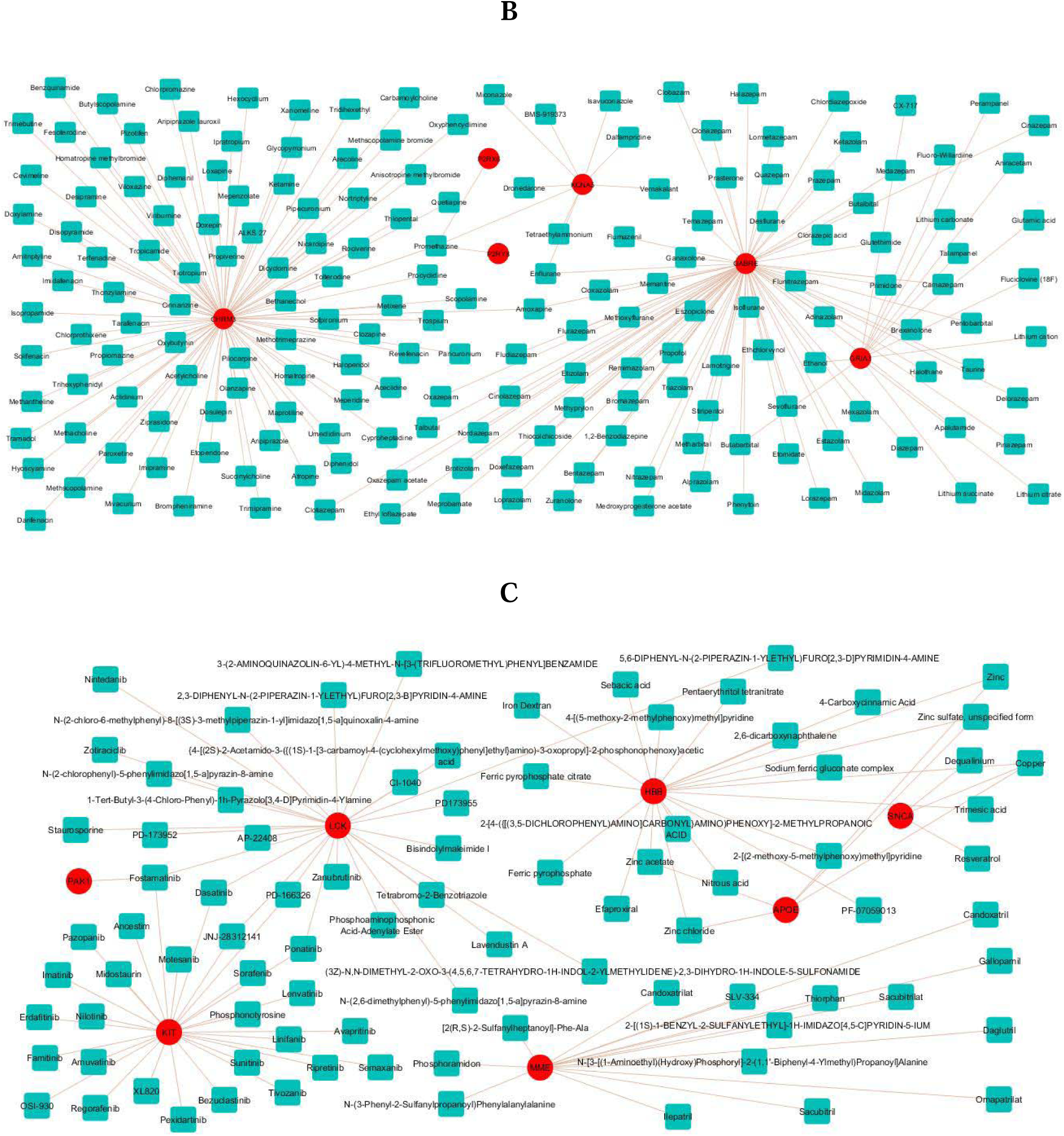
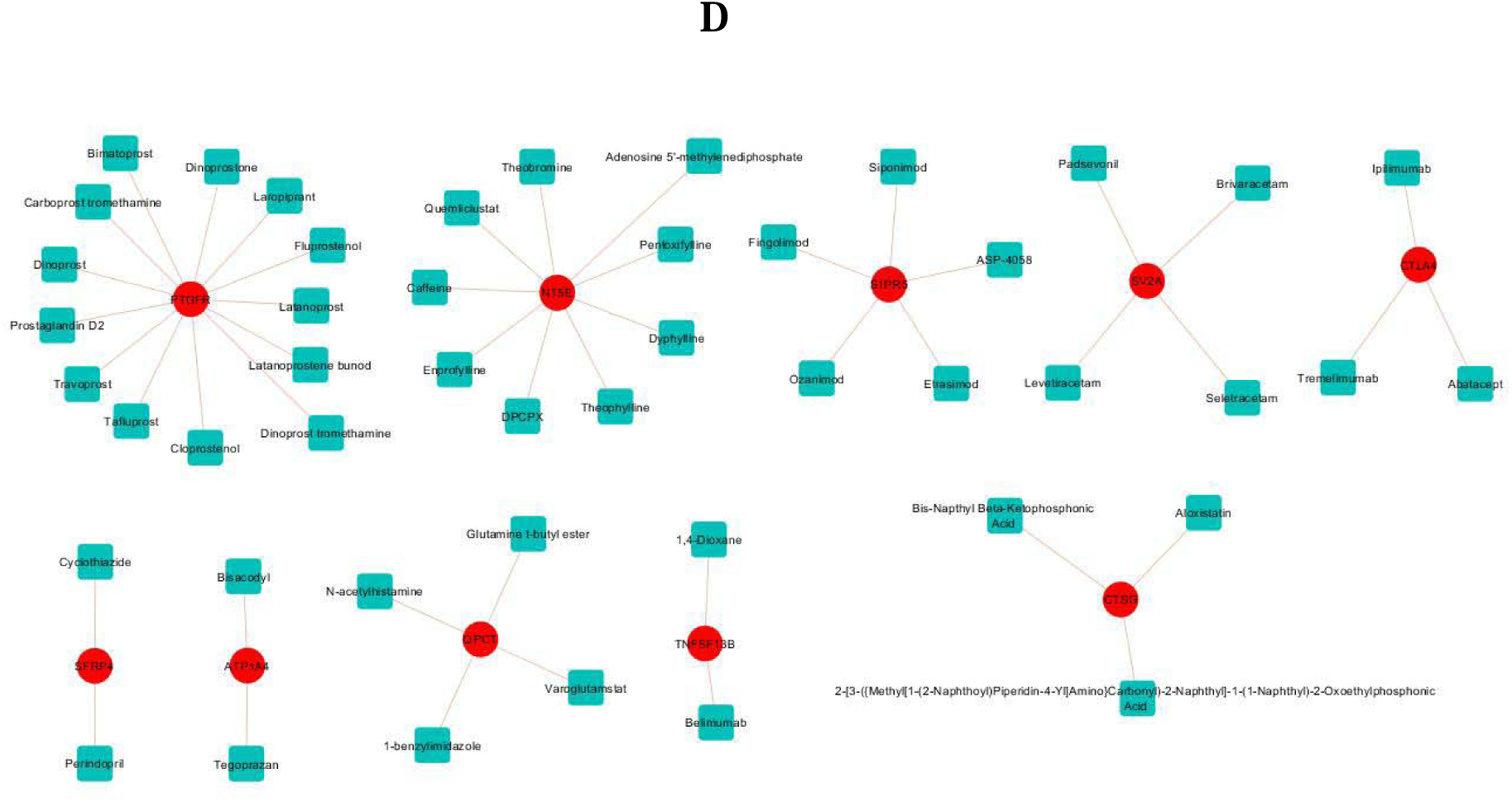
Four drug-hub gene interaction networks include 1) Network A 2) Network B 3) Network C 4) Network D. The blue rectangle nodes represent the drug molecule; down regulated genes are marked in red

**Table 7.**
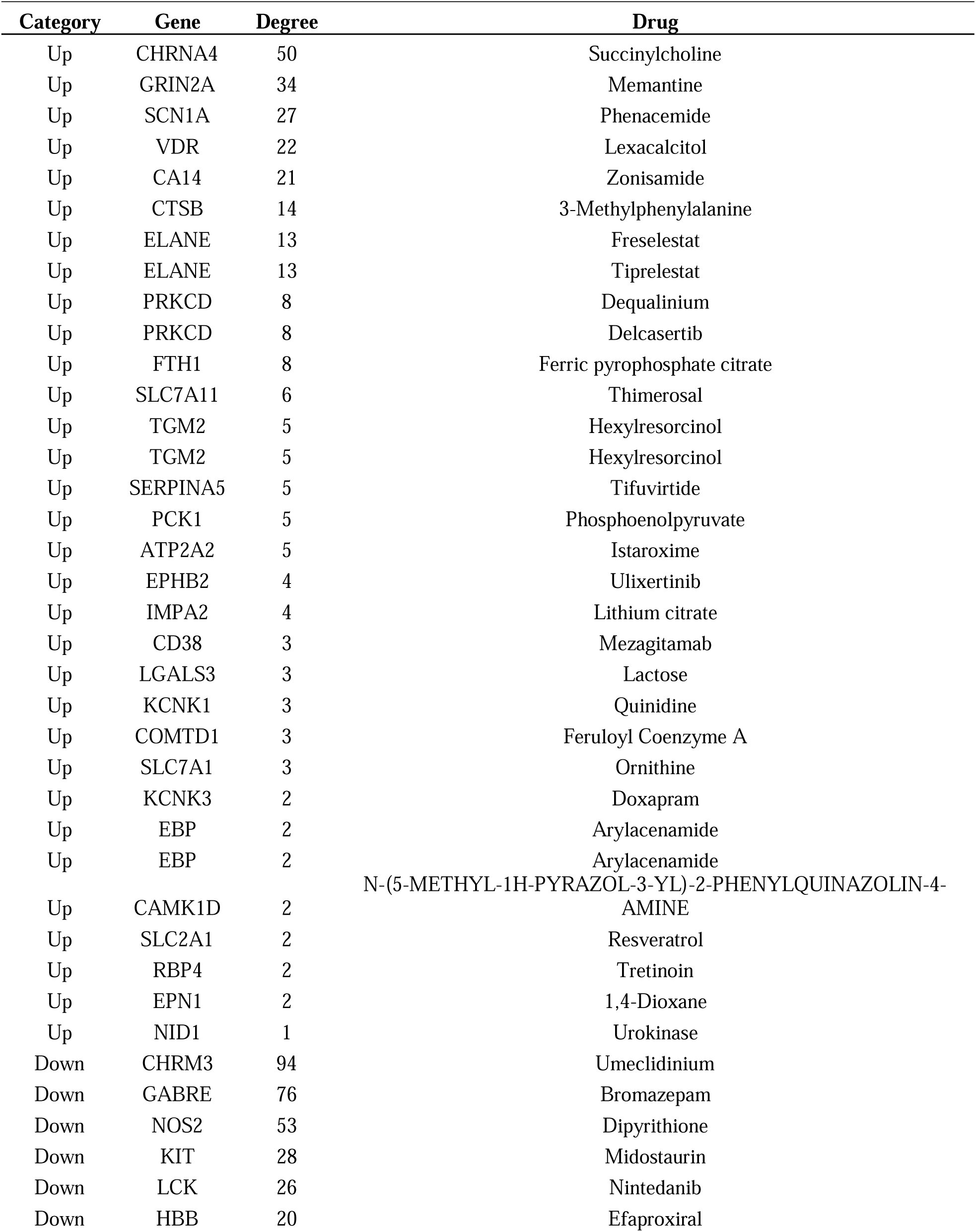
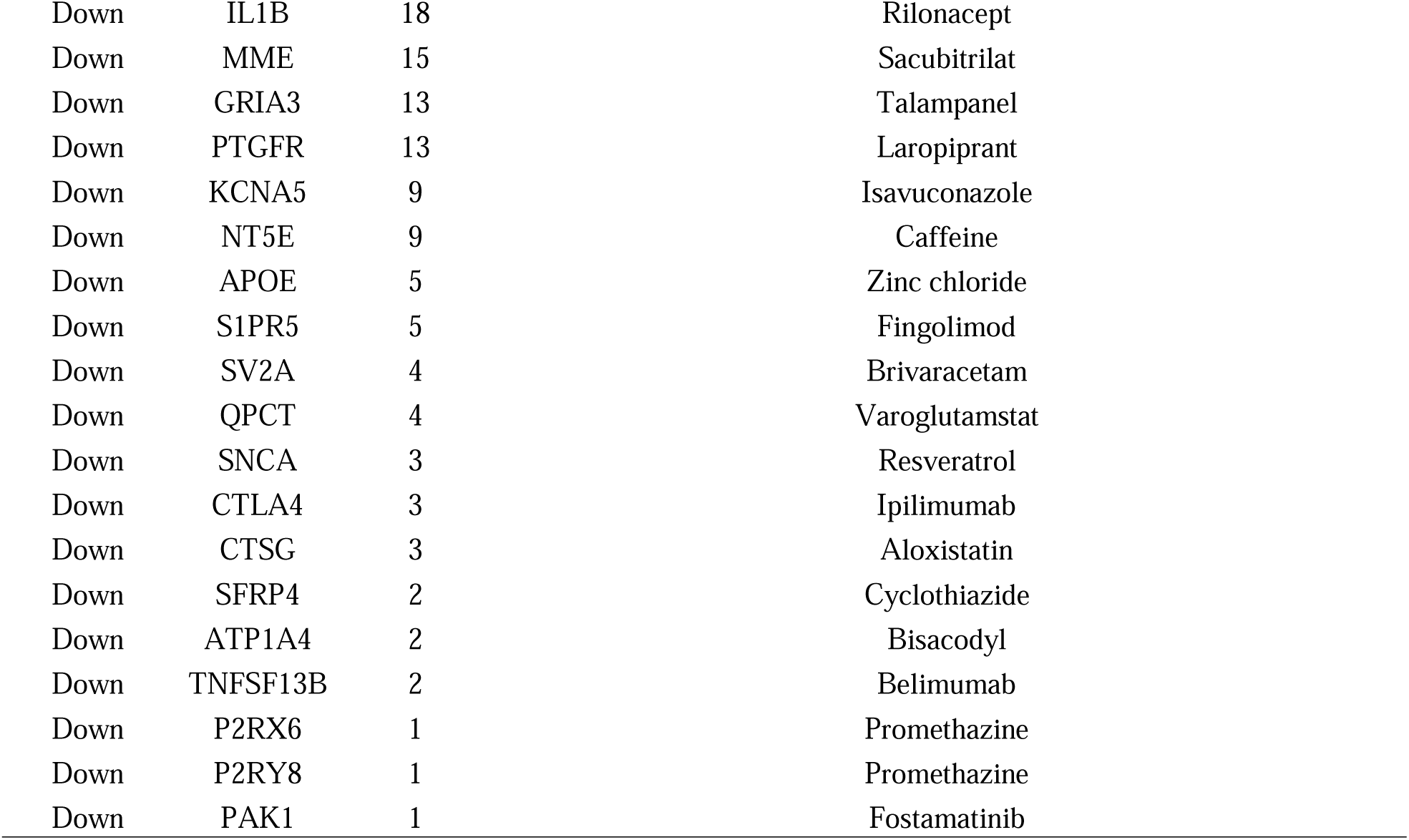
Drug- hub gene topology table.

### Receiver operating characteristic curve (ROC) analysis

The ROC curve was utilized to assess the diagnostic efficacy of hub genes in predicting HCM occurrence and advancement. A greater AUC value demonstrate better predictive efficacy. The AUC values of the hub genes were FN1-AUC:0.934, SOX2-AUC:0.919, TUBA4A-AUC:0.923, RPS2-AUC:0.891, TUBA1C-AUC:0.908, ESR1-AUC:0.927, SNCA-AUC:0.900, LCK-AUC:0.938, PAK1-AUC:0.895 and APLNR-AUC:0.897 suggesting that these hub genes all display remarkable predictive ability for obesity occurrence and development (Fig. 15).

**Fig. 15.**
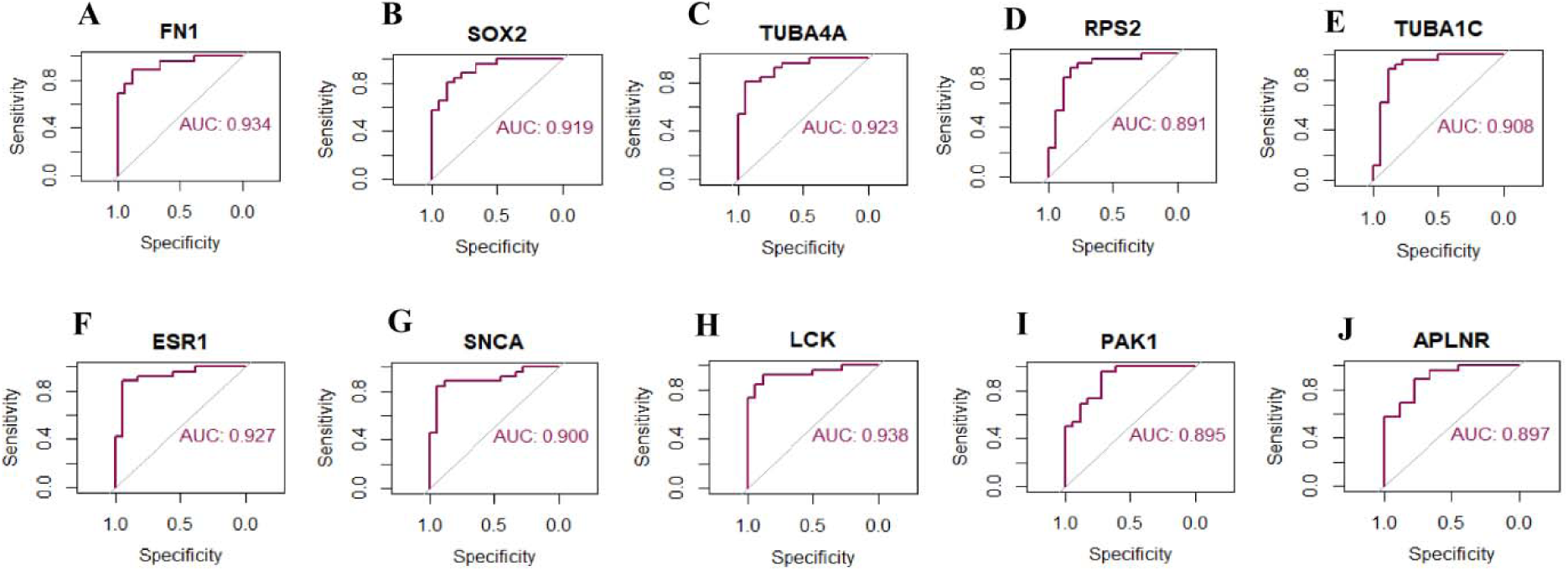
ROC curve analyses of hub genes. A) FN1 B) SOX2 C) TUBA4A D) RPS2 E) TUBA1C F) ESR1G) SNCA H) LCK I) PAK1 J) APLNR

### *Insilico* molecular docking studies

#### Analysis of molecular docking with LGALS3

A molecular docking investigation was conducted to evaluate the binding behavior of various phytoconstituents inside the carbohydrate recognition domain (CRD) of LGALS3, an elevated mediator of fibrosis and inflammatory remodeling in hypertrophic cardiomyopathy. All tested compounds fit effectively into the expected active pocket, suggesting structural compatibility with the LGALS3 binding cavity.

The reference co-crystallized ligand QB2 has a binding affinity of -6.565 kcal/mol and interacts with critical residues involved in galectin-3 carbohydrate recognition, making it a reliable baseline for comparison study. Notably, numerous phytochemicals had binding affinities comparable to or greater than that of the reference ligand (Table 8). Puerarin had the highest binding affinity (-6.807 kcal/mol) of all chemicals tested. Puerarin formed stable hydrogen bond interactions with GLU72 and ARG50, which are important residues for ligand anchoring within the CRD. In addition, pi-pi stacking interaction with TRP69, a residue often described as crucial for galectin-3 ligand stability, strengthened the binding pose. Hydrophobic interactions with VAL34 led to pocket stability, indicating a good mix of polar and non-polar interactions.Oleuropein had a high binding affinity (-6.423 kcal/mol) and established several hydrogen bonds with ARG50, HIS46, ASN62, and ASN67, showing extensive polar interaction networks in the CRD. The presence of pi-pi interaction with TRP69 strengthens its stable accommodation in the active pocket. Oleuropein’s many interactions suggest that it may effectively compete with endogenous ligands implicated in galectin-3-mediated fibrotic signaling.Swertiamarin and Mangiferin showed moderate but constant binding affinities. Swertiamarin particularly targeted residues like ARG32, ASP36, and HIS46, generating hydrogen bonds that kept the ligand in the binding groove. Mangiferin formed hydrogen connections with LYS64 and ASN67, as well as hydrophobic contacts with VAL34, demonstrating that pocket occupancy (Fig. 16 and Fig. 17) was sustained despite the significantly lower docking scores.

**Fig. 16.**
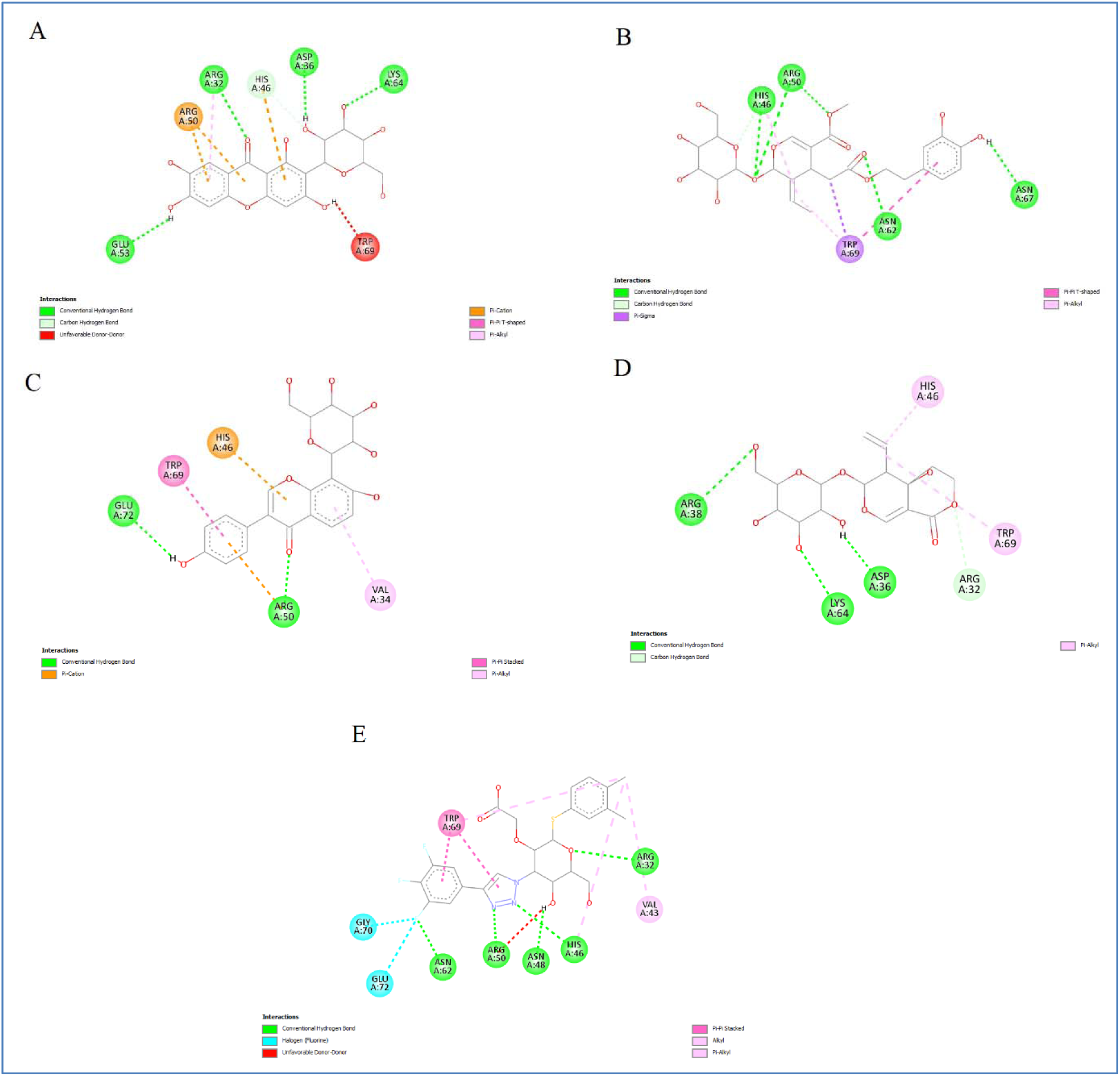
Two-dimensional interaction maps of docked phytoconstituents within the LGALS3 binding pocket (PDB ID: 8IU1). A. Mangiferin, B. Oleuropein, C. Puerarin, D. Swertiamerin and E. cocrystallised ligand (QB2).

**Fig. 17.**
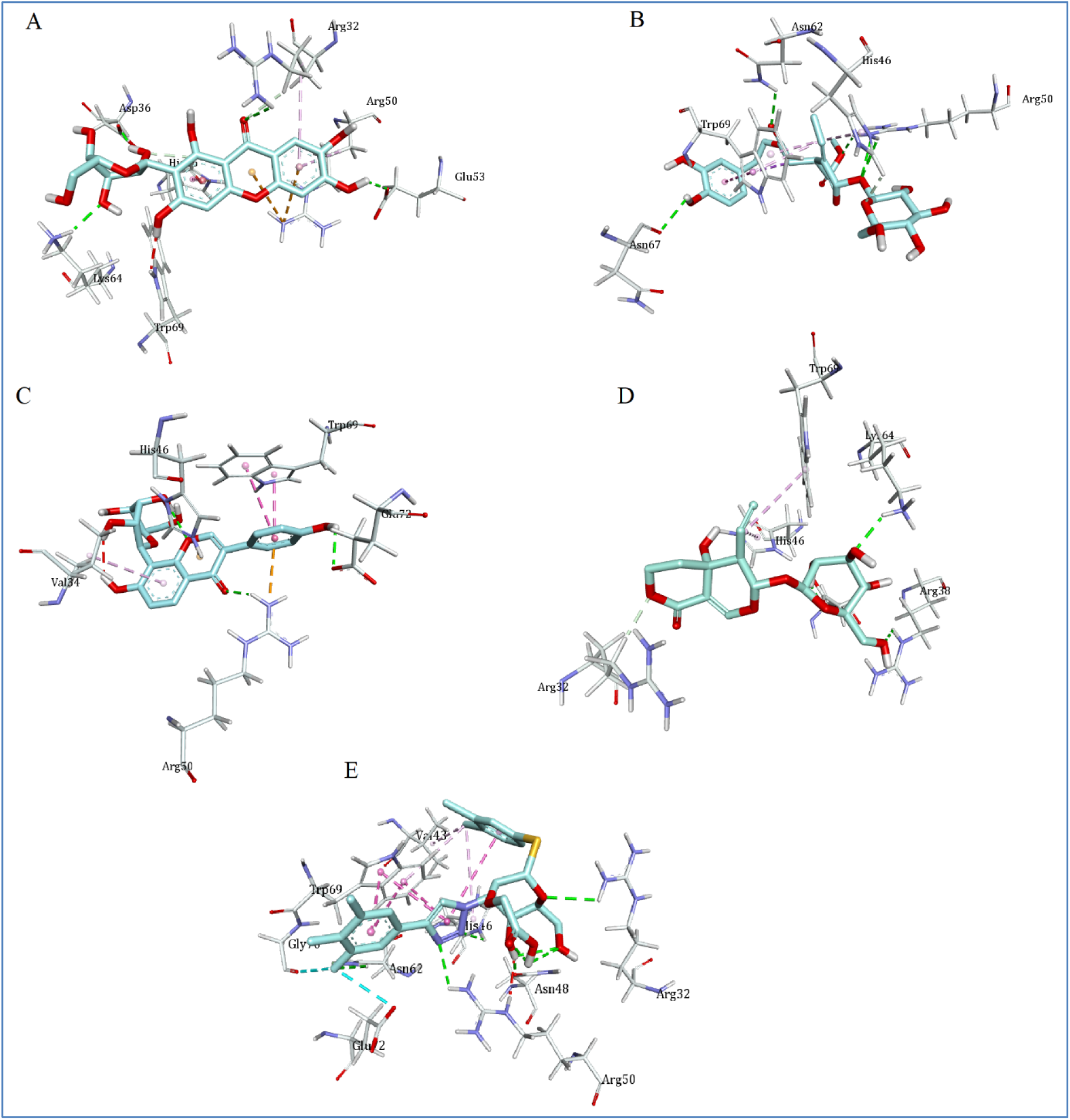
Three-dimensional binding conformations of screened phytoconstituents at the LGALS3 active site (PDB ID: 8IU1). A. Mangiferin, B. Oleuropein, C. Puerarin, D. Swertiamerin and E. cocrystallised ligand (QB2).

**Table 8.**
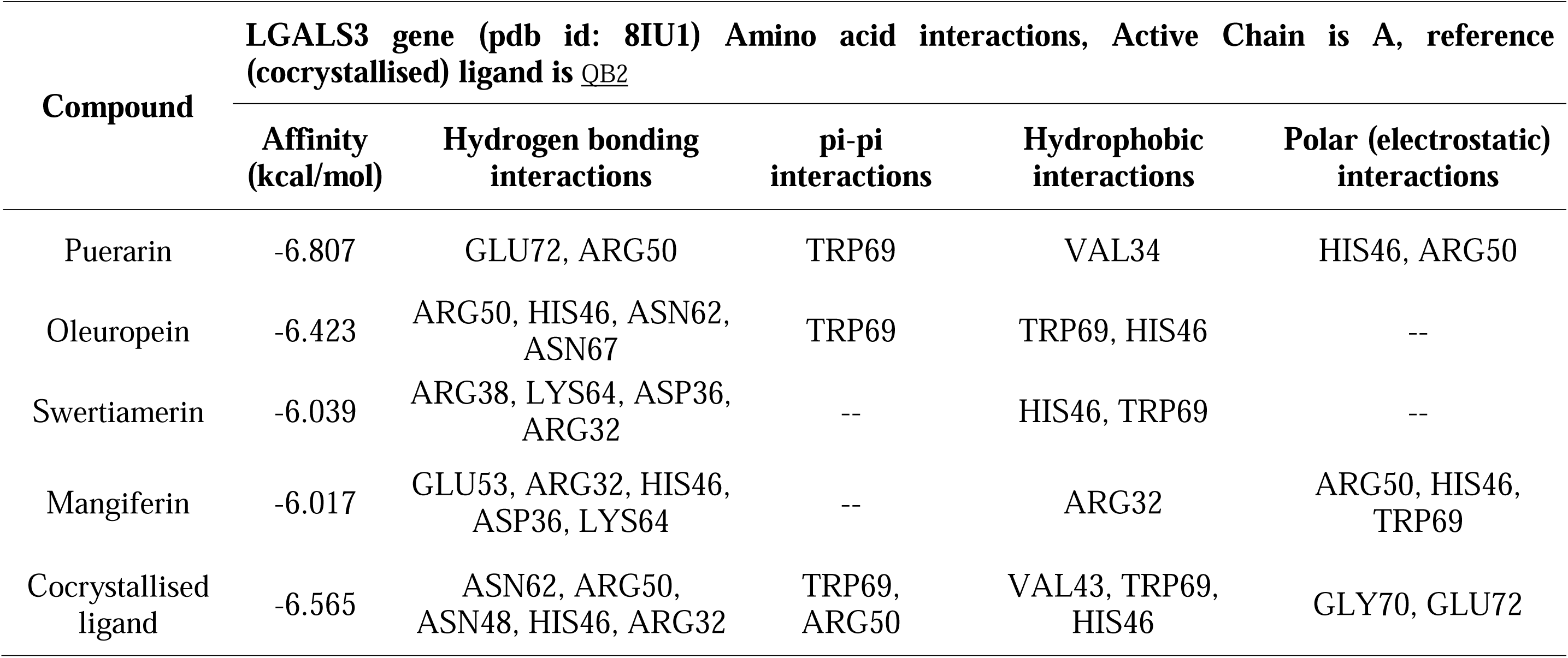
Molecular dockinginteractions of screened phytoconstituents with LGALS3 (PDB ID: 8IU1)

| Compound | LGALS3 gene (pdb id: 8IU1) Amino acid interactions, Active Chain is A, reference (cocrystallised) ligand is <u>QB2</u> |  |  |  |  |
| --- | --- | --- | --- | --- | --- |
|  | Affinity (kcal/mol) | Hydrogen bonding interactions | pi-pi interactions | Hydrophobic interactions | Polar (electrostatic) interactions |
| Puerarin | -6.807 | GLU72, ARG50 | TRP69 | VAL34 | HIS46, ARG50 |
| Oleuropein | -6.423 | ARG50, HIS46, ASN62, ASN67 | TRP69 | TRP69, HIS46 | -- |
| Swertiamerin | -6.039 | ARG38, LYS64, ASP36, ARG32 | -- | HIS46, TRP69 | -- |
| Mangiferin | -6.017 | GLU53, ARG32, HIS46, ASP36, LYS64 | -- | ARG32 | ARG50, HIS46, TRP69 |
| Cocrystallised ligand | -6.565 | ASN62, ARG50, ASN48, HIS46, ARG32 | TRP69, ARG50 | VAL43, TRP69, HIS46 | GLY70, GLU72 |

#### Molecular docking analysis of ESR1

Docking tests were carried out against ESR1 (PDB ID: 1A52), which reflects the receptor’s agonist-bound active conformation, to investigate the potential restoration of cardioprotective estrogensignaling. ESR1 was found to be a downregulated gene in hypertrophic cardiomyopathy, which justifies further research into ligand interactions inside the agonist-binding pocket.

The co-crystallized ligand 17β-estradiol has the highest binding affinity (-10.754 kcal/mol), establishing hydrogen bonds with residues that activate receptors and stabilize helix-12. This binding mode served as a standard for evaluating phytochemical docking scores (Table. 9). Oleuropein has the highest binding affinity (-7.745 kcal/mol) of all the substances tested. Oleuropein formed several hydrogen bonds with GLY217, ASP47, LYS225, LEU221, HIS220, and ALA46, demonstrating a wide range of polar interactions within the ligand-binding cavity. Furthermore, π-π stacking interaction with PHE100 and hydrophobic interactions with LEU83 and LEU87 enhanced overall complex stability. These interactions indicate that oleuropein effectively fills the estradiol-binding pocket, potentially supporting receptor stability in an agonist-like orientation.Puerarin had a moderate binding affinity (-6.559 kcal/mol) and interacted with GLU49, THR43, and LYS225 via hydrogen bonding, as well as hydrophobic interactions with LEU221 and ALA46. These interactions overlap with residues involved in agonist stabilization, implying a potential modulatory effect.Swertiamarin and Mangiferin exhibited reduced binding affinities but remained stable within the agonist-binding cavity. Their interaction patterns were dominated by hydrogen bonds with polar residues and hydrophobic interactions with aliphatic amino acids bordering the pocket (Fig. 18 and Fig. 19), indicating ligand accommodation without causing adverse steric conflicts.

**Fig. 18.**
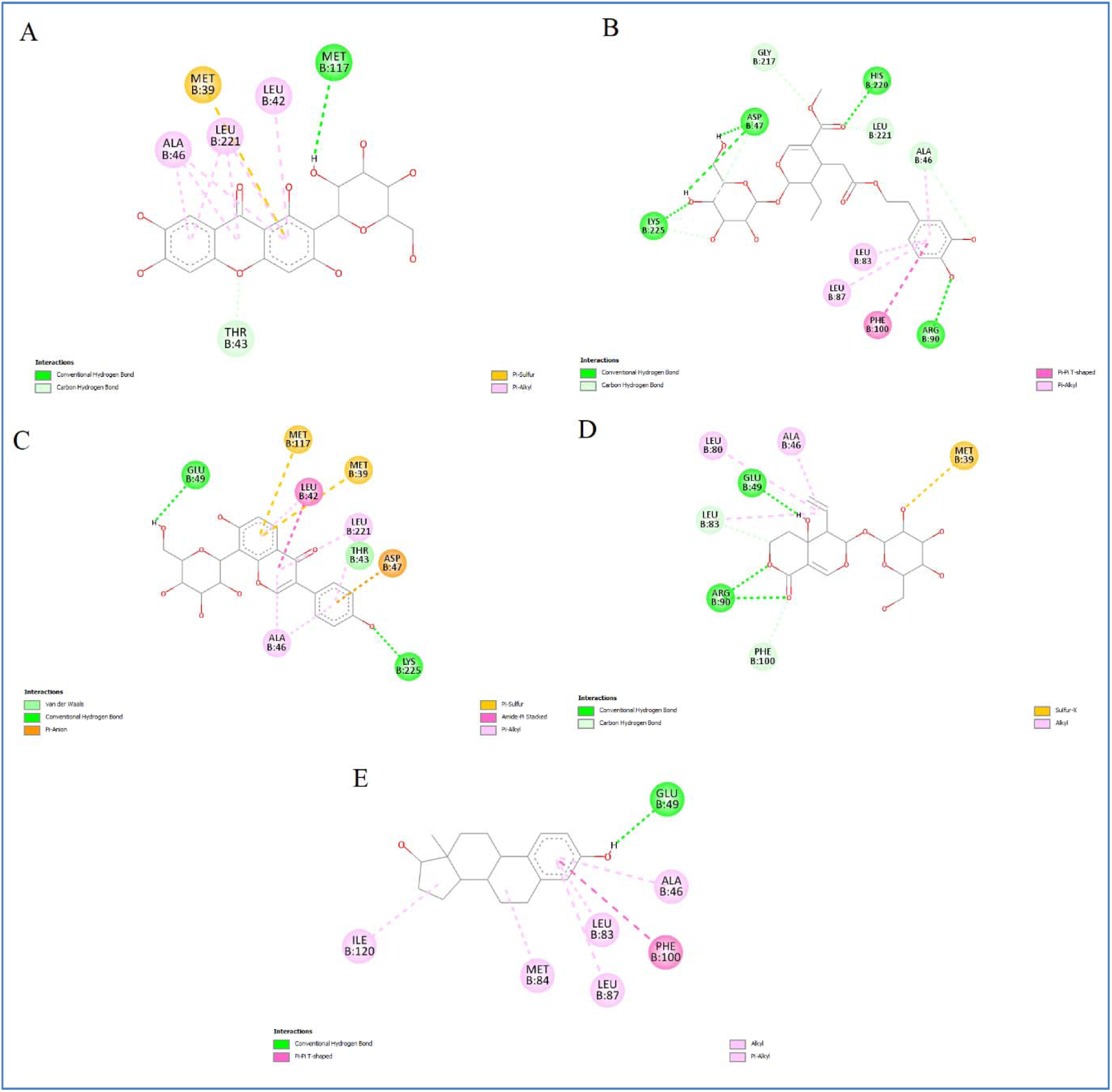
Two-dimensional interaction maps of docked phytoconstituents within the ESR1 binding pocket (PDB ID: 1A52). A. Mangiferin, B. Oleuropein, C. Puerarin, D. Swertiamerin and E. cocrystallised ligand (Estradiol).

**Fig. 19.**
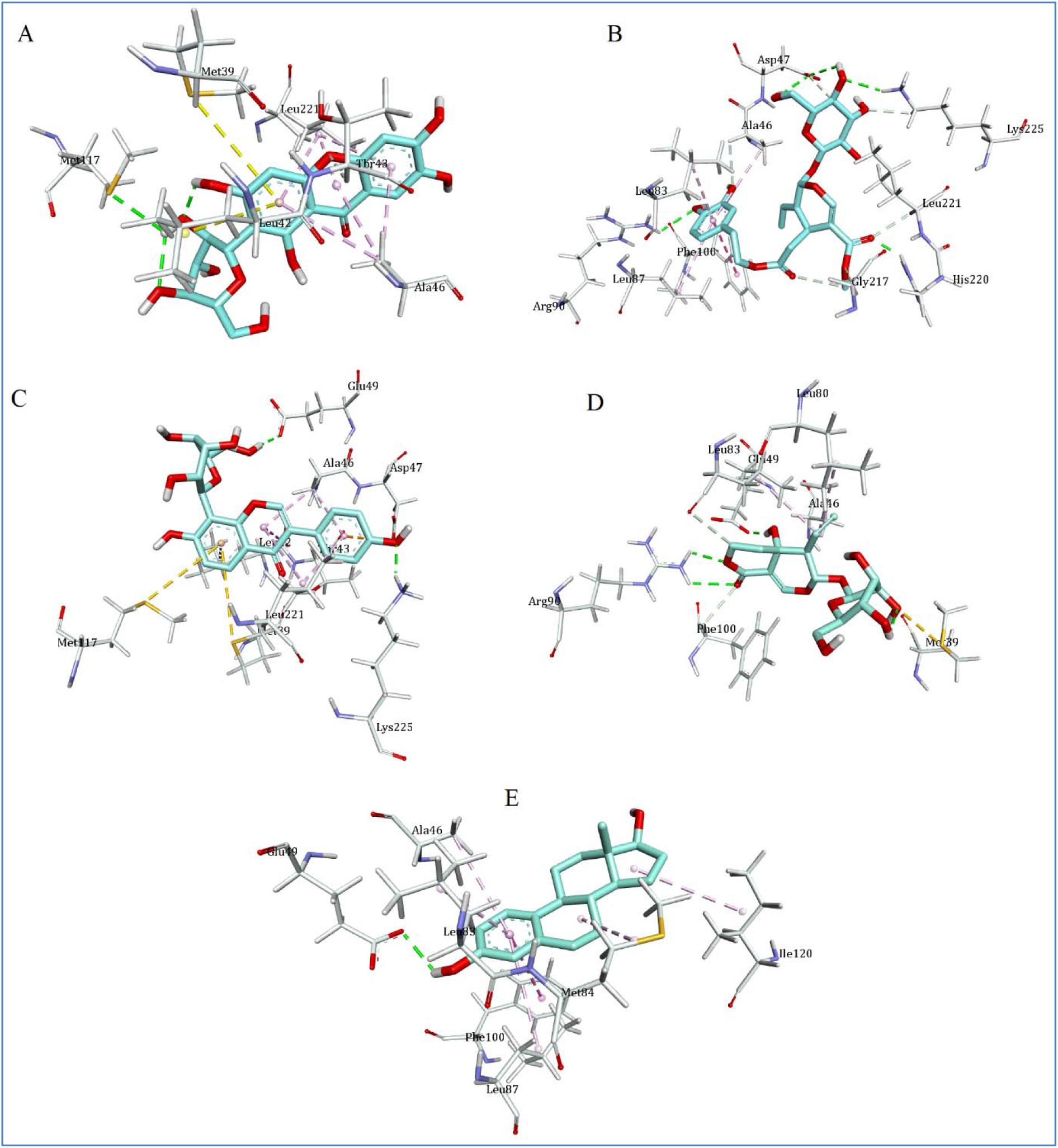
Three-dimensional binding conformations of screened phytoconstituents at the LGALS3 active site (PDB ID: 1A52). A. Mangiferin, B. Oleuropein, C. Puerarin, D. Swertiamerin and E. cocrystallised ligand (Estradiol).

#### Drug-likeness properties evaluation

The drug-likeness assessment found that Swertiamarin and Puerarin met Lipinski’s rule of five, indicating desirable physicochemical qualities for oral bioavailability. Oleuropein and Mangiferin, on the other hand, did not meet Lipinski’s criterion, owing to their larger molecular weight and increased polar surface area, which are typical of polyphenolic phytoconstituents.Despite these violations, all compounds had acceptable rotatable bond numbers and polarity ranges, indicating that aberrations may not rule out biological activity, especially in the context of natural product-based therapies.

#### ADMET properties analysis

ADMET profiling (Table.10) revealed that all investigated phytoconstituents had appropriate aqueous solubility, as measured by ESOL Log S values. All substances were anticipated to have high human intestinal absorption, indicating a favorable oral absorption potential.Caco-2 permeability values ranged from low to moderate, which is consistent with the molecules’ polar nature. All drugs demonstrated minimal BBB permeability, lowering the possibility of central nervous system-related side effects, which is beneficial for cardiovascular therapies.According to P-glycoprotein substrate prediction, Puerarin, Oleuropein, and Mangiferin may be P-gp substrates, but Swertiamarin has lower P-gp liability. Importantly, all substances showed modest inhibitory activity against CYP2C9 and CYP3A4, indicating a lower chance of drug-drug interactions. Cardiotoxicity prediction found modest hERG inhibition probability, while carcinogenicity prediction was limited across all drugs, indicating an overall positive safety profile.

## Discussion

HCM is a cardiovascular disorder with high rates of pervasiveness and mortality [Siontis et al. 2014]. Therefore, sensitive and specific biomarkers of HCM are urgently needed to be detected. In the current investigations, bioinformatic and NGS methods are promising methods to analyze the key genes and signaling pathways, which might contribute new indication for diagnosis, therapy, and prognosis of HCM.

In this investigation, we analyzed the HCM NGS data GSE146638 screened from the GEO database. It includes 7 normal control samples and 13 HCM samples. We identified 958 DEGs, including 479 up-regulated genes and 479 down-regulated genes. SAA1 [Pan and Zhang, 2020], MYH6 [Yang et al. 2024] and MSTN (myostatin) [Qi et al. 2020] are mainly associated with HCM. SAA1 [Pan and Zhang, 2020], MYH6 [Zhao et al. 2021] and MSTN (myostatin) [Mahmoudabady et al. 2008] plays a role in diagnosis of dilated cardiomyopathy. SAA1 [Xiao et al. 2023], SERPINA3 [Delrue et al. 2021], MYH6 [Anfinson et al. 2022], ESM1 [Watanabe et al. 2021] and MSTN (myostatin) [Wang et al. 2023] served as a biomarkers for heart failure diagnosis and prognosis. SAA1 [Wang et al. 2014], SERPINA3 [Zhao et al. 2020], ESM1 [Watanabe et al. 2021] and MSTN (myostatin) [Lim et al. 2018] have been known to be involved in myocardial infarction progression. SAA1 [Yeh et al. 2022], SERPINA3 [Li et al. 2021], IL1RL1 [Lin et al. 2017], ESM1 [Sun et al. 2019] and MSTN (myostatin) [Dschietzig, 2014] have a significant prognostic potential in coronary artery disease. SAA1 [Leow et al. 2013] and MSTN (myostatin) [Verzola et al. 2017] are molecular markers for the diagnosis and prognosis of atherosclerosis. Studies had shown that SAA1 [Xiao et al. 2023] and MSTN (myostatin) [Biesemann et al. 2015] were associated with cardiac fibrosis. Studies have found that SAA1 [Bich et al. 2022], RNASE2 [Ong et al. 2023], SERPINA3 [Soman et al. 2022], IL1RL1 [Yu et al. 2020], ESM1 [Zeng et al. 2021] and MSTN (myostatin) [Verzola et al. 2020] are altered expressed in inflammation. The altered expression of SAA1 [Alharbi et al. 2021], IL1RL1 [Yu et al. 2020], ESM1 [Janke et al. 2006] and MSTN (myostatin) [Amor et al. 2019] might indicate increased risk of obesity. SAA1 [Wang et al. 2023], PTPRN (protein tyrosine phosphatase receptor type N) [Garranzo-Asensio et al. 2022], MYH6 [Trombetta et al. 2012], ESM1 [Qiu et al. 2017] and MSTN (myostatin) [Amor et al. 2019] plays an indispensable role in diabetes mellitus. MYH6 [Fu et al. 2021] and CUX2 [Wang et al. 2018] could be an early detection markers for atrial fibrillation. MYH6 [Zhao et al. 2021] plays a significant role in cardiac death. Study suggested that MYH6 [Lam et al. 2015] as potential diagnostic biomarker for cardiac arrest. Recently, mounting investigation has revealed that ESM1 [Hong et al. 2022] was vital for the onset and developmental process of mitral valve disease. MSTN (myostatin) [Castillero et al. 2015] has been demonstrated to be involved in myocardial ischemia. These findings suggest that significant DEGs can serve as candidate diagnostic biomarkers or drug targets for the clinical treatment of HCM.

g:Profiler online tool was used to predict the significantly enriched GO and REACTOME pathways. It has been confirmed that the pathogenesis of HCM is related to signaling pathways include metabolism [Timmer and Knaapen, 2013], cardiac conduction [Fitzgerald and Kusumoto, 2018], muscle contraction [Harada and Potter, 2004], neutrophil degranulation [Riascos-Bernal and Sibinga, 2022], metabolism of lipids [Frey and Olson, 2002], extracellular matrix organization [Viola et al. 2023], GPCR ligand binding [Liu et al. 2019], signaling by receptor tyrosine kinases [Sadoshima et al. 1995] and signal transduction [Jamshidi et al. 2004]. The expression of CD177 [Yang et al. 2019], ALOX15B [Magnusson et al. 2012], EREG (epiregulin) [Harada et al. 2015], FCN3 [Plovsing et al. 2016], SSTR5 [Liu et al. 2022], SFRP5 [Sun et al. 2021], SLAMF1 [Wang et al. 2015], CD163 [Etzerodt et al. 2014], AQP4 [Tourdias et al. 2011], PCK1 [Ko et al. 2018], VDR (vitamin D receptor) [Garg et al. 2019], SERPINE1 [Chen et al. 2021], AZGP1 [Na et al. 2017], VSIG4 [Liu et al. 2023], ELANE (elastase, neutrophil expressed) [Thanarajasingam et al. 2021], FKBP5 [Liu et al. 2023], MFSD2A [Ungaro et al. 2017], RBP4 [Pazos-Pérez et al. 2024], HMGCS2 [Chen et al. 2022], MOG (myelin oligodendrocyte glycoprotein) [Lalive et al. 2006], C3 [Jia et al. 2015], FGFR4 [Czaya et al. 2022], ITLN1 [Kerr et al. 2014], IL18R1 [Liu et al. 2012], TIMP4 [Koskivirta et al. 2006], LYVE1 [Johnson et al. 2007], GRIP1 [Miao et al. 2016], IL20RA [Lamichhane et al. 2021], HCRTR1 [Dong et al. 2024], LGR6 [Krishnamoorthy et al. 2023], SOX2 [Njouendou et al. 2023], TWIST2 [Ding et al. 2022], NPPC (natriuretic peptide C) [Bükülmez et al. 2014], SPP1 [Wu et al. 2024], CD109 [Aono et al. 2023], S1PR3 [Tian et al. 2024], BMP7 [Smoljan et al. 2023], SLC7A11 [Lv et al. 2024], MLXIPL (MLX interacting protein like) [Guo et al. 2023], AGTR1 [Musso et al. 2019], PTX3 [Ramirez et al. 2019], EPHB2 [Ernst et al. 2019], SLC2A1 [Petakh et al. 2024], TGM2 [Chang et al. 2024], FHL2 [van de Pol et al. 2020], CD38 [Zhang et al. 2024], FGF7 [Geervliet et al. 2023]. OSR1 [Liu et al. 2023], STXBP2 [Fujikawa et al. 2023], CRLF1 [Lu et al. 2024], POMC (proopiomelanocortin) [Ito et al. 2013], APOB (apolipoprotein B) [Webb et al. 2022], OLFM4 [Ren et al. 2021], BMAL1 [Chen et al. 2023], HEY2 [Lina et al. 2019], ARG2 [Li et al. 2019], LPCAT3 [Hu et al. 2007], IL15RA [Meghnem et al. 2022], FABP4 [Ma et al. 2024], PTGS1 [Plaza-Díaz et al. 2017], C1QTNF1 [Kim and Park, 2019], LMCD1 [Bai et al. 2023], TPO (thyroid peroxidase) [Nielsen et al. 2009], LOX (lysyl oxidase) [Huang et al. 2021], ADAMTS5 [Sharma et al. 2020], CD63 [Yu et al. 2021], HAS2 [Kim et al. 2022], NQO1 [Mondal et al. 2018], SIGLEC1 [Boz et al. 2024], NGFR (nerve growth factor receptor) [Zhao et al. 2024], PRDX6 [Yang et al. 2022], DHCR24 [Martiskainen et al. 2017], PDPN (podoplanin) [Quintanilla et al. 2019], AKR1C1 [Wang et al. 2007], CTSB (cathepsin B) [Fang et al. 2020], GLRX2 [Liu et al. 2024], PRKCD (protein kinase C delta) [Yang et al. 2019], RAMP1 [Liu et al. 2022], LGALS3 [Sun et al. 2020], SLAMF8 [Zeng et al. 2020], FTH1 [Sun et al. 2023], HMOX1 [Wu et al. 2024], BCL6 [Zhu et al. 2019], MPST (mercaptopyruvate sulfurtransferase) [Zhang et al. 2022], GAREM1 [Wu et al. 2024], RNASE6 [Li et al. 2022], PMVK (phosphomevalonate kinase) [Berner et al. 2023], FIBCD1 [Shahgoli et al. 2023], SLCO4A1 [Koller et al. 2022], SYN2 [Chugh et al. 2015], TRPC4 [Lee et al. 2020], ELN (elastin) [Wu et al. 2020], NALCN (sodium leak channel, non-selective) [Wang et al. 2023], NINJ2 [Zhang et al. 2024], ADAMTS15 [Zuo et al. 2023], LAMB3 [Liu et al. 2024], MMP19 [Bister et al. 2004], SLCO2A1 [Nakanish et al. 2020], STEAP4 [Qiao et al. 2024], GPSM1 [Yan et al. 2022], CYP4B1 [Ashkar et al. 2004], LDHA (lactate dehydrogenase A) [Song et al. 2019], GNMT (glycine N-methyltransferase) [Liu et al. 2020], PTDSS1 [Sekar et al. 2022], B3GNT7 [Wang et al. 2023], RPSA (ribosomal protein SA) [Jiang et al. 2023], BCAT1 [Papathanassiu et al. 2017], RPS5 [Li et al. 2024], SFRP4 [Kida et al. 2023], CD1C [Benlahrech et al. 2015], TRPM8 [Straub, 2014], CST1 [Du et al. 2023], UCMA (upper zone of growth plate and cartilage matrix associated) [Seuffert et al. 2018], RSPO2 [Li et al. 2020], ARMS2 [Zeng et al. 2013], ALOX15 [Caravaca et al. 2022], CTLA4 [Miska et al. 2018], PENK (proenkephalin) [Hu et al. 2016], IL1B [Yi et al. 2018], SCUBE2 [Lin et al. 2024], CTSG (cathepsin G) [Gao et al. 2018], CX3CR1 [Feng et al. 2015], CX3CR1 [Flamant et al. 2021], FRZB (frizzled related protein) [Huang et al. 2021], ZNF365 [Wu et al. 2021], CMA1 [Carl-McGrath et al. 2009], E2F7 [Zhou et al. 2022], RGS4 [Meng et al. 2018], THBS4 [Mäemets-Allas et al. 2023], GDF6 [Cui et al. 2021], ENPP2 [Chattopadhyay et al. 2023], GDF15 [Schwarz et al. 2023], TGFB2 [Huang et al. 2015], C1QTNF3 [Lv et al. 2022], MYOC (myocilin) [Itakura et al. 2015], CXCL10 [Gotsch et al. 2007], MME (membrane metalloendopeptidase) [Wang et al. 2023], BDNF (brain derived neurotrophic factor) [Carniel and da Rocha, 2021], EGR3 [Kwon et al. 2024], PTN (pleiotrophin) [Linnerbauer et al. 2022], OLR1 [Kang et al. 2015], THY1 [Schubert et al. 2011], MDK (midkine) [Sakamoto and Kadomatsu, 2012], ANGPTL7 [Zhao et al. 2019], TNFRSF11B [Zhang et al. 2023], DIO2 [Cheng et al. 2012], TAGLN3 [Arnaud et al. 2022], SMOC2 [Huang et al. 2024], PLAC8 [Zhao et al. 2022], HTR2B [Lu et al. 2021], UCHL1 [Zhu et al. 2024], LUM (lumican) [Li et al. 2021], TP73 [Zhang et al. 2023], TRPV3 [Szöllősi et al. 2018], PRF1 [Sidore et al. 2021], PROS1 [Maimon et al. 2021], CD7 [Liu et al. 2000], ARID5A [Nyati et al. 2020], FREM1 [Kashem et al. 2021], SLC26A4 [Do et al. 2021], ESR1 [Zhang et al. 2023], NOS2 [Béchade et al. 2014], SNCA (synuclein alpha) [Forloni, 2023], EPHA3 [Xu et al. 2023], ACE (angiotensin I converting enzyme) [Gaddam et al. 2017], NT5E [Schädlich et al. 2022], PDE5A [Li et al. 2021], PLCE1 [Li et al. 2019], PAK1 [Chen et al. 2024], SOX8 [Hu et al. 2021], APOE (apolipoprotein E) [Christensen et al. 2011], P2RX1 [Wang et al. 2022], BCL6B [Cai et al. 2019], EGR2 [Bo et al. 2022], GLI1 [Sheng et al. 2021], PRDM1 [Lee et al. 2020], SCT (secretin) [Sato et al. 2017], HLA-DPA1 [Lantermann et al. 2002], FCN1 [Munthe-Fog et al. 2012], EGR1 [Lehman et al. 2022], TFPI2 [Hisaka et al. 2004], DUSP6 [Liu et al. 2022], IL34 [Lin et al. 2019], IER3 [Arlt and Schäfer, 2011], RUNX1 [Lappas, 2018], LTB (lymphotoxin beta) [Giles et al. 2018], SCN8A [Alrashdi et al. 2021], TPM3 [Mansfield et al. 2016], LAMP5 [Gracia-Maldonado et al. 2022], IGFBP2 [Wang et al. 2023], RHCG (Rh family C glycoprotein) [Liu et al. 2024], GZMB (granzyme B) [Turner et al. 2019], CXCL9 [Richmond et al. 2023], MMP25 [Blumenthal et al. 2010], PALM3 [Chen et al. 2017], PRSS8 [Keppner et al. 2016], SELL (selectin L) [Kumari et al. 2021], GZMA (granzyme A) [Garzón-Tituaña et al. 2021], KCNN3 [Dolga et al. 2012], FSCN1 [Huang et al. 2022], HAS3 [Lee et al. 2022], SYTL2 [Xie et al. 2024], SV2A [Metaxas et al. 2019], PTPRH (protein tyrosine phosphatase receptor type H) [Chen et al. 2022], LSP1 [Le et al. 2015], TREH (trehalase) [Panigrahi et al. 2019], CXCL11 [Kochumon et al. 2020], CXCL3 [Bao et al. 2024], CXCL2 [Tsutsui et al. 2018], CXCL14 [Witte et al. 2017], ADAMTS8 [Wågsäter et al. 2008], ANGPTL8 [Abu-Farha et al. 2023] and FMOD (fibromodulin) [Zhao et al. 2023] have been observed to be altered in inflammation. IGFBP2 [Wang et al. 2023] HOPX (HOP homeobox) [Güleç et al. 2014], EREG (epiregulin) [Han et al. 2022], KCNIP2 [Liu et al. 2019], AQP4 [Berland et al. 2018], CCN5 [Yoon et al. 2010], VDR (vitamin D receptor) [Simpson, 2011], KCNK3 [Lambert et al. 2018], RBP4 [Gao et al. 2016], FGFR4 [Liu et al. 2024], TIMP4 [Yarbrough et al. 2014], LYVE1 [Jiang et al. 2020], LGR6 [Zhao et al. 2024], SOX2 [Zhao et al. 2018], NPPC (natriuretic peptide C) [Agrawal et al. 2019], SLC7A11 [Zhang et al. 2022], TFEC (transcription factor EC) [Zhao et al. 2021], AGTR1 [Kelly et al. 2011], ITGA7 [Bugiardini et al. 2022], FHL2 [Friedrich et al. 2014], CD38 [Guan et al. 2017], HCN4 [Wei-qing et al. 2011], POMC (proopiomelanocortin) [Chalmers et al. 2008], APOB (apolipoprotein B) [Ye et al. 2018], BMAL1 [Yu et al. 2022], HEY2 [Yu et al. 2010], RAP1GAP [Gao et al. 2021], IGFBP2 [Sharples et al. 2013], FABP4 [Sun et al. 2021], C1QTNF1 [Wu et al. 2018], LMCD1 [Ferreira et al. 2019], LOX (lysyl oxidase) [Galán et al. 2017], DMPK (DM1 protein kinase) [O’Cochlain et al. 2004], EDN1 [Castro et al. 2007], CTSB (cathepsin B) [Wu et al. 2015], KCNH2 [Bains et al. 2022], HMOX1 [Song et al. 2024], CORIN (corin, serine peptidase) [Wang et al. 2008], CA14 [Vargas and Alvarez, 2012], TRPC4 [Camacho Londoño et al. 2015], ELN (elastin) [Zhang et al. 2022], ADAMTS2 [Wang et al. 2017], LDHA (lactate dehydrogenase A) [Dai et al. 2020], LEF1 [Lai et al. 2019], IRX2 [Wang et al. 2023], CX3CR1 [Weisheit et al. 2021], RGS4 [Wang et al. 2008], THBS4 [Sure and Katakam, 2016], GDF15 [Hanatani et al. 2014], KIT (KIT proto-oncogene, receptor tyrosine kinase) [Sonnenschein et al. 2021], ABCG2 [Higashikuni et al. 2012], MDK (midkine) [Zhang et al. 2020], DIO2 [Ferdous et al. 2020], PCDH17 [Long et al. 2020], UCHL1 [Cao et al. 2023], LUM (lumican) [Rixon et al. 2023], TRPV3 [Qi et al. 2019], SLC26A4 [Tang et al. 2021], ACE (angiotensin I converting enzyme) [Yuan et al. 2017], PDE5A [Liu et al. 2018], PAK1 [Wang et al. 2014], APOE (apolipoprotein E) [Qin et al. 2010], EGR1 [Hsu et al. 2013], RUNX1 [Xu et al. 2023], KIF1A [Akasaka et al. 2020], ATRNL1 [He et al. 2024] and ANGPTL8 [Hu et al. 2022] plays a vital role in the development of HCM. HOPX (HOP homeobox) [Trivedi et al. 2011], FCN3 [Prohászka et al. 2013], SFRP5 [An et al. 2021], CD163 [Ptaszynska-Kopczynska et al. 2016], TFRC (transferrin receptor) [Zhou et al. 2024], AQP4 [Xu et al. 2024], CCN5 [Huang et al. 2020], VDR (vitamin D receptor) [Hao and Chen, 2019], SERPINE1 [Li et al. 2024], AZGP1 [Huscher et al. 2021], VSIG4 [Xie et al. 2022], FGF12 [Liu et al. 2022], RBP4 [An et al. 2021], HMGCS2 [Wang et al. 2024], FGFR4 [Fuchs et al. 2023], TIMP4 [Chaturvedi and Tyagi, 2016], SOX2 [Liang et al. 2021], NPPC (natriuretic peptide C) [Agrawal et al. 2019], PRODH (proline dehydrogenase 1) [Moreira et al. 2019], BMP7 [Aluganti Narasimhulu et al. 2020], SLC7A11 [Wang et al. 2023], AGTR1 [de Denus et al. 2018], PTX3 [Helleberg et al. 2022], GPD1L [Valdivia et al. 2009], ITGA7 [Bugiardini et al. 2022], ATP2A2 [Angrisano et al. 2014], HCN4 [Paterek et al. 2021], POMC (proopiomelanocortin) [Bienertová-Vasků et al. 2009], APOB (apolipoprotein B) [Li et al. 2024], BMAL1 [Ma et al. 2024], HEY2 [Liu et al. 2010], IGFBP2 [Girerd et al. 2020], FABP4 [Harada et al. 2020], LOX (lysyl oxidase) [López et al. 2013], ADAMTS5 [Barallobre-Barreiro et al. 2021], HAS2 [Chowdhury et al. 2017], NQO1 [Blanco et al. 2008], PRDX6 [Xiong et al. 2024], EDN1 [Taylor et al. 2009], HSPB7 [Cappola et al. 2010], CTSB (cathepsin B) [Wu et al. 2015], RAMP1 [Cueille et al. 2002], LGALS3 [Vlachou et al. 2022], KCNH2 [Sugiyama et al. 2011], F8 [Raffield et al. 2020], CD276 [Anzalone et al. 2013], CORIN (corin, serine peptidase) [Dong et al. 2010], ELN (elastin) [Hou et al. 2021], SLC6A6 [Li et al. 2023], PALLD (palladin, cytoskeletal associated protein) [Liu et al. 2021], DHRS7C [Lu et al. 2012], TKT (transketolase) [Wang et al. 2022], SFRP4 [Ren et al. 2024], IRX1 [Zeng et al. 2019], ALOX15 [Sandstedt et al. 2018], CTLA4 [Shang et al. 2024], PENK (proenkephalin) [Siranart et al. 2023], RGS4 [Borges et al. 2023], THBS4 [Meng et al. 2024], CRISPLD1 [Khadjeh et al. 2020], GDF15 [Kosum et al. 2024], CXCL10 [Altara et al. 2016], BDNF (brain derived neurotrophic factor) [Behnoush et al. 2024], THY1 [Li et al. 2020], ABCG2 [Higashikuni et al. 2012], ANGPTL7 [Zhang et al. 2020], SMOC2 [Chen et al. 2024], UCHL1 [Wu et al. 2022], LUM (lumican) [Engebretsen et al. 2013], NOS2 [Nishijima et al. 2011], ACE (angiotensin I converting enzyme) [McNamara et al. 2004], PDE5A [Westermann et al. 2012], PAK1 [Ai et al. 2011], SSC5D [Ge et al. 2023], IL34 [Tao et al. 2017], BOC (BOC cell adhesion associated, oncogene regulated) [Ahmed et al. 2020], RUNX1 [Qi et al. 2023], GZMB (granzyme B) [Zeglinski and Granville, 2020], KCNN3 [Rahm et al. 2021], CXCL14 [Zeng et al. 2019], ADAMTS8 [Li et al. 2023], FMOD (fibromodulin) [Andenæs et al. 2018] and CX3CR1 [Flamant et al. 2021] expression were significantly higher in patients with heart failure. The expression levels of ALOX15B [Wuest et al. 2014], SFRP5 [Jia et al. 2023], F5 [Redondo et al. 1999], CD163 [Aristoteli et al. 2006], PLA2G2A [Shuvalova et al. 2015], VDR (vitamin D receptor) [Raljević et al. 2021], SERPINE1 [Koch et al. 2010], AZGP1 [Huscher et al. 2021], FKBP5 [Wang et al. 2020], RBP4 [Ji et al. 2023], C3 [Cai et al. 2019], ITLN1 [Güçlü-Geyik et al. 2022], TIMP4 [Kuliczkowski et al. 2017], LYVE1 [Cho et al. 2019], NPPC (natriuretic peptide C) [Prickett et al. 2020], MLXIPL (MLX interacting protein like) [Loika et al. 2023], AGTR1 [Tran et al. 2024], PTX3 [Knapp et al. 2024], GPD1L [Westaway et al. 2011], FN1 [Page et al. 2022], CMTM5 [Zhang et al. 2017], APOB (apolipoprotein B) [Pencina et al. 2015], IGFBP2 [Wang et al. 2023], FABP4 [Zhao et al. 2024], C1QTNF1 [Zhao et al. 2020], LOX (lysyl oxidase) [Ma et al. 2011], ADAMTS5 [Wang et al. 2019], CD63 [Murakami et al. 1996], NQO1 [Zhou et al. 2008], PROCR (protein C receptor) [Kallel et al. 2012], SIGLEC1 [Xiong et al. 2009], EDN1 [Gupta, 2022], LGALS3 [Li et al. 2022], F8 [Raffield et al. 2020], HMOX1 [Sarutipaiboon et al. 2020], CORIN (corin, serine peptidase) [Wang et al. 2021], NINJ2 [Yan et al. 2024], SFRP4 [Ji et al. 2017], ALOX15 [Kaur et al. 2018], IL1B [Dorajoo et al. 2023], CX3CR1 [Loh et al. 2023], FRZB (frizzled related protein) [Guan et al. 2021], CRISPLD1 [Wang et al. 2018], GDF15 [Li et al. 2020], APLNR (apelin receptor) [Wang et al. 2020], CXCL10 [Karimabad et al. 2021], BDNF (brain derived neurotrophic factor) [Shobeiri et al. 2023], EGR3 [Li et al. 2014], OLR1 [Mohammed et al. 2022], ABCG2 [Han et al. 2015], S1PR5 [Xiong et al. 2022], ESR1 [Zhang et al. 2023], CUBN (cubilin) [Park et al. 2020], ACE (angiotensin I converting enzyme) [Danchin et al. 2006], PDE5A [Dang et al. 2021], GNB3 [von Beckerath et al. 2003], APOE (apolipoprotein E) [Anoop et al. 2010], F2RL3 [Zhao et al. 2022], F2R [Gigante et al. 2007], TNNT1 [Guay et al. 2016], FCN1 [Mo et al. 2019], EGR1 [Peng and Xiang, 2019], TFPI2 [Zhou et al. 2019], IL34 [Fan et al. 2016], RUNX1 [Zhong et al. 2022], GZMB (granzyme B) [Zeglinski and Granville, 2020], TREH (trehalase) [Jamialahmadi et al. 2022], CXCL2 [Yang et al. 2020], CXCL14 [Schories et al. 2023] and ANGPTL8 [Morinaga et al. 2023] have been proved to be altered in coronary artery disease. The study showed that ALOX15B [Magnusson et al. 2012], EREG (epiregulin) [Wang et al. 2023], SFRP5 [Tong et al. 2019], CD163 [Zhang et al. 2024], PLA2G2A [Monroy-Muñoz et al. 2017], VDR (vitamin D receptor) [Schnatz et al. 2012], SERPINE1 [Koch et al. 2010], AZGP1 [Huscher et al. 2021], ELANE (elastase, neutrophil expressed) [Wen et al. 2018], RBP4 [Zhou et al. 2018], C3 [Garcia-Arguinzonis et al. 2021], FGFR4 [Sellier et al. 2023], TIMP4 [Hu et al. 2021], NPPC (natriuretic peptide C) [Lumsden et al. 2010], SPP1 [Wu et al. 2024], BMP7 [Davies et al. 2003], SLC7A11 [Wang et al. 2024], CFD (complement factor D) [Murakami et al. 2023], AGTR1 [Campbell et al. 2010], PTX3 [Otani et al. 2024], EPHB2 [Braun et al. 2011], SLC2A1 [Zhou et al. 2024], FHL2 [Chu et al. 2010], CD38 [Kong et al. 2024], FGF7 [Wu et al. 2023], C1QA [Dong et al. 2024], CRLF1 [Lan et al. 2024], POMC (proopiomelanocortin) [Rinne et al. 2018], APOB (apolipoprotein B) [Singh and Prabhakaran, 2024], BMAL1 [Xie et al. 2020], IGFBP2 [Wang et al. 2024], ARG2 [Rafnsson et al. 2020], FABP4 [Gormez et al. 2020], C1QTNF1 [Zhang et al. 2022], LOX (lysyl oxidase) [Ovchinnikova et al. 2014], ADAMTS5 [Didangelos et al. 2012], CD63 [Cha et al. 2004], HAS2 [Zhu et al. 2020], NQO1 [Hur et al. 2010], SIGLEC1 [Nasser et al. 2024], PRDX6 [Phelan et al. 2003], EDN1 [Gupta et al. 2020], CTSB (cathepsin B) [Li et al. 2024], LGALS3 [Djordjevic et al. 2022], HMOX1 [Wu et al. 2022], BCL6 [Wei et al. 2015], RNASE6 [Fang et al. 2021], EPN1 [Brophy et al. 2019], CORIN (corin, serine peptidase) [Jiang et al. 2018], ELN (elastin) [Maedeker et al. 2021], PRIMA1 [Wang et al. 2019], NINJ2 [Zhang et al. 2016], PALLD (palladin, cytoskeletal associated protein) [Hoke et al. 2011], GNMT (glycine N-methyltransferase) [Chen et al. 2012], BCAT1 [Tan et al. 2022], SFRP4 [Zhang et al. 2022], RSPO2 [Singla et al. 2021], CTLA4 [Poels et al. 2020], CTSG (cathepsin G) [Wang et al. 2014], CX3CR1 [Rowinska et al. 2017], ENPP2 [Chattopadhyay et al. 2023], GDF15 [Huang et al. 2021], CXCL10 [Prapiadou et al. 2024], BDNF (brain derived neurotrophic factor) [Bi et al. 2020], PTN (pleiotrophin) [Li et al. 2010], OLR1 [Yang et al. 2023], MDK (midkine) [Zhang et al. 2021], TNFRSF11B [Wang et al. 2020], LUM (lumican) [Onda et al. 2002], TP73 [Ni et al. 2021], ACE (angiotensin I converting enzyme) [Okwan-Duodu et al. 2019], PAK1 [Singh et al. 2015], GNB3 [Grove et al. 2007], APOE (apolipoprotein E) [Marais, 2021], EGR1 [Blaschke et al. 2004], TFPI2 [Crawley et al. 2002], IL34 [Law et al. 2021], TNFSF13B [Fajar et al. 2024], LTB (lymphotoxin beta) [Owens et al. 2010], RSAD2 [Hayderi et al. 2024], ITGAX (integrin subunit alpha X) [Wang et al. 2023], CXCL9 [Wu et al. 2024], GZMB (granzyme B) [Hiebert et al. 2013], TREH (trehalase) [Zhong et al. 2023], CXCL2 [Chang et al. 2024], CXCL14 [He et al. 2023], ADAMTS8 [Wågsäter et al. 2008] and ANGPTL8 [Zheng et al. 2018] were related to atherosclerosis. Altered expression of EREG (epiregulin) [Han et al. 2022], FCN3 [Lidani et al. 2021], CD163 [Walker et al. 2014], CCN5 [Yoon et al. 2010], VDR (vitamin D receptor) [Panizo et al. 2017], SERPINE1 [Khan et al. 2024], VSIG4 [Wang et al. 2023], TIMP4 [Ye et al. 2021], LYVE1 [Jiang et al. 2020], BMP7 [Ouyang et al. 2022], SLC7A11 [Wang et al. 2023], PTX3 [Xu et al. 2022], EPHB2 [Su et al. 2017], TGM2 [Griffin et al. 2020], FHL2 [Anwaier et al. 2022], CRLF1 [Luo et al. 2023], BMAL1 [Hang et al. 2023], GPD1 [Wang et al. 2022], FABP4 [Du et al. 2024], LOX (lysyl oxidase) [López et al. 2013], CTSB (cathepsin B) [Wu et al. 2015], PRKCD (protein kinase C delta) [Chintalgattu and Katwa, 2009], LGALS3 [Fu et al. 2020], CORIN (corin, serine peptidase) [Gladysheva et al. 2023], SMOC1 [Wang and Wu, 2018], ELN (elastin) [Wang et al. 2021], LDHA (lactate dehydrogenase A) [Wu et al. 2024], RPS5 [Zhang et al. 2021], SFRP4 [Matsushima et al. 2010], ALOX15 [Sandstedt et al. 2018], IRX2 [Ma et al. 2023], CYTL1 [Kim et al. 2016], RGS4 [Guo et al. 2021], GDF15 [Guo et al. 2021], CXCL10 [Aziz et al. 2023], BDNF (brain derived neurotrophic factor) [Zou and Hao, 2024], SULF1 [Ji et al. 2024], SMOC2 [Rui et al. 2023], HTR2B [Castillero et al. 2024], UCHL1 [Gong et al. 2021], LUM (lumican) [Mohammadzadeh et al. 2020], TRPV3 [Liu et al. 2018], ACE (angiotensin I converting enzyme) [Liu et al. 2016], APOE (apolipoprotein E) [Hada et al. 2022], GLI1 [Wang et al. 2018], EGR1 [Li et al. 2021], DKK2 [Li et al. 2022], RUNX1 [Ni et al. 2021], GZMB (granzyme B) [Shen et al. 2016], CXCL9 [Lin et al. 2019] and ADAMTS8 [Zha et al. 2022] have been observed to be associated with cardiac fibrosis. EREG (epiregulin) [Song et al. 2022], SFRP5 [Koutaki et al. 2021], SPINK1 [Abass et al. 2022], CD163 [Cinkajzlová et al. 2017], PLA2G2A [Varastehpour et al. 2006], TFRC (transferrin receptor) [Qiu et al. 2024], CCN5 [Kim et al. 2018], PCK1 [Vimaleswaran et al. 2010], VDR (vitamin D receptor) [Akter et al. 2022], SERPINE1 [Su et al. 2023], KCNK3 [Chen et al. 2017], ELANE (elastase, neutrophil expressed) [Mansuy-Aubert et al. 2013], FKBP5 [Willmer et al. 2022], RBP4 [Kilicarslan et al. 2020], CPNE5 [Wang et al. 2015], MOG (myelin oligodendrocyte glycoprotein) [Stiebel-Kalish et al. 2022], C3 [Castellano-Castillo et al. 2018], FGFR4 [Ge et al. 2014], ITLN1 [Güçlü-Geyik et al. 2023], TIMP4 [Sakamuri et al. 2017], LYVE1 [Michurina et al. 2016], NPPC (natriuretic peptide C) [Agrawal et al. 2019], SPP1 [Yang et al. 2019], BMP7 [Casana et al. 2022], SLC7A11 [Li et al. 2023], CFD (complement factor D) [Mathews et al. 2014], AGTR1 [Adiyeva et al. 2023], PTX3 [Bonacina et al. 2019], EPHB2 [Suzuki et al. 2022], GPD1L [He et al. 2017], FHL2 [Sommer et al. 2023], CD38 [Barbosa et al. 2007], OSR1 [Davies et al. 2014], POMC (proopiomelanocortin) [Ito et al. 2013], APOB (apolipoprotein B) [Ying et al. 2012], BMAL1 [Zhan et al. 2024], IGFBP2 [Boughanem et al. 2021], ARG2 [Zaytouni et al. 2017], IL15RA [Loro et al. 2015], GPD1 [Li et al. 2017], FABP4 [Osorio-Conles et al. 2023], PTGS1 [Plaza-Díaz et al. 2017], LOX (lysyl oxidase) [Huang et al. 2021], NQO1 [Di Francesco et al. 2020], EDN1 [Li et al. 2015], CTSB (cathepsin B) [Araujo et al. 2018], PRKCD (protein kinase C delta) [Patel et al. 2016], RAMP1 [Hosono et al. 2024], TSKU (tsukushi, small leucine rich proteoglycan) [Li et al. 2021], LGALS3 [Baek et al. 2015], NID1 [Pérez-Díaz et al. 2023], ACACB (acetyl-CoA carboxylase beta) [Ma et al. 2013], BCL6 [Senagolage et al. 2018], MPST (mercaptopyruvate sulfurtransferase) [Katsouda et al. 2022], CD276 [Picarda et al. 2022], HIF3A [Shen et al. 2020], ADH1B [Morales et al. 2021], ELN (elastin) [Martinez-Santibanez et al. 2015], AIF1L [Parikh et al. 2022], LAMB3 [Jiao et al. 2016], MMP19 [Pendás et al. 2004], SLC6A6 [Bhutia et al. 2022], STEAP4 [Ozmen et al. 2016], GPSM1 [Tang et al. 2024], CHDH (choline dehydrogenase) [Chirita-Emandi et al. 2020], RETSAT (retinol saturase) [Schupp et al. 2009], PNMT (phenylethanolamine N-methyltransferase) [Peters et al. 2003], RPLP0 [Pérez-Gómez et al. 2023], TKT (transketolase) [Tian et al. 2020], TNMD (tenomodulin) [Tolppanen et al. 2010], SFRP4 [Bukhari et al. 2019], TRPM8 [Sanders et al. 2021], ALOX15 [Ke et al. 2012], CTLA4 [Santos et al. 2018], IL1B [Maculewicz et al. 2022], CX3CR1 [Sun et al. 2022], RGS4 [Michaelides et al. 2020], ENPP2 [Nishimura et al. 2014], GDF15 [Siddiqui et al. 2022], C1QTNF3 [Maeda and Wakisaka, 2020], APLNR (apelin receptor) [De Los Santos et al. 2023], CXCL10 [Meyhöfer et al. 2024], BDNF (brain derived neurotrophic factor) [Sandrini et al. 2018], OLR1 [Khaidakov et al. 2011], THY1 [Picke et al. 2018], CHRNA3 [Varga et al. 2018], ABCG2 [Cheng et al. 2017], ANGPTL7 [Abu-Farha et al. 2017], DIO2 [Bradley et al. 2018], PLAC8 [Barragán-Zúñiga et al. 2022], PHOSPHO1 [Suchacki et al. 2020], HTR2B [Choi et al. 2021], LUM (lumican) [Strieder-Barboza et al. 2022], TRPV3 [Hu et al. 2024], ESR1 [Guclu-Geyik et al. 2020], NOS2 [Vilela et al. 2022], EPHA3 [Zhang et al. 2022], ACE (angiotensin I converting enzyme) [Sawada et al. 2015], GNB3 [Ozdemir et al. 2017], APOE (apolipoprotein E) [Farup et al. 2020], GDF9 [Arıkan et al. 2023], SCT (secretin) [Xue et al. 2022], EGR1 [Zhang et al. 2013], IL34 [Chang et al. 2014], RUNX1 [Zhang et al. 2023], GZMB (granzyme B) [El Mesallamy et al. 2014], CXCL9 [Duarte et al. 2015], TREH (trehalase) [Arai et al. 2013], CXCL11 [Kochumon et al. 2020], IGSF1 [Ghanny et al. 2021], CXCL2 [Pan et al. 2024], CXCL14 [Cereijo et al. 2021], ANGPTL8 [Perdomo et al. 2021], C1QTNF12 [Wei et al. 2012] and INHBE (inhibin subunit beta E) [Deaton et al. 2022] might serve as therapeutic targets for obesity. The altered expression of FCN3 [Lu et al. 2012], SFRP5 [Li et al. 2019], CD163 [Jasiewicz et al. 2014], TFRC (transferrin receptor) [Yang et al. 2024], AQP4 [González-Marrero et al. 2022], VDR (vitamin D receptor) [Nabil et al. 2024], KCNK3 [Lambert et al. 2018], FGF12 [Yeo et al. 2020], RBP4 [Li et al. 2019], HMGCS2 [Singh et al. 2018], MOG (myelin oligodendrocyte glycoprotein) [Chaudhuri et al. 2022], C3 [Chen et al. 2020], FGFR4 [Al-Daghri et al. 2017], TIMP4 [Wetzl et al. 2017], GRIP1 [Wang et al. 2016], SOX2 [Jiang et al. 2022], NPPC (natriuretic peptide C) [Agrawal et al. 2019], EIF4EBP1 [Li et al. 2022], SPP1 [Chen et al. 2023], BMP7 [Liu et al. 2016], KCNK2 [Shima et al. 2024], PPP1R1A [Grimm et al. 2023], CFD (complement factor D) [Cheng et al. 2024], AGTR1 [Zeng et al. 2023], PTX3 [Helleberg et al. 2022], SEPTIN4 [Zhang et al. 2023], EPHB2 [Xing et al. 2019], ATP2A2 [Wang et al. 2024], FHL2 [Li et al. 2019], CD38 [Qiu et al. 2023], FGF7 [Zhou et al. 2019], OSR1 [Brown et al. 2021], POMC (proopiomelanocortin) [do Carmo et al. 2011], APOB (apolipoprotein B) [Dong et al. 2022], BMAL1 [Kurbatova et al. 2014], IGFBP2 [Yang et al. 2020], CHGB (chromogranin B) [Zhang et al. 2009], FABP4 [Furuhashi et al. 2015], SDK1 [Oguri et al. 2010], LOX (lysyl oxidase) [Vadasz et al. 2019], HAS2 [Tseng et al. 2022], NQO1 [Kim et al. 2013], CAMK1D [Du et al. 2021], NGFR (nerve growth factor receptor) [Goten et al. 2021], PRDX6 [Liao et al. 2023], EDN1 [Schiffrin, 2018], PRKCD (protein kinase C delta) [Novokhatska et al. 2013], RAMP1 [Sabharwal et al. 2019], LGALS3 [Barman et al. 2021], KCNK1 [Shima et al. 2024], HMOX1 [Song et al. 2024], BCL6 [Chen et al. 2019], HIF3A [Xu et al. 2023], CORIN (corin, serine peptidase) [Wang et al. 2008], ADH1B [Yokoyama et al. 2013], TRPC4 [Alzoubi et al. 2013], ELN (elastin) [Maedeker et al. 2021], LDHA (lactate dehydrogenase A) [Wu et al. 2024] PNMT (phenylethanolamine N-methyltransferase) [Huang et al. 2011], TRPM8 [Huang et al. 2017], CTLA4 [Tomaszewski et al. 2022], IL1B [Yanagisawa et al. 2009], RXFP2 [Zhang et al. 2024], CX3CR1 [Ahadzadeh et al. 2018], THBS4 [Palao et al. 2018], GDF15 [Sökmen et al. 2019], APLNR (apelin receptor) [Stanchev et al. 2024], CXCL10 [Hong et al. 2022], BDNF (brain derived neurotrophic factor) [Nemcsik et al. 2016], LEFTY2 [Ashiq et al. 2023], OLR1 [Sciacqua et al. 2014], CHRNA3 [Wu et al. 2020], MDK (midkine) [Guzel et al. 2018], DIO2 [Kang et al. 2023], CHRM3 [Cowley et al. 2018], HTR2B [Fang et al. 2024], UCHL1 [Tang et al. 2024], SLC26A4 [Kim et al. 2017], ESR1 [Zhou et al. 2022], NOS2 [Shnayder et al. 2021], ACE (angiotensin I converting enzyme) [Ishigami et al. 1995], PDE5A [Li et al. 2021], PLCE1 [Atchison et al. 2020], SCG2 [Wen et al. 2007], KCNA5 [Vera-Zambrano et al. 2023], GNB3 [Sydorchuk et al. 2023], APOE (apolipoprotein E) [de Leeuw et al. 2004], F2RL3 [Deng et al. 2024], F2R [Gigante et al. 2007], GLI1 [Chu et al. 2018], PRDM1 [Wu et al. 2023], SCT (secretin) [Zaw et al. 2019], EGR1 [Suginobe et al. 2023], RUNX1 [Ma et al. 2024], TNFRSF4 [Mashimo et al. 2008], TPM3 [Xu et al. 2021], DLGAP1 [Takahashi et al. 2021], GZMB (granzyme B) [Mao et al. 2018], CXCL9 [Jandl et al. 2022], SLC16A9 [Simino et al. 2013], KCNN3 [Koot et al. 2016], TREH (trehalase) [Forte et al. 2021], ADAMTS8 [Omura et al. 2019], ANGPTL8 [Jiao et al. 2023] and FMOD (fibromodulin) [Liang et al. 2023] were observed to be associated with the progression of hypertension. FCN3 [Barkai et al. 2019], SSTR5 [Sawicki, 2006], SFRP5 [Li et al. 2019], SLAMF1 [Tabassum et al. 2012], SPINK1 [Schneider et al. 2002], CD163 [Cao et al. 2024], PLA2G2A [Monroy-Muñoz et al. 2017], AQP4 [Li et al. 2022], CCN5 [Kim et al. 2018], PCK1 [Hu et al. 2019], VDR (vitamin D receptor) [Alfaqih et al. 2022], SERPINE1 [Fan et al. 2018], ACVR1C [Emdin et al. 2019], GRB14 [Popineau et al. 2016], FKBP5 [Sidibeh et al. 2018], RBP4 [Ji et al. 2023], HMGCS2 [Shukla et al. 2017], C3 [Castellano-Castillo et al. 2018], FGFR4 [Ge et al. 2014], ITLN1 [Güçlü-Geyik et al. 2023], TIMP4 [Kuliczkowski et al. 2017], LYVE1 [Michurina et al. 2016], SOX2 [Gu et al. 2011], NPPC (natriuretic peptide C) [Prickett et al. 2019], SPP1 [Xiao et al. 2024], BMP7 [Casana et al. 2022], SLC7A11 [Xia et al. 2023], MLXIPL (MLX interacting protein like) [Mtiraoui et al. 2012], AGTR1 [Liu et al. 2024], PTX3 [Mutlu et al. 2017], EPHB2 [Broquères-You et al. 2012], SLC2A1 [Petakh et al. 2024], TGM2 [Kolokotronis et al. 2024], FHL2 [Habibe et al. 2022], CD38 [Antonelli and Ferrannini, 2004], FGF7 [Alwahsh et al. 2021], OSR1 [Guo et al. 2021], HCN4 [Parveen et al. 2023], POMC (proopiomelanocortin) [Ito et al. 2013], APOB (apolipoprotein B) [Hwang et al. 2012], BMAL1 [Gao et al. 2023], IGFBP2 [Boughanem et al. 2021], CHGB (chromogranin B) [Herold et al. 2020], FCGR3A [Alizadeh et al. 2007], IL15RA [Bobbala et al. 2017], GPD1 [Li et al. 2017], FABP4 [Wang et al. 2021], TPO (thyroid peroxidase) [Chang et al. 1998], NQO1 [Zhang et al. 2022], SIGLEC1 [Guo et al. 2024], CAMK1D [Du et al. 2021], PRDX6 [Pacifici et al. 2014], EDN1 [Maslat et al. 2023], CTSB (cathepsin B) [Li et al. 2023], TSKU (tsukushi, small leucine rich proteoglycan) [Li et al. 2024], LGALS3 [Zhou et al. 2022], NID1 [Khattab and Torkamani, 2022], HMOX1 [Jirásková et al. 2023], AKR1B1 [Dieter et al. 2022], ACACB (acetyl-CoA carboxylase beta) [Chan et al. 2022], HIF3A [Main et al. 2016], CORIN (corin, serine peptidase) [Fathy et al. 2015], ADH1B [Yokoyama et al. 2013], TRPC4 [Sung et al. 2014], NINJ2 [Peng et al. 2023], STEAP4 [Zhong et al. 2024], GPSM1 [Ding et al. 2020], LDHA (lactate dehydrogenase A) [Sanchez et al. 2021], ETNPPL (ethanolamine-phosphate phospho-lyase) [Wang et al. 2023], HK3 [Islam et al. 2024], BCAT1 [Alfaqih et al. 2021], NMRAL1 [Gao et al. 2022], SFRP4 [Bukhari et al. 2019], FOXA1 [Fogarty et al. 2014], ALOX15 [He et al. 2023], CTLA4 [Ahmadi et al. 2013], PENK (proenkephalin) [van Hateren et al. 2015], IL1B [Jiao et al. 2021], HBB (hemoglobin subunit beta) [Liu et al. 2022], SCUBE2 [Ali et al. 2019], CX3CR1 [Njerve et al. 2012], RGS4 [Zhang et al. 2022], ENPP2 [Nishimura et al. 2014], GDF15 [Al-Kuraishy et al. 2022], TGFB2 [Gao et al. 2022], CXCL10 [Antonelli et al. 2014], BDNF (brain derived neurotrophic factor) [Bi et al. 2020], LRRN1 [Antikainen et al. 2024], LEFTY2 [Ashiq et al. 2023], ABCG2 [Szabó et al. 2021], ANGPTL7 [Xu et al. 2020], TNFRSF11B [Biscetti et al. 2014], DIO2 [Nair et al. 2012], CHRM3 [Guo et al. 2006], PLAC8 [Sasaki et al. 2015], PHOSPHO1 [Suchacki et al. 2020], HTR2B [Choi et al. 2021], UCHL1 [Costes et al. 2011], LUM (lumican) [Strieder-Barboza et al. 2022], ESR1 [Dahlman et al. 2008], NOS2 [Gusti et al. 2021], CUBN (cubilin) [Tsekmekidou et al. 2020], ACE (angiotensin I converting enzyme) [Pedersen-Bjergaard et al. 2001], PAK1 [Veluthakal et al. 2018], GNB3 [Rizvi et al. 2017], APOE (apolipoprotein E) [El-Lebedy et al. 2018], EGR2 [Lu et al. 2018], OAS3 [Park et al. 2018], SCT (secretin) [Gilliam-Vigh et al. 2023], HLA-DPA1 [Varney et al. 2010], FCN1 [Anjosa et al. 2016], EGR1 [Ke et al. 2023], TFPI2 [Guo et al. 2023], IL34 [Chang et al. 2014], RUNX1 [Zhong et al. 2022], KIF1A [Charles Bronson et al. 2021], KCNJ15 [Fukuda et al. 2013], GZMB (granzyme B) [El Mesallamy et al. 2014], GRIA3 [Li et al. 2019], COLEC12 [Peng et al. 2015], GPR27 [Chopra et al. 2020], TREH (trehalase) [Arai et al. 2013], KCNIP1 [Xu et al. 2018], CXCL11 [Kochumon et al. 2020], CXCL14 [Cereijo et al. 2021], ANGPTL8 [Zheng et al. 2018], C1QTNF12 [Wei et al. 2012], INHBE (inhibin subunit beta E) [Sugiyama et al. 2018] and FMOD (fibromodulin) [Lee et al. 2021] are the main regulators of diabetes mellitus. SFRP5 [Ding et al. 2022], F5 [Ridker et al. 1995], CD163 [Shao et al. 2020], AQP4 [Warth et al. 2007], CCN5 [Zolfaghari et al. 2022], VDR (vitamin D receptor) [Raljević et al. 2021], SERPINE1 [Morange et al. 2007], AZGP1 [Huscher et al. 2021], VSIG4 [Wang et al. 2023], RBP4 [Zhang et al. 2021], C3 [Fang et al. 2023], TIMP4 [Weir et al. 2011], SPP1 [Shen et al. 2024], BMP7 [Jin et al. 2018], SLC7A11 [Li et al. 2023], TFEC (transcription factor EC) [Wang et al. 2022], CFD (complement factor D) [Hao et al. 2023], AGTR1 [Tran et al. 2024], PTX3 [Xu et al. 2022], EPHB2 [Jiang et al. 2024], FGF7 [Mei et al. 2022], FN1 [Yao et al. 2022], HCN4 [Song et al. 2011], STXBP2 [Yamada et al. 2017], POMC (proopiomelanocortin) [Gumede et al. 2024], APOB (apolipoprotein B) [Marston et al. 2022], BMAL1 [Ruan et al. 2024], RAP1GAP [Shan et al. 2024], FABP4 [Aleksandrova et al. 2019], LOX (lysyl oxidase) [González-Santamaría et al. 2016], ADAMTS5 [Lee et al. 2011], CD63 [van der Zee et al. 2006], EDN1 [Palacín et al. 2009], DHCR24 [Han et al. 2019, PDPN (podoplanin) [Cimini et al. 2019], CTSB (cathepsin B) [Wu et al. 2015], FCER1G [Xiao et al. 2021], LGALS3 [Chalise et al. 2022], KCNH2 [Wang et al. 2020], HMOX1 [Yang et al. 2024], CORIN (corin, serine peptidase) [Feistritzer and Metzler, 2016], TRPC4 [Jung et al. 2011], ELN (elastin) [Skjøt-Arkil et al. 2013], ADAMTS2 [Lee et al. 2012], QSOX1 [Vanhaverbeke et al. 2019], BCAT1 [Lai et al. 2021], TRPM8 [Alves et al. 2020], LEF1 [Cho et al. 2020], CTLA4 [Yip et al. 2007], PENK (proenkephalin) [Ng et al. 2014], IL1B [Pan et al. 2022], CX3CR1 [Pucci et al. 2013], F2RL2 [Wu et al. 2023], GDF15 [Dogon et al. 2024], BDNF (brain derived neurotrophic factor) [Okada et al. 2012], OLR1 [Trabetti et al. 2006], ABCG2 [Higashikuni et al. 2010], MDK (midkine) [Takenaka et al. 2009], HTR2B [Snider et al. 2021], UCHL1 [Wu et al. 2022], ESR1 [Puzianowska-Kuźnicka, 2012], NOS2 [Zheng et al. 2004], CUBN (cubilin) [McMahon et al. 2014], ACE (angiotensin I converting enzyme) [Liu et al. 2016], PDE5A [Li et al. 2021], PAK1 [Liu et al. 2020], GNB3 [Chang et al. 2012], APOE (apolipoprotein E) [Shao et al. 2022], F2RL3 [Corbin et al. 2022], F2R [Gigante et al. 2010], EGR2 [Bo et al. 2022], GLI1 [Wang et al. 2018], EGR1 [Li et al. 2021], TFPI2 [Guo et al. 2023], DUSP6 [Wu and Zhang, 2024], IL34 [Kashiwagi et al. 2022], GZMB (granzyme B) [Xu et al. 2024], CXCL9 [Lin et al. 2019], RUNX1 [Ni et al. 2021], TREH (trehalase) [Farag et al. 2023], CXCL2 [Guo et al. 2020] and ADAMTS8 [Zhao et al. 2011] are associated to the risk of myocardial infarction. SFRP5 [Zeng et al. 2023], CD163 [Chen et al. 2020], AQP4 [Jiang et al. 2021], TIMP4 [Takawale et al. 2014], SOX2 [Li et al. 2023], NPPC (natriuretic peptide C) [Tarazón et al. 2019], SPP1 [Yuan and Fu, 2024], SLC7A11 [Pang et al. 2024], TGFA (transforming growth factor alpha) [Herrmann et al. 2011], PTX3 [Shimizu et al. 2015], EPHB2 [Yang et al. 2016], GPD1L [Valdivia et al. 2009], FHL2 [Goltz et al. 2016], CD38 [Zhang et al. 2020], FN1 [Zhang et al. 2022], HCN4 [Yu et al. 2011], APOB (apolipoprotein B) [Benn et al. 2007], BMAL1 [Yu et al. 2022], LOX (lysyl oxidase) [Valls-Lacalle et al. 2021], PRDX6 [Mu et al. 2022], CTSB (cathepsin B) [Wu et al. 2022], PRKCD (protein kinase C delta) [Sivaraman et al. 2009], LGALS3 [Rong et al. 2020], HMOX1 [Tan et al. 2024], CORIN (corin, serine peptidase) [Wang et al. 2018], ELN (elastin) [Wang et al. 2021], CACNA2D3 [Liu et al. 2024], STEAP4 [Luo et al. 2019], BCAT1 [Fu et al. 2022], SFRP4 [Zeng et al. 2019], TRPM8 [Cheng et al. 2019], ALOX15 [Cai et al. 2023], IL1B [Yu et al. 2010], GDF15 [Dogon et al. 2024], APLNR (apelin receptor) [Zheng et al. 2023], CXCL10 [Safa et al. 2016], BDNF (brain derived neurotrophic factor) [Fioranelli et al. 2023], PTN (pleiotrophin) [Christman et al. 2005], MDK (midkine) [Horiba et al. 2006], UCHL1 [Geng et al. 2022], ESR1 [Kabir et al. 2015], ACE (angiotensin I converting enzyme) [Pepine, 1998], PLCE1 [Li et al. 2019], PAK1 [Liu et al. 2020], APOE (apolipoprotein E) [Shao et al. 2022], EGR1 [Li et al. 2024], TFPI2 [Luo et al. 2023], DUSP6 [Wu et al. 2019], IL34 [Zhuang et al. 2023], RUNX1 [Zhang et al. 2022], GZMB (granzyme B) [Santos-Zas et al. 2021], EDA2R [Guan et al. 2023], FSCN1 [Huang et al. 2023], HAS3 [Piroth et al. 2022], TREH (trehalase) [Wang et al. 2024] and CXCL14 [Liu et al. 2021] are found to be associated with myocardial ischemia. Studies show that CD163 [Urbonaviciene et al. 2011], PLA2G2A [Akinkuolie et al. 2019], GPD1L [Huang et al. 2018], GSTM3 [Juang et al. 2020], APOB (apolipoprotein B) [Nakano et al. 2008], HEY2 [Bezzina et al. 2013], FABP4 [Saito et al. 2021], CTSB (cathepsin B) [Dai et al. 2021], LGALS3 [Maiolino et al. 2015], GDF15 [Andersson et al. 2020], CUBN (cubilin) [McMahon et al. 2014], ACE (angiotensin I converting enzyme) [Domanski et al. 1999], GNB3 [Birkner et al. 2023], EGR1 [Xie et al. 2023], KCNN3 [Ivanova et al. 2017] and TREH (trehalase) [Iwai et al. 2012] are involved in the cardiac death. CD163 [Sbrana et al. 2020], PLA2G2A [Lemaitre et al. 2016], AQP4 [Nakayama et al. 2016], FGF12 [Velíšková et al. 2021], IL18R1 [Zhang et al. 2022], PTX3 [Ristagno et al. 2015], GPD1L [Valdivia et al. 2009], KCNH2 [Bileišienė et al. 2021], ELN (elastin) [Markush et al. 2023], GDF15 [Herrmann et al. 2022], BDNF (brain derived neurotrophic factor) [D’Cruz et al. 2002, PTN (pleiotrophin) [Liu et al. 2021], UCHL1 [Ebner et al. 2020], KCNA5 [Nielsen et al. 2007] and APOE (apolipoprotein E) [Kida et al. 1995] plays an essential role in facilitating cardiac arrest. Altered expression of TFRC (transferrin receptor) [Xu et al. 2015], CCN5 [Kim et al. 2018], FKBP5 [Wada et al. 2021], RBP4 [Bobbert et al. 2009], HMGCS2 [Wang et al. 2024], XRCC4 [Fredette et al. 2019], NPPC (natriuretic peptide C) [Tarazón et al. 2019], BMP7 [Ouyang et al. 2022], SLC7A11 [Lin et al. 2024], HMOX2 [Cetin-Atalay et al. 2023], GPD1L [Huang et al. 2018], DIRAS3 [Zhuo et al. 2017], ITGA7 [Esposito et al. 2013], FHL2 [Arimura et al. 2007], CD38 [Wang et al. 2023], HCN4 [Yildirim et al. 2022], APOB (apolipoprotein B) [Yokoyama et al. 2004], BMAL1 [Li et al. 2020], HEY2 [Sakata et al. 2002], IGFBP2 [Yu et al. 2023], LPCAT3 [Iqbal et al. 2023], LOX (lysyl oxidase) [Sivakumar et al. 2008], DMPK (DM1 protein kinase) [Damanafshan et al. 2018], NQO1 [Wu et al. 2022], EDN1 [Matsa et al. 2014], DHCR24 [Dong et al. 2018], HSPB7 [Stark et al. 2010], CTSB (cathepsin B) [Mehra et al. 2017], LGALS3 [Nguyen et al. 2019], KCNH2 [Maddali et al. 2022, HMOX1 [Zhen et al. 2024], HIF3A [Guo et al. 2021], CORIN (corin, serine peptidase) [Tripathi et al. 2019], ADAMTS2 [Rau et al. 2017], SLC6A6 [Ansar et al. 2020], PALLD (palladin, cytoskeletal associated protein) [Mastrototaro et al. 2023], SERINC2 [Hu et al. 2023], NMRAL1 [Gao et al. 2022], CTLA4 [Ruppert et al. 2010], CX3CR1 [Jeyalan et al. 2023], GDF15 [May et al. 2022], CXCL10 [Nogueira et al. 2012], BDNF (brain derived neurotrophic factor) [Costa et al. 2018], DIO2 [Wang et al. 2010], MYOZ1 [Arola et al. 2007], TP63 [Poloni et al. 2019], ACE (angiotensin I converting enzyme) [Küçükarabaci et al. 2008], PAK1 [Raut et al. 2015], GNB3 [Dewi et al. 2023], APOE (apolipoprotein E) [Jurkovicova et al. 2006], CACNG8 [Ortega et al. 2015], TNNT1 [Streff et al. 2019], HLA-DPA1 [Liu et al. 2006], IL34 [Xi et al. 2023], IER3 [Zhou et al. 2017], GZMB (granzyme B) [Zeglinski and Granville, 2020], CXCL9 [Nogueira et al. 2012], KCNN3 [Ortega et al. 2015], HAS3 [Wang et al. 2014] and TREH (trehalase) [Hieda et al. 2022] have been reported to be associated with dilated cardiomyopathy. VDR (vitamin D receptor) [Al-Nbaheen, 2023], PTX3 [Zanetti et al. 2014], FN1 [Page et al. 2022], APOB (apolipoprotein B) [Akdim et al. 2010], ADAMTS5 [Muneshige et al. 2011], CX3CR1 [Combadière et al. 2008], ABCG2 [Zhang et al. 2020], MDK (midkine) [Narita et al. 2008], ACE (angiotensin I converting enzyme) [Kondo et al. 2015], GNB3 [Suwazono et al. 2006], APOE (apolipoprotein E) [Sullivan et al. 1997] and CXCL3 [Martín-Fuentes et al. 2009] have been revealed to be associated with hypercholesterolemia. VSIG4 [Ding et al. 2023], FKBP5 [Wang et al. 2023], C3 [Dernellis et al. 2006], TIMP4 [Ye et al. 2021], SPP1 [Wang et al. 2021], BMP7 [Zhao et al. 2017], CFD (complement factor D) [Ding et al. 2023], AGTR1 [Feng et al. 2017], PTX3 [Bostan et al. 2020], FN1 [Zhu et al. 2022], HCN4 [Fraile et al. 2024], APOB (apolipoprotein B) [Zhong et al. 2022], BMAL1 [Wang et al. 2024], C1QC [Ding et al. 2023], FABP4 [López-Canoa et al. 2021], LOX (lysyl oxidase) [Adam et al. 2011], CD63 [Choudhury et al. 2007], NQO1 [Li et al. 2018], KCNH2 [Li et al. 2015], CORIN (corin, serine peptidase) [Chen et al. 2015], CDHR3 [Nakano et al. 2016], ELN (elastin) [Wang et al. 2021], RGS4 [Borges et al. 2023], GDF15 [Fragão-Marques et al. 2021], CXCL10 [Chen et al. 2021], BDNF (brain derived neurotrophic factor) [Pressler et al. 2024], ABCG2 [Wu et al. 2023, SMOC2 [Zhang et al. 2022], UCHL1 [Bi et al. 2020], ESR1 [Golubić et al. 2014], MYOZ1 [Martin et al. 2015], ACE (angiotensin I converting enzyme) [Gensini et al. 2003], PAK1 [DeSantiago et al. 2018], KCNA5 [Wang et al. 2022], GNB3 [Schreieck et al. 2004], APOE (apolipoprotein E) [Li et al. 2022], IL34 [Ma et al. 2023], KCNN3 [Rahm et al. 2021], KCNIP1 [Tsai et al. 2016] and FMOD (fibromodulin) [Liang et al. 2023] have been identified as a targets for atrial fibrillation. AGTR1 [Chou et al. 2022], PTX3 [Polat et al. 2015], LOX (lysyl oxidase) [Purushothaman et al. 2017], TGFB2 [Renard et al. 2013], ACE (angiotensin I converting enzyme) [Meurs et al. 2018] and GLIS1 [Yu et al. 2019] are correlates positively with the incidence of mitral valve disease. The results suggest that enriched genes might affect the disease progression of HCM and its associated complications include inflammation, heart failure, coronary artery disease, atherosclerosis, cardiac fibrosis, obesity, hypertension, diabetes mellitus, myocardial infarction, myocardial ischemia, cardiac death, cardiac arrest, dilated cardiomyopathy, hypercholesterolemia, atrial fibrillation and mitral valve disease. Novel biomarkers for HCM is listed in Table 11.

**Table 9.** Molecular docking interactions of screened phytoconstituents with ESR1 (PDB ID: 1A52)

| **Compound** | **ESR1 gene (pdb: 1A52) Amino acid interactions, Active Chain is B, reference ligand is Estradiol** |  |  |  |  |
| --- | --- | --- | --- | --- | --- |
|  | **Affinity**<br>**(kcal/mol)** | **Hydrogen bonding**<br>**interactions** | **pi-pi**<br>**interactions** | **Hydrophobic**<br>**interactions** | **Polar (electrostatic)**<br>**interactions** |
| Oleuropein | -7.745 | GLY217, ASP47,<br>LYS225, LEU221,<br>HIS220, ALA46, ARG90 | PHE100 | LEU83, LEU87,<br>ALA46 | -- |
| Puerarin | -6.559 | GLU49, THR43, LYS225 | LEU42 | LEU221, ALA46 | MET117, MET39,<br>ASP47 |
| Swertiamerin | -5.572 | GLU49, LEU83, ARG90,<br>PHE100 | -- | LEU80, ALA46,<br>LEU83 | MET39 |
| Mangiferin | -4.144 | MET117, THR43 | -- | LEU42, LEU221,<br>ALA46 | MET39 |
| Cocrystallised ligand | -10.754 | GLU49 | PHE100 | ILE120, MET84,<br>LEU87, LEU83,<br>ALA46 | -- |

**Table 10.** Predicted drug-likeness and ADMET properties of screened phytoconstituents.

| Parameter | Swertiamarin | Puerarin | Oleuropein | Mangiferin |
| --- | --- | --- | --- | --- |
| Molecular Weight (MW) | 374.12 | 416.11 | 540.18 | 422.34 |
| Topological Polar Surface Area (TPSA) | 155.14 | 160.82 | 201.67 | 201.12 |
| Hydrogen Bond Donors (HBD) | 5 | 6 | 6 | 8 |
| Hydrogen Bond Acceptors (HBA) | 10 | 9 | 13 | 11 |
| Rotatable Bonds | 4 | 3 | 11 | 2 |
| Lipinski | Accepted | Accepted | Rejected | Rejected |
| Log S (ESOL) | -2.013 | -3.924 | -1.443 | -3.514 |
| Log Po/w (XLOGP3) | -0.589 | 0.694 | 0.498 | 0.136 |
| Human Intestinal Absorption (HIA) | 0.926 | 0.841 | 0.908 | 0.956 |
| Caco-2 Permeability | -6.051 | -6.199 | -5.677 | -6.286 |
| BBB Permeant | 0.219 | 0.046 | 0.311 | 0.019 |
| P-gp Substrate | 0.075 | 0.967 | 0.99 | 0.988 |
| CYP2C9<br>Inhibitor | 0.004 | 0.056 | 0.061 | 0.022 |
| CYP3A4<br>Inhibitor | 0.016 | 0.068 | 0.198 | 0.02 |
| hERG Inhibitor | 0.083 | 0.041 | 0.097 | 0.105 |
| Carcinogenicity | 0.018 | 0.021 | 0.029 | 0.023 |

**Table 11.**
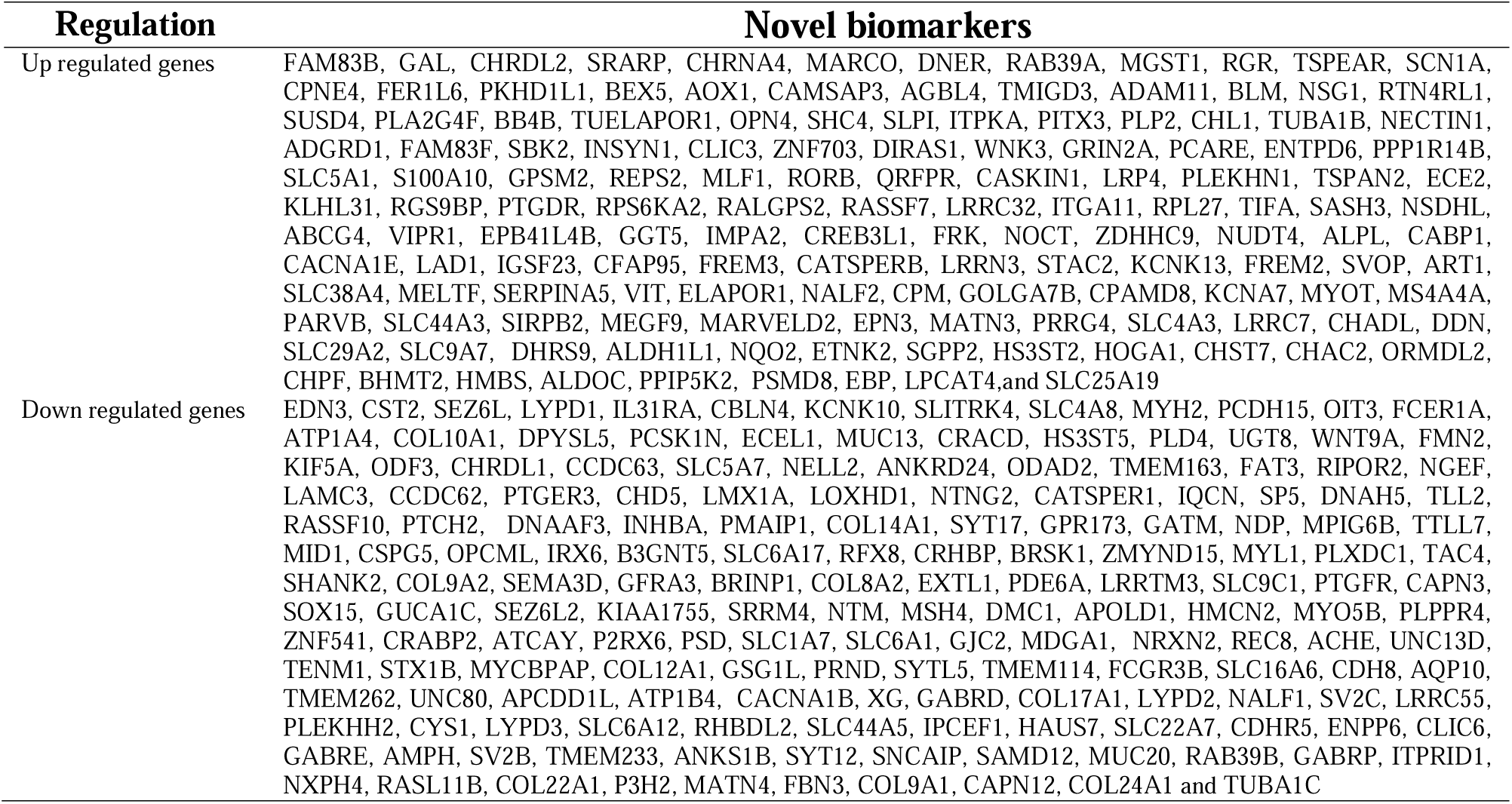
Novel biomarkers for HCM.

The PPI network was constructed using the HIPPIE database, and the analysis of significant modules was performed. Finally, we identified hub genes form PPI network and modules. FN1 [Page et al. 2022], ESR1 [Zhang et al. 2023], APLNR (apelin receptor) [Wang et al. 2020], LOX (lysyl oxidase) [Ma et al. 2011] and APOE (apolipoprotein E) [Anoop et al. 2010] can be used as an important therapeutic targets for coronary artery disease. FN1 [Page et al. 2022] and APOE (apolipoprotein E) [Sullivan et al. 1997] have become a potential biomarkers for hypercholesterolemia. Altered expression of FN1 [Yao et al. 2022], ESR1 [Puzianowska-Kuźnicka, 2012], PAK1 [Liu et al. 2020], LOX (lysyl oxidase) [González-Santamaría et al. 2016] and APOE (apolipoprotein E) [Shao et al. 2022] were associated with prognosis in myocardial infarction. FN1 [Zhang et al. 2022], SOX2 [Li et al. 2023], ESR1 [Kabir et al. 2015], PAK1 [Liu et al. 2020], APLNR (apelin receptor) [Zheng et al. 2023], LOX (lysyl oxidase) [Valls-Lacalle et al. 2021] and APOE (apolipoprotein E) [Shao et al. 2022] expression was shown to be regulated in myocardial ischemia. FN1 [Zhu et al. 2022], ESR1 [Golubić et al. 2014], PAK1 [DeSantiago et al. 2018], LOX (lysyl oxidase) [Adam et al. 2011] and APOE (apolipoprotein E) [Li et al. 2022] are implicated in atrial fibrillation. In many clinical studies, SOX2 [Njouendou et al. 2023], ESR1 [Zhang et al. 2023], SNCA (synuclein alpha) [Forloni, 2023], PAK1 [Chen et al. 2024], TGM2 [Chang et al. 2024], LOX (lysyl oxidase) [Huang et al. 2021], APOE (apolipoprotein E) [Christensen et al. 2011] and HLA-DPA1 [Lantermann et al. 2002] have been linked to the start of inflammation. Previous study confirmed that SOX2 [Zhao et al. 2018], PAK1 [Wang et al. 2014], LOX (lysyl oxidase) [Galán et al. 2017] and APOE (apolipoprotein E) [Qin et al. 2010] are strongly associated with HCM. The altered expression of SOX2 [Liang et al. 2021], PAK1 [Ai et al. 2011] and LOX (lysyl oxidase) [López et al. 2013] are related to prognosis in heart failure. SOX2 [Jiang et al. 2022], ESR1 [Zhou et al. 2022], APLNR (apelin receptor) [Stanchev et al. 2024], LOX (lysyl oxidase) [Vadasz et al. 2019] and APOE (apolipoprotein E) [de Leeuw et al. 2004] played an important role in hypertension. SOX2 [Gu et al. 2011], ESR1 [Dahlman et al. 2008], PAK1 [Veluthakal et al. 2018], TGM2 [Kolokotronis et al. 2024], APOE (apolipoprotein E) [El-Lebedy et al. 2018] and HLA-DPA1 [Varney et al. 2010] were involved in regulating the occurrence of diabetes mellitus. Previous studies have been reported that altered expression of ESR1 [Guclu-Geyik et al. 2020], APLNR (apelin receptor) [De Los Santos et al. 2023], LOX (lysyl oxidase) [Huang et al. 2021] and APOE (apolipoprotein E) [Farup et al. 2020] increased the risk of obesity. PAK1 [Singh et al. 2015], LOX (lysyl oxidase) [Ovchinnikova et al. 2014] and APOE (apolipoprotein E) [Marais, 2021] have been found to have a strong association with atherosclerosis. PAK1 [Raut et al. 2015], LOX (lysyl oxidase) [Sivakumar et al. 2008], APOE (apolipoprotein E) [Jurkovicova et al. 2006] and HLA-DPA1 [Liu et al. 2006] expression has been reported with dilated cardiomyopathy progression. TGM2 [Griffin et al. 2020], LOX (lysyl oxidase) [López et al. 2013] and APOE (apolipoprotein E) [Hada et al. 2022] are important in the successful progression of cardiac fibrosis. Altered expression and activity of LOX (lysyl oxidase) [Purushothaman et al. 2017] has been demonstrated in mitral valve disease. recent studies have proposed that APOE (apolipoprotein E) [Kida et al. 1995] is associated with the development and progression of cardiac arrest. Our studies have shown that novel biomarkers such as TUBA4A, RPS2 and LCK (LCK proto-oncogene, Src family tyrosine kinase) are involved in the regulation of HCM. We assumed that these hub genes might play a essential role in the pathogenesis of HCM; however, this hypothesis needs to be verified in future investigation.

It is evident that miRNA-hub gene regulatory network and TF-hub gene regulatory network play a key part in the development of HCM. In this way, it increases the knowledge of HCM identification and contributes to targeted therapeutic management strategies and HCM prediction. FN1 [Page et al. 2022], FKBP5 [Wang et al. 2020], ESR1 [Zhang et al. 2023], hsa-miR-34a-5p [Yalım et al. 2023], BRCA1 [Powell et al. 2018], STAT1 [Zhao et al. 2024], STAT3 [Ouyang et al. 2022], MEF2A [Muhammad Sulaiman et al. 2024] and GATA2 [Luo et al. 2021] participates in the regulation of the pathophysiological process in coronary artery disease. Studies have demonstrated that the altered expression of FN1 [Page et al. 2022] in patients with hypercholesterolemia. Recent studies have demonstrated that the FN1 [Yao et al. 2022], ESR1 [Puzianowska-Kuźnicka, 2012], PAK1 [Liu et al. 2020], hsa-miR-34a-5p [Wang and Cao, 2020], PRRX2 [Bai et al. 2020], BRCA1 [Shukla et al. 2011], STAT1 [Zhang et al. 2023], STAT3 [Bao et al. 2024], MEF2A [Guella et al. 2009] and GATA2 [Izadpanah et al. 2021] are associated with myocardial infarction. FN1 [Zhang et al. 2022], FHL2 [Goltz et al. 2016], ESR1 [Kabir et al. 2015], PAK1 [Liu et al. 2020], SRY (sex determining region Y) [Liu et al. 2019], STAT1 [Zhou et al. 2019] and STAT3 [Mahdiani et al. 2022] are important in the development of myocardial ischemia. Recent studies have proposed that the FN1 [Zhu et al. 2022], FKBP5 [Wang et al. 2023], ESR1 [Golubić et al. 2014], PAK1 [DeSantiago et al. 2018], STAT1 [Tsai et al. 2008] and STAT3 [Hsu et al. 2019] are associated with in atrial fibrillation. TFRC (transferrin receptor) [Zhou et al. 2024], PAK1 [Ai et al. 2011], BRCA1 [Wang et al. 2024], STAT1 [Shi et al. 2024] and STAT3 [Zhuang et al. 2022] are associated with progression of heart failure. TFRC (transferrin receptor) [Qiu et al. 2024], FKBP5 [Willmer et al. 2022], FHL2 [Sommer et al. 2023], RPLP0 [Pérez-Gómez et al. 2023], ESR1 [Guclu-Geyik et al. 2020], BRCA1 [Bhardwaj et al. 2023], STAT1 [Li et al. 2023], STAT3 [Su et al. 2020] and GATA2 [Menghini et al. 2005] are involved in the progression of obesity. The abnormal expression of TFRC (transferrin receptor) [Yang et al. 2024], FHL2 [Li et al. 2019], TPM3 [Xu et al. 2021], ESR1 [Zhou et al. 2022], CHRM3 [Cowley et al. 2018], BRCA1 [Chida-Nagai et al. 2019], STAT1 [Jandl et al. 2022], STAT3 [Bisserier et al. 2021], FOS (Fos proto-oncogene, AP-1 transcription factor subunit) [Altura et al. 2003] and GATA2 [Zhang et al. 2022] have been reported in hypertension. TFRC (transferrin receptor) [Xu et al. 2015], FKBP5 [Wada et al. 2021], FHL2 [Arimura et al. 2007], PAK1 [Raut et al. 2015], TP63 [Poloni et al. 2019], hsa-miR-34a-5p [Wu et al. 2021], hsa-miR-194-5p [Wang et al. 2024], BRCA1 [Nozynski et al. 2016], STAT3 [Li et al. 2019] and MEF2A [Qiao et al. 2020] are the risk factors of dilated cardiomyopathy. FKBP5 [Liu et al. 2023], FHL2 [van de Pol et al. 2020], TPM3 [Mansfield et al. 2016]. ESR1 [Zhang et al. 2023], PAK1 [Chen et al. 2024], SNCA (synuclein alpha) [Forloni, 2023], hsa-mir-374a-5p [Perez-Sanchez et al. 2022], hsa-miR-34a-5p [Mirra et al. 2022], BRCA1 [Bruand et al. 2021], STAT1 [Ploeger et al. 2022], STAT3 [Hu et al. 2019], FOS (Fos proto-oncogene, AP-1 transcription factor subunit) [Catar et al. 2013], MEF2A [Cilenti et al. 2021] and GATA2 [Baba et al. 2024] might be involved in the process of inflammation. Studies have also confirmed that FKBP5 [Sidibeh et al. 2018], FHL2 [Habibe et al. 2022], ESR1 [Dahlman et al. 2008], CHRM3 [Guo et al. 2006], PAK1 [Veluthakal et al. 2018], STAT1 [Cao et al. 2022] and STAT3 [Zhang et al. 2022] participates in diabetes mellitus. FHL2 [Friedrich et al. 2014], PAK1 [Wang et al. 2014, STAT1 [Shi et al. 2024], STAT3 [Wang et al. 2018] and MEF2A [Konno et al. 2010] have been reported to be correlated with HCM. FHL2 [Chu et al. 2010], PAK1 [Singh et al. 2015], hsa-miR-34a-5p [Yalım et al. 2023], hsa-miR-524-5p [Li et al. 2018], SRY (sex determining region Y) [Cai et al. 2015], BRCA1 [van Bommel et al. 2021], STAT1 [Zhang et al. 2022], STAT3 [Yang et al. 2024], MEF2A [Zhou et al. 2015] and GATA2 [Yin et al. 2020] have been reported to be associated with atherosclerosis. FHL2 [Anwaier et al. 2022], PRRX2 [Bai et al. 2020], BRCA1 [Yao et al. 2013], STAT1 [Dai et al. 2013], STAT3 [Jiang et al. 2024], MEF2A [Chen et al. 2016] and GATA2 [Wang et al. 2022] are a pivotal regulator of cardiac fibrosis. STAT3 [Huang et al. 2015] is associated with the progression of cardiac arrest. MEF2A [Naya et al. 2002] serves a vital role in cardiac death. SH3KBP1, TMEM30B, PTPRR (protein tyrosine phosphatase receptor type R), hsa-miR-520d-5p, hsa-mir-4646-5p, hsa-mir-548aw, hsa-miR-8052, hsa-mir-302b-3p, hsa-mir-548i, USF2 and EN1 might be a novel biomarkers associated with HCM and might be involved in the development of HCM. The results therefore were in accordnce with previous studies and suggested that the hub genes, miRNAs and TFs served significant roles in the development of HCM and its associated complications.

For further investigating the action of drug on hub gene targets in ARDS. Our findings suggest that drugs- Succinylcholine, Memantine, Umeclidinium, Bromazepam target hub genes include CHRNA4, GRIN2A, CHRM3 and GABRE, potentially controlling the development of HCM.

Hypertrophic cardiomyopathy (HCM) is a multifactorial myocardial condition in which pathological hypertrophy, increasing interstitial fibrosis, and poor cardioprotective signaling work together to cause abnormal cardiac remodeling. Therapeutic techniques that address both fibrotic remodeling and the loss of protective molecular pathways are therefore particularly important. In this study, an integrated in-silico framework was used to test four bioactive phytoconstituents Swertiamarin, Puerarin, Oleuropein, and Mangiferin against two biologically different but complimentary targets, LGALS3 and ESR1, found from HCM transcriptome analysis.

LGALS3 (galectin-3) upregulation in HCM indicates its critical role in fibroblast activation, collagen deposition, and inflammatory signaling, all of which contribute to myocardial stiffness and diastolic dysfunction. Docking study revealed that all tested phytoconstituents effectively occupied LGALS3’s carbohydrate recognition domain, engaging conserved residues such as TRP69, ARG50, and HIS46, which are required for galectin-3 ligand recognition and downstream signalling. Notably, Puerarin and Oleuropein had binding affinities comparable to or greater than those of the co-crystallized reference ligand, indicating a high potential for competitive interference with galectin-3-mediated fibrotic signaling. These chemicals’ capacity to form a dense hydrogen-bonding network with π-π interactions within the LGALS3 binding pocket suggests a persistent and physiologically relevant inhibitory binding mechanism. From a disease standpoint, such inhibition may translate into attenuation of extracellular matrix buildup and suppression of maladaptive cardiac remodeling typical of HCM.

In parallel, transcriptome study indicated ESR1 downregulation, which is consistent with reduced estrogen-mediated cardioprotective signaling in HCM. Estrogen receptor-α inhibits pathological hypertrophy via regulating calcium levels, oxidative stress, and inflammation in cardiomyocytes. Docking against the agonist-bound ESR1 structure (PDB ID: 1A52) revealed that phytoconstituents may be stably accommodated within the estradiol-binding cavity, generating interactions with residues implicated in receptor stability and activation. Although their binding affinities were lower than those of estradiol, the observed interaction patterns point to a modulatory rather than antagonistic role, which could be especially useful in restoring basal cardioprotective signaling without generating excessive hormonal activation. Oleuropein had the most favorable interaction profile with ESR1, indicating that it could operate as a partial agonist or stabilizer of receptor activation in the context of ESR1 downregulation.

The current study examines the convergence of binding efficacy and pharmacokinetic feasibility. Despite the structural complexity associated with phytochemicals, ADMET analysis found that all screened compounds have adequate solubility and high projected intestine absorption, indicating their appropriateness for oral administration. Importantly, the consistently low anticipated hERG inhibition across all drugs addresses a crucial safety concern in HCM, where arrhythmogenic risk is naturally high. The lack of considerable CYP2C9 and CYP3A4 inhibition suggests a minimal risk of harmful drug-drug interactions, which is critical for patients who frequently get long-term treatment. Furthermore, the minimal BBB permeability seen across all drugs suggests target specificity for peripheral cardiac tissues, reducing undesired central nervous system effects.

The integration of docking and ADMET properties with *in silico* drug development promotes a multi-target treatment approach in HCM. While LGALS3 suppression may reduce fibrosis-induced structural remodeling, ESR1 modulation provides a parallel pathway for restoring endogenous cardioprotective signaling. Within this paradigm, Puerarin and Oleuropein appear as leading contenders, combining significant target engagement with favorable safety and pharmacokinetic profiles, while Swertiamarin and Mangiferin provide supportive interaction patterns that may contribute to synergistic cardioprotective benefits.

In conclusion, via bioinformatics analysis of NGS data from HCM human models, we identified and selected several essential DEGs that are associated in the development of HCM and its associated complication. Furthermore, our results indicate that the identified DEGs not only are linked with numerious GO terms and functions, and signaling pathways, but also could construct complex PPI, miRNA-hub gene regulatory network and TF-hub gene regulatory network. In addition, a lot of DEGs with distinct GO functions and signaling pathways could form significant modules. Our investigations provides novel insight on HCM advancement not only in the gene level, but also in the GO functions, signaling pathways and module levels. The identified DEGs, associated GO functions, signaling pathways, and modules could serve as potentially diagnostic and therapeutic targets for HCM. In addition, our investigations has some limitations. For example, the identified DEGs, associated GO functions, signaling pathways, and modules were only generated from bioinformatics analysis. These analysed data still need to be further confirmed in clinical specimens in future investigations. Despite these limitations, the results of our investigations for further examining the roles and associated molecular mechanisms mediated by identified key genes, associated GO functions, signaling pathways, and modules in HCM.

Our study used an integrated *insilico* strategy to identify phytoconstituents with possible significance to hypertrophic cardiomyopathy by focusing on both upregulated LGALS3 and downregulated ESR1. The docking results showed stable and positive interactions of the screened phytochemicals within functionally relevant binding pockets of both targets, whereas ADMET analysis revealed acceptable pharmacokinetic features and low predicted cardiotoxicity. These findings support the potential of the selected phytoconstituents as multi-target modulators capable of reducing fibrosis-related remodeling and restoring cardioprotective signaling in hypertrophic cardiomyopathy.

Despite these encouraging findings, the research is hampered by its dependence on computational projections. Molecular docking and ADMET analyses do not adequately account for dynamic protein behavior, downstream signaling consequences, or *invivo* pharmacokinetics. As a result, experimental validation with biochemical tests, cellular hypertrophy models, and animal investigations is required to confirm the therapeutic utility of the identified lead candidates.

## Acknowledgement

I thanks very much to Daniel Bernstein, Stanford University School of Medicine, Stanford, California, USA, the author who deposited their NGS dataset GSE180313, into the public GEO database.

## Conflict of interest

The authors declare that they have no conflict of interest.

## Ethical approval

This article does not contain any studies with human participants or animals performed by any of the authors.

## Informed consent

No informed consent because this study does not contain human or animals participants.

## Availability of data and materials

The datasets supporting the conclusions of this article are available in the GEO (Gene Expression Omnibus) (https://www.ncbi.nlm.nih.gov/geo/) repository. [(GSE180313) https://www.ncbi.nlm.nih.gov/geo/query/acc.cgi?acc=GSE180313]

## Consent for publication

Not applicable.

## Competing interests

The authors declare that they have no competing interests.

## Funding

The authors received no financial support for the research

## Author Contributions

B. V. - Writing original draft, and review and editing

S.P. - Formal analysis and validation

C. V. - Software and investigation

## Notes

### Competing Interest Statement

The authors have declared no competing interest.

### Summary of Updates

we included drug-hug gene interaction network for prediction of drug molecules, Molecular docking studies and ADMET studies for screening of drug molecules

